# TANGO-Light - optogenetic control of transcriptional modulators

**DOI:** 10.1101/2023.05.31.543150

**Authors:** Alicja Przybyszewska-Podstawka, Joanna Kałafut, Jakub Czapiński, Thu Ha Ngo, Arkadiusz Czerwonka, Adolfo Rivero-Müller

## Abstract

Cell signalling pathways, in particular downstream receptor activation, frequently converge in the activation of transcriptional modulators. Yet, cells are able to differentiate the stimulation of each receptor. It has become clear that transcriptional modulators, such as transcription factors, do not work in on or off states but rather in patterns of active/inactivate conformations. Thus, it is the intensity and duration of such fluctuating activity that result in differential cellular and genes expression changes, and this is challenging to replicate using traditional methods such as inhibitors or genetic constructs. Optogenetics, which is based on the use of light-responsive proteins, offers precise control over biological processes, in a spatio-temporal manner, allowing targeted fine-tuned modulation of specific proteins or signalling pathways. Here, we engineered an optogenetic system to control transcriptional modulators, by fusing a photoactivatable receptor and the TANGO system. By this mean we show that we are able to control a plethora of transcriptional modulators by light. And by doing so, changing cells fate - inducing cells to acquire a more mesenchymal or epithelial phenotype. This optogenetic system was also adapted to mimic signalling pathways such as Notch and Wnt in a light-dose dependent manner. Finally, we show that this light-responsive system can be induced by natural light sources upon cell-cell contact/proximity.

## INTRODUCTION

In response to environmental stimulations or developmental cues, the activity of transcriptional modulators varies over time, and these fluctuations are essential for regulating downstream gene expression^1^. The kinetics of transcriptional modulators depend on many factors, including concentration, affinity for the DNA binding site, presence of co-activators, co-repressors, and cellular context^2, 3^. Recent findings suggest that dynamic changes in transcriptional modulators (TMs) activity are essential for regulating gene expression patterns and cell fate decisions^4, 5^. For example, prolonged or transient transcription factor (TF) activity can lead to different gene expression profiles, while oscillatory TF activity regulate the timing and coordination of gene expression during development^6^.

Our understanding of how transcriptional modulators function has only recently started to be unraveled, thanks to new techniques that allow observing and imaging the dynamic behavior of individual TMs in real-time within living cells.

The phenotype of a cell is a consequence of the cell’s signalling status, which in turn of mostly controlled by the crosstalk and activity of transcriptional modulators^7–9^. The expression and activity of these key regulatory molecules is thus tightly regulated^10–13^. In order to study and control cell signalling, and thus fate, there is a need to develop tools to control single or orthogonal pathways with spatio-temporal precision. Likely, the most precise systems to-date are based on optogenetics due to their quick response to light stimulation,^14^ non-invasive nature, reversibility and inexpensive modulation, yet mostly due to the spatio-temporal control of the input (light) down to single cells or area of a cell^15, 16^. Various optogenetic tools have been recently developed to control signalling pathways^17, 18^, gene expression^19, 20^ and DNA recombination^21^, although in most cases these are engineered in case to case basis what hinders the exchanging of components easily. For example, previous light inducible systems involving membrane receptors, such as iTango^22^ and SPARK2^23^, utilize different GPCRs where a fused transcriptional activator (usually tTA) is protected from proteolytical cleavage by the LOV (Light-Oxygen Voltage) domain. While these methods are excellent to study or control individual GPCRs, ligand activation of these receptors is still required^24, 25^. To engineer a universal system that releases transcriptional modulators upon illumination, we engineered a synthetic system where the photoactivatable-CXCR4 receptor (PA-CXCR4), a rhodopsin-chemokine receptor chimera^26^, was fused to the TANGO system^27^ to regulate transcriptional modulators – we named it Tango-Light (TL for short).

TL was applied to control a variety of transcriptional modulators that result in specific signal-dependent gene expression and phenotypical changes such as epithelial to mesenchymal transition (EMT) and its reverse process (MET), cell proliferation, migration and invasiveness, or to control Notch, or Wnt signalling in a spatio-temporal manner. Moreover, we show that TL can be also activated by cell-cell contact by presentation of a natural light source.

## RESULTS

### Optogenetic control of transcriptional modulators

To engineer the photo-activatable release of transcriptional modulators (TMs) from the plasma membrane, we generated a chimera between the photoactivatable-CXCR4 receptor (PA-CXCR4)^28^ and the TANGO system^27^. The TANGO system consists of any G protein-coupled receptor (GPCR) fused with the C-terminus of the vasopressin receptor (V2 tail), containing a TEV cleavage site (TEVcs) followed by the tetracycline transactivator (tTA)^28^. Upon activation of the GPCR, β-arrestin-TEV is recruited, resulting in the proteolytical release of tTA from the plasma membrane. By fusing the PA-CXCR4 receptor to the C-terminal to that of TANGO (V2 tail-TEV site-tTA) (TANGO-light-tTA, or TL-tTA for short), we expected that upon blue light illumination it will recruit β-arrestin-TEV follow by the release the tTA from the membrane so it then translocate to the nucleus (**Fig. 1A**).

**Figure 1.**
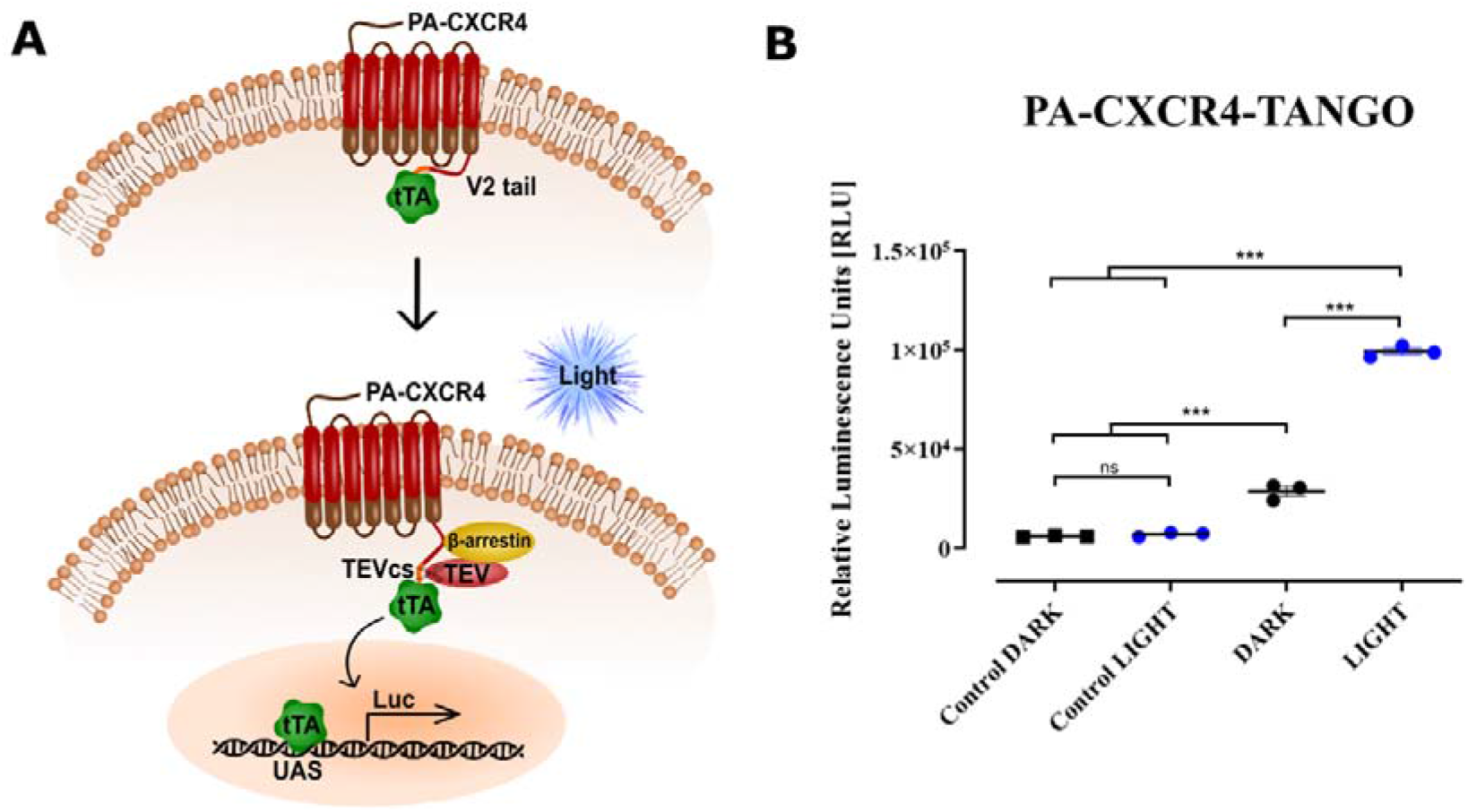
TANGO-Light with tTA (TL-tTA) system. (**A**) Diagram depicting the structure and function of a light activated system. Fusing PA-CXCR4 receptor with TANGO results in a light activatable receptor that, upon stimulation, recruits β-arrestin-TEV who cleaves and releases tTA from the membrane to translocate to the nucleus. (**B)** Relative luminescence level in HTLA cells after photoactivation of TL-tTA or kept in dark. Mock-transfected cells as controls. The results analyzed with one-way ANOVA test and Tukey’s *post-hoc* test vs. after light activation with pulses in a pattern of activation 0.05s/5s break (LIGHT) and non-stimulated (DARK). *p□≤□0.05; **p□≤□0.01; ***p□≤□0.001 were considered statistically significant, ns – non significant.

As a proof-of-concept, HTLA^28^ cells, stably transfected with a tTA-dependent Firefly luciferase (pTET-FFluc) reporter and β-arrestin2-TEV, were transfected with TL-tTA. 24h after transfection, these cells were either kept in the dark (DARK) or photostimulated (LIGHT) using pulses of light 0.05s/5s break. Control cells, mock transfected, were also irradiated (LIGHT) or not (DARK) (**Fig. 1B)**. As expected, light activation resulted in statistical significant increase of reporter activity, while mock transfected cells showed baseline reporter activity whether in the dark or after light illumination (**Fig 1B**).

### Optogenetic induction of EMT

Knowing that TL activation resulted in the release of tTA, as determined by downstream gene activation, we then exchanged tTA by a factor that is known to induce epithelial to mesenchymal transition (EMT). In cancer cells, multiple transcription factors have been shown to promote tumor progression, invasion and drug resistance by inducing EMT^29, 30^. Among these factors is SNAI1, whose overexpression increases cell invasiveness, angiogenesis, immune evasion^31, 32^, and chemoresistance^12^.

Thus, we hypothesized that blue light activation of TL-SNAI1 would allow for the control of its downstream target genes and EMT induction (**Fig. 2A**).

**Figure 2.**
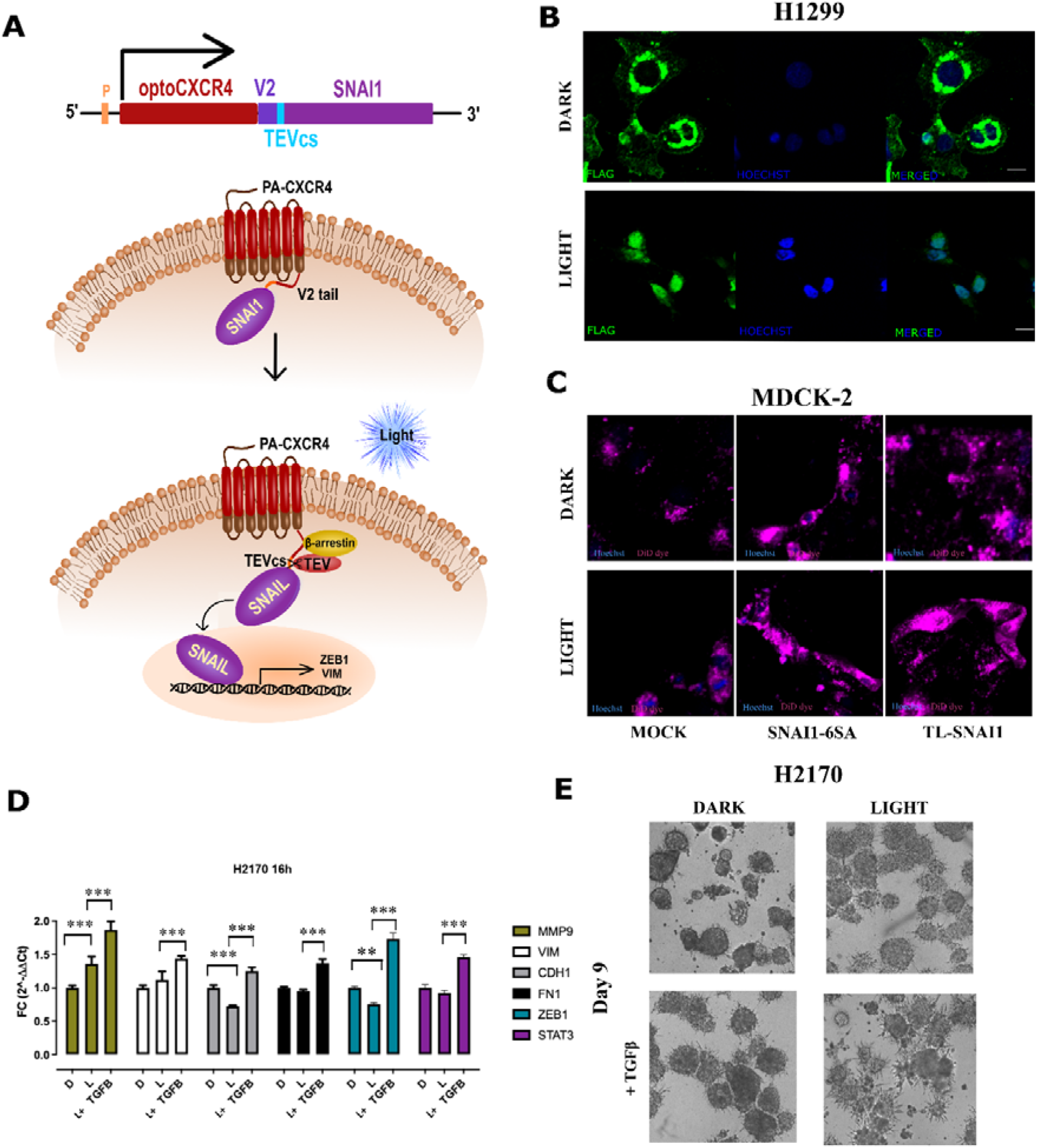
Optogenetic induction of EMT by TL-SNAI1. (**A**) Diagram showing the architecture of the TL-SNAI1 system. (**B**) Immunocytochemical staining of SNAI1 (green) before (DARK) and after photoactivation pattern of 0.05s activation and 5 s brake (LIGHT), showing the relocalisation of SNAI1 from the membrane to the nucleus. Hoechst 33,342 (blue) was used to stain nuclei, (**C**) MDCK-2 cells stained with Vybrant™ DiD (purple) dye and Hoechst 33,342 (blue) before and after blue illumination: TL-SNAI1, mock transfected, as a negative control, and transfected with SNAI1 as a positive control. **(D)** The mRNA expression of *MMP9*, *VIM*, *CDH1*, *FN1*, *ZEB1*, and *STAT3* genes was determined by qPCR (2–ΔΔCt) 16 after light stimulation of TL-SNAI1 H2170. Some of the light-activated cells were additionally cultured with TGFβ in medium. The results are presented as fold change (FC) values (mean□±□SD) normalized to the *GAPDH* gene expression. All data were analyzed with a one-way ANOVA test and Tukey’s multiple comparisons post-hoc test vs. not light stimulated cells (the time point 0 h and D; dark, respectively). **(E)** TL-SNAI1 H2170 organoids were cultured between two layers of Matrigel (lower gel − 4 mg/ml and upper gel − 2 mg/ml) with TGFβ-supplemented in the gel and medium of the top of spheroids and controls without this EMT inducer. Spheroids were imaged for 9 days. Cells were light-activated using short pulses for 3 h every day (LIGHT). Images represents comparison of the morphology of spheroids formed by TL-SNAI1 H2170 photo-stimulated (LIGHT) or kept in the dark (DARK).

To validate that SNAI1 was indeed released from the cell plasma membrane and translocated to the nucleus, TL-SNAI1-expressing H1299 cells were fixed before and after light activation, and immunofluorescence staining of SNAI1 was performed. As expected, SNAI1 was found at the plasma membrane prior to photostimulation, and in the nucleus after (**Fig. 2B**).

To prove that we could photo-induce EMT using TL-SNAI1, we picked a highly epithelial cell line (MDCK-2) to visualize EMT as this conversion results in clear phenotypical changes from “square” shape to “spindle” shape (fibroblastoid)^33^. To assist with the delineation of cell shape, cells were stained with Vybrant™ DiD dye. Light activated TL-SNAI1 MDCK-2 cells became more mesenchymal, acquiring a fibroblastoid (spindle) shape and losing of cell-cell contacts (**Fig. 2C, right panel)** in comparison to mock transfected cells, whether illuminated or not, which grew in tight clusters until become a monolayer, which is characteristic of these cells (**Fig. 2C, left panel**). MDCK-2 cells transfected with a plasmid over-expressing SNAI1 were used as control and these cells also became fibroblastoid-like (**Fig. 2C middle panels**). To confirm that TL-SNAI1 retains the gene targeting functions of SNAI1, we analysed the response of downstream target genes by qPCR after light activation. Genes that are dependent on SNAI1 activity, such as *MMP9, VIM, CDH1, FN1, ZEB1,* and *STAT3* were selected for analysis^34^. The level of expression of selected genes was analysed 16 hours after light activation. Some of the light-activated cells were additionally cultured with TGFβ in medium. All analysed genes showed statistically significant higher expression in optogenetically stimulated cells vs control cells (kept in dark) **(Fig. 2D)**.

Since SNAI1 signalling has been well characterized in lung cancer^35, 36^, H2170 lung cancer cells were selected for further functional analyses using the TL-SNAI1 system. To confirm the effect of the TL-SNAI1 system on the development of cancer cells under conditions that mimic physiology, cells were cultured in 3D spheroids using Matrigel supplemented with TGFβ, the latter to enhance the response as single transcriptional activators require long periods of induction before causing EMT^37, 38^. In TL-SNAI1-H2170 cells, light activation resulted in a noticeably increased growth of spheroids, and morphological changes, including a more migratory phenotype, especially when culturing with EMT enhancer TGFβ, which boosted these phenotypical changes **(Fig. 2E)**.

### TL-Elf3 and SOX-2 as a repressors of EMT

Knowing we can induce EMT, we decided to analyse whether we could apply TL to do the opposite. It is widely known that Elf3 can act as a repressor of EMT-associated genes in lung cancer cell lines, leading to the inhibition of EMT, decreased cell migration, and invasion^39^. Similarly, overexpression of SOX2 may inhibit the EMT process in lung cancer, resulting in decreased cell migration and invasion. SOX2 is known to upregulate the expression of Fibronectin (*FN1*) ^40^ among other epithelial genes.

To prove this, we checked the expression levels of EMT genes in embryonic kidney HEK293T cells transfected with TL-Elf3 and TL-SOX2.To demonstrate the effect of the system with transcriptional repressors, we analyzed the response of Elf3 and SOX-2 target genes by qPCR, before and after blue light activation. The level of expression of selected genes was analysed 16 hours after light activation. Blue light activation of TL-Elf3 led to increased expression of *MMP9* and *SNAI1* in HEK293T cells (**Fig. 3A**). In the case of light activation of TL-SOX2, an increase in the expression of *CDH1* and *FN1* was observed, with a simultaneous decrease in the expression of *MMP9*.

**Figure. 3.**
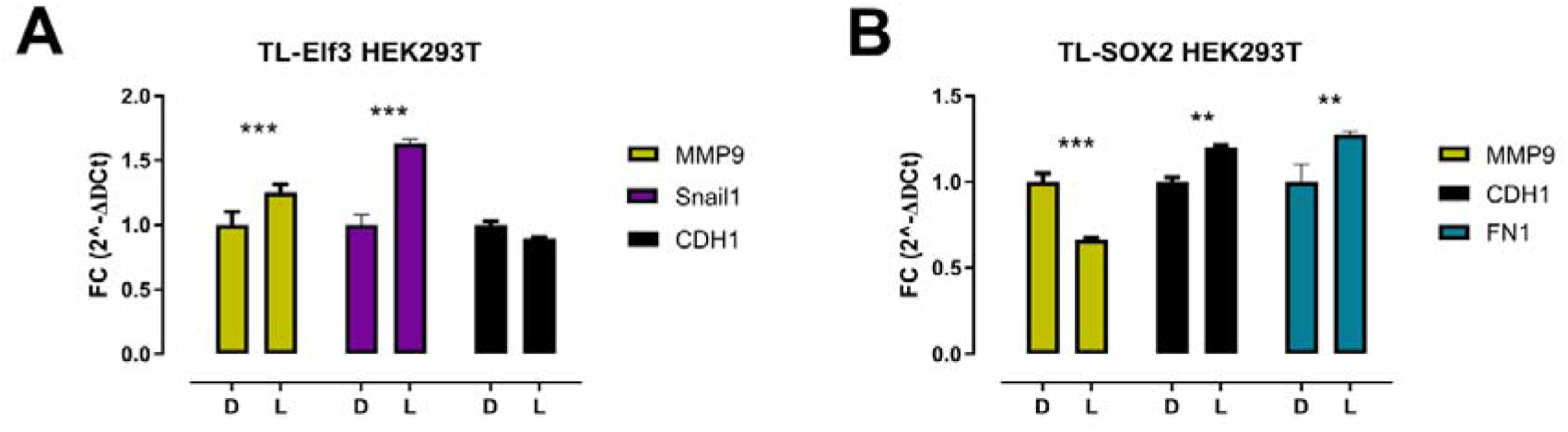
Optogenetic control of Elf3 and SOX2 with TL. Analysis of target genes downstream of (**A**) TL-Elf 3 and (**B**) TL-SOX-2. We analyzed the expression of these genes by qPCR in cell keep in the dark (D) and those light stimulated (L). The results analyzed with one-way ANOVA test and Tukey’s *post-hoc* test vs. after light activation LIGHT and without DARK. *p□≤□0.05; **p□≤□0.01; ***p□≤□0.001 were considered statistically significant, ns – non significant.

### Optogenetic modulation Notch1 activity

We then tested whether the conceptual design of the TL assay could be applied to different transcriptional modulators, for which we selected the intracellular domain of NOTCH-1 (N1ICD), as NOTCH1 is known to promote tumor growth and metastasis by enhancing the angiogenic and invasive potential of breast and lung cancer cells, as well as contributing to chemoresistance^41, 42^. Moreover, we and others have previous reported an optogenetic-controlling system for Notch signalling^43, 44^.

N1ICD was anchored to the C-terminal of TL, imitating the inactive endogenous Notch receptor. Just like the endogenous NOTCH, where mechano-activation results in the release of N1ICD and translocation to the cell’s nucleus where it acts as a transcriptional activator for target genes, photoactivation of TL-N1ICD should also result in release and translocation of N1ICD (**Fig. 4A**).

**Figure 4.**
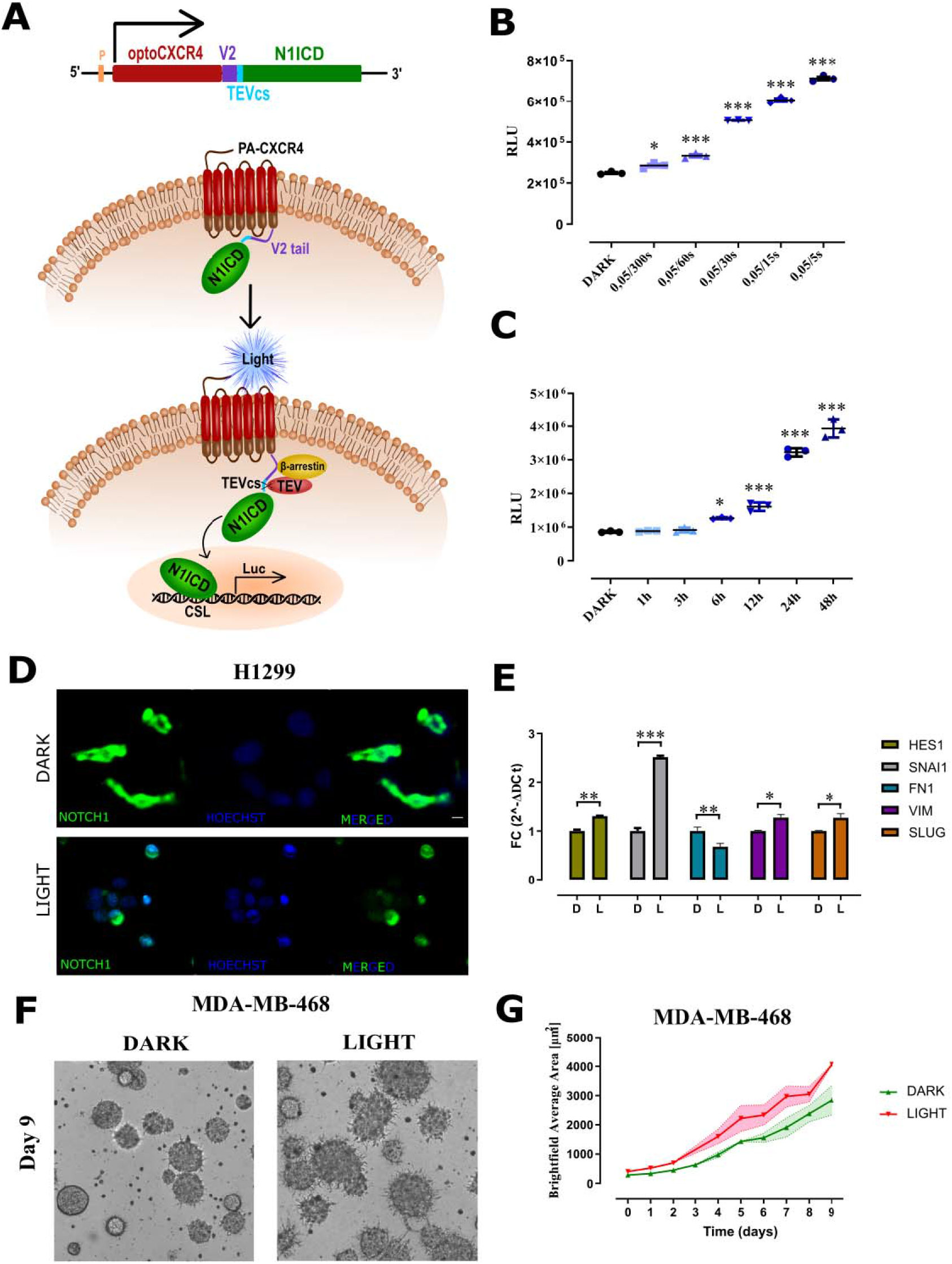
Optogenetic control of Notch1 signalling. **(A)** Diagram showing the structure and action of a light activated system. **(B)** Light activation of TL-N1ICD, resulted in a dose-dependent increase of FFluc expression as compared to those kept in the dark. **(C)** In order to select the optimal length of irradiation, cells were pulse-irradiated for 1, 3, 6, 12, 24 and 48 h. Cells were photoactivated 24 h after transfection and lysed 48 h after the start of irradiation to measure FFluc activity. The reporter’s response was significantly higher for longer exposure times. **(D)** Immunocytochemical staining of N1ICD (green) before (DARK) and after photoactivation (LIGHT), showing the relocalisation of NOTCH1 from the membrane to the nucleus. Hoechst 33,342 was used to stain nuclei (blue). **(E)** The mRNA expression of Notch1-target genes: *HES1*, *SNAI1*, *FN1 (FN1), VIMENTIN (VIM) and SLUG* was determined by qPCR (2–ΔΔCt) 24 after light stimulation of TL-N1CD HEK293T cells. The results are presented as fold change (FC) values (mean□±□SD) normalized to the *GAPDH* gene expression. **(F)** 3D organoid images of MDA-MB-468 cells in 9th day not photostimulated (DARK) and photostimulated (LIGHT). Light stimulation resulted in a noticeable increase in the growth of the spheroids, and morphological changes. **(G)** Comparison of the growth MDA-MB-468 organoids in dark and light-activating condition for 9 days. All data were analyzed with a one-way ANOVA test and Tukey’s multiple comparisons post-hoc test vs. not light stimulated cells (the time point 0 h and D; dark, respectively). *p□ ≤ □0.05; **p□ ≤ □0.01; ***p□ ≤ □0.001 were considered statistically significant, ns – non significant.

HEK293T cells stably carrying the Notch reporter, 12xCSL-Firefly luciferase (12xCSL-FFluc), were transfected with the TL-N1ICD and β-arrestin-TEV constructs. Transfected cells were then exposed to blue light (456 nm) in different patterns of light activation (pulses) or kept in the dark. We first tested which light/dark pattern would be the most optimal for the signal intensity. For this purpose, the cells were subjected to 0.05 s pulses of light at various time intervals (300, 60, 30, 15 and 5 s) within 1 h, which corresponds to 12, 60, 120, 240 and 720 pulses of blue light, respectively. Untransfected cells were also irradiated to eliminate other effects of light, these cells only showed baseline luminescence values (not shown). Samples were collected to measure the relative FFluc activity 24 h after activation. As expected, the greater the number of light pulses, the higher reporter’s response. Light activation of TL-N1ICD, resulted in a statistical significant increase of FFluc expression as compared to those kept in the dark, or mock transfected exposed to light, validating the specificity of signalling (**Fig. 4B**). A pattern of 0.05s blue light pulses with a 5s interval which resulted in the highest reporter response was therefore selected for further analyses.

In order to select the optimal length of irradiation, cells were then activated for 1, 3, 6, 12, 24 and 48 h. All cells were photoactivated 24 h after transfection and lysed 48 h after the start of irradiation. Again, the reporter’s response was significantly higher for longer exposure times (**Fig. 4C**). For subsequent analyses, the overnight activation pattern of 0.05s/5s was used.

Immunofluorescence assays were performed before and after light activation to determine the cellular localization of the N1ICD. As expected, N1ICD was at the plasma membrane prior to activation, from which it was released upon blue light activation, and translocated to the nucleus (**Fig. 4D**). To confirm that TL-N1ICD retains the gene targeting functions of endogenous N1ICD, we analysed the response of Notch target genes by qPCR after light activation. Genes that are dependent on NOTCH1 activity, such as *HES1*, *SNAI1*, *FN1, VIMENTIN* and *SLUG*^45–47^ were selected for analysis. The level of expression of selected genes was analysed 16 hours after light activation. All analysed genes showed statistically significant higher expression compared to control cells (**Fig. 4E**).

The role of Notch1 signalling as a tumour promotor has been well characterized in breast cancer, therefore the MDA-MB-468 breast cancer cells were selected to further test the TL-N1ICD system. To confirm the effect of the TL-N1ICD system on the development of cancer cells under physiological mimicking conditions, cells were cultured in 3D organoids using Matrigel. In MDA-MB-468 cells, light activation resulted in a noticeably increased growth of the organoids, as well as a more aggressive phenotype (**Fig. 4F and G**).

### TL- **β** -catenin

We then decided to focus in other signalling pathways, and selected WNT signalling. For this we selected β-catenin, a key transcriptional co-activator that is regulated by degradation by a protein complex called the destruction complex, which consists of several proteins including adenomatous polyposis coli (APC), Axin, casein kinase 1 (CK1), and glycogen synthase kinase 3β (GSK-3β). Within the destruction complex, GSK-3β phosphorylates the N-terminal β-catenin, primarily at serine and threonine residues, which are recognized by a component of the E3 ubiquitin ligase complex, called β-TrCP (β-transducin repeat-containing protein), target the protein to subsequent degradation by the 26S proteasome^48–50^. In order to bypass the mechanism of β-catenin degradation in the cell, we used the N90-terminal truncated^51, 52^ β-catenin. Thus, we fused this constitutively active β-catenin mutant^53^ to the TL system so we can modulate its activity by light.This allows studying the direct control of prolonged β-catenin (βCat) activity on gene expression and phenotypical manifestations.

Thus, activation of TL-βCat by blue light shall result in the activation of downstream βCat-target genes (**Fig. 5A**). In order to confirm the functionality of our system, we used the well-described TOP-Flash-FFluc reporter^54^. The promoter region of the TOP-Flash reporter contains multiple TCF/LEF binding sites. TCF/LEF factors are transcription factors that bind to DNA in conjunction with β-cat leading to FFluc expression. HEK293T cells expressing TL-βCat were exposed to different pulses of light activation, and as expected, the longer the exposure time of the blue light pulses, the higher the reporter’s response, however, only up to 6h, as 12 h did not increase further. This phenomenon is something we have previously reported for Notch signalling^43^. A longer activation time did not further increase the amount of reporter FFluc (**Fig. 5B**). This, once again, tells the importance of signalling control, as more is not always better, and cells might best to respond to fluctuations rather than continuous signalling^55–58^. To demonstrate specificity and functionality, we analysed β-cat target genes by qPCR, in cells kept in dark and those after blue light activation (**Fig. 5C**).

**Figure 5.**
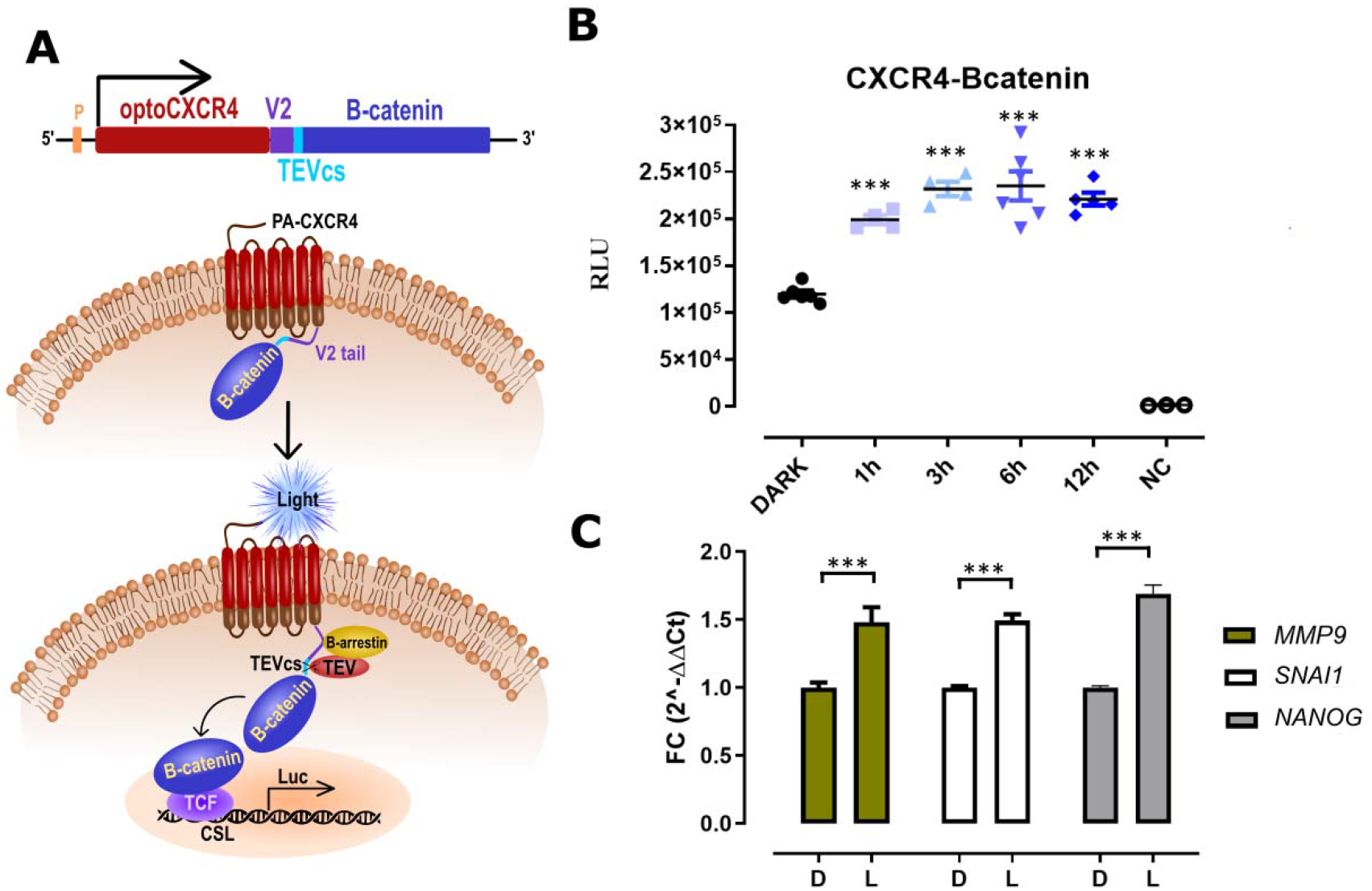
Optogenetic control of WNT signalling. (**A**) Diagram showing the principle where TL is linked to β-Catenin (βCat). (**B**) Relative Firefly luminescence level in HEK293T cells upon photoactivation of TL-βCat for different periods of time. Non transfected cells were used as negative control (NC). (**C**) The mRNA expression of *MMP9*, *SNAI1* and *NANOG* genes was determined by qPCR (2–ΔΔCt) 24 after light stimulation of TL-N1CD HEK293T cells. The results are presented as fold change (FC) values (mean□±□SD) normalized to the *GAPDH* gene expression. All data were analyzed with a one-way ANOVA test and Tukey’s multiple comparisons post-hoc test vs. not light stimulated cells (the time point 0 h and D; dark, respectively). *p□≤□0.05; **p□≤□0.01; ***p□≤□0.001 were considered statistically significant, ns – non significant. All analysed genes showed statistically significant higher expression compared to control cells.

### Cell-cell contact/proximity detection using TANGO-Light

Finally, being inspired by SPARK2^25^ we wanted to assess whether TL activation was possible by presentation of a natural light source. For this we selected *Gaussia* luciferase (Gluc), as it is one of the brightest luciferases, it is normally secreated and the wavelength of its luminescence emission (at around 470 nm) is close to that required by the TL receptor^59^, and fused it to two plasma membrane receptors in “sender” cells.

We have previously characterised an engineered Gluc-Cannabinoid receptor (Gluc-CB1) fusion that is expressed only on the plasma membrane of transfected cells^60^. In addition, we generated yet another receptor Gluc fusion, in this case the CXCR4 receptor, which was placed under the TetON conditional promoter. As with Gluc-CB1, we first assessed that the receptor is only on the plasma membrane and not cleaved or secreted to the medium. For this we transfected HeLa TET-ON cells^59^ with Gluc-CXCR4 plasmid and cultured them in presence of 100 ng/mL doxycycline (Dox) after 24h of transfection. Gluc activity measurement was performed on the following day using Luminometer Synergy H1 (Biotek Instruments, Winooski, VT, USA), which injected 50 mL of 20 mM coelenterazine and record photon counts for 1 second immediately after injecting the substrate. The data were collected for both culture media (from cells with and without addition of coelenterazine) and Gluc-CXCR4 expressing cells in triplicates (**Suppl. Fig. 2B**). Moreover, the activity of Gluc is proportional to the amount of Dox (**Suppl. Fig 2A**).

To analyse whether Gluc on the surface of a different cell (sending) is able to activate the TL system on another cell (receiving), we transfected two cell populations of each cell line, HEK293T and H1299: one with the TL-N1ICD and the 12xCSL-FFluc reporter (receiver cells), while the second population was transfected with a membrane tethered Gluc (Gluc-CB1 or Gluc-CXCR4) (sender cells) (**Fig. 6 A**). We hypothesised that Gluc would be able to photo-activate the light-sensitive TL if in close proximity. A day after transfection, sender and receiver cells were transferred into the same well at three ratios: 1:1, 5:1, and 10:1 sender to receiver cells. To activate the Gluc, we added coelenterazine to the co-cultured cells. After 48h, we then measured the reporter FFluc activity in H1299 (**Fig. 6 B**) and HEK293T cells (**Fig.6 C**). As expected the more donor cells the higher the amount of FFluc luminescence in receiving cells (**Fig.6 B-C**). Note that FFluc and Gluc have different substrates and therefore they can be analysed separately.

**Figure. 6.**
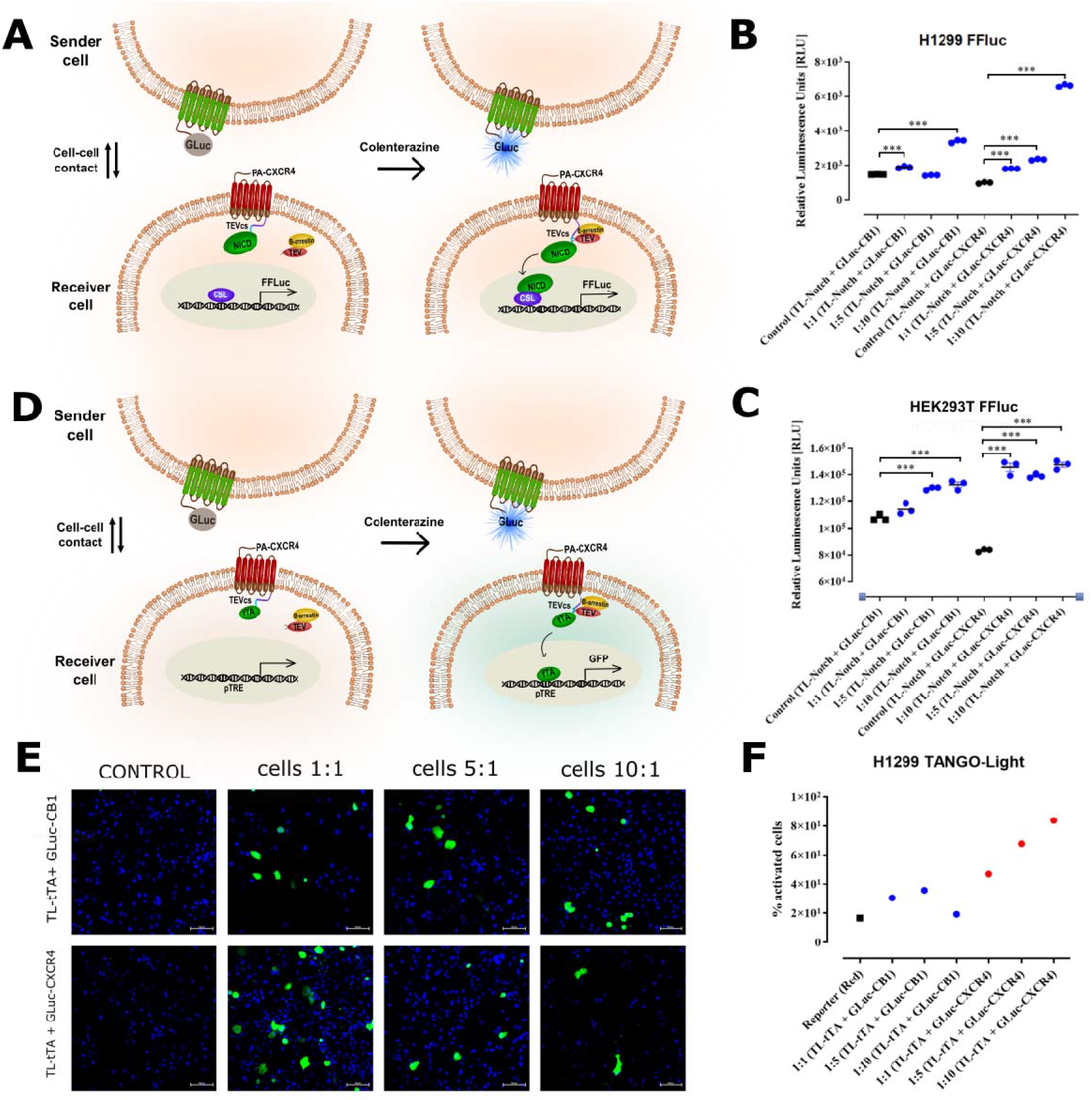
TL can be activated by cell-cell contact or proximity. (**A**) Diagram of structure and action of the system activated by natural source of light –*Gaussia* luciferase (Gluc), for detecting cell-cell contact/proximity. Sender cells expressing Gluc-CB1 or Gluc-CXCR4, and a receiver cells expressing TL-N1ICD, and 12xCSL-FFluc reporter. Relative FFluc luminescence level on H1299 (**B**) and HEK293T (**C**) cells upon activation of TL by Gluc during cell-cell contact/proximity. (**D**). Scheme of structure and action of the system where a sender cells expresses Gluc-CB1 or GLuc-CXCR4, and a receiver cells expressing TL-tTA and the pTRE-GFP reporter. As result of cell-cell contact/proximity, receiver cells be activated by the sender cells and respond by expressing GFP (**E**). Results as percentage of activated cells after flow cytometer analysis of H1299 receiver/sender cells (**F**). Images of two distinct groups of H1299 cells, sender cells expressing Gluc-CB1or Gluc-CXCR4, and receiver cells expressing TL-tTA and GFP reporter. Both populations were co-cultured and then Gluc was activated, or not (negative control cells), by addition of coelenterazine. Cells were then imaged after 48h.

Applying the same concept, we performed and analogical experiment using H1299 cells expressing TL-tTA and the pTRE-GFP reporter (receiver cells), and cells expressing Gluc-CB1 or Gluc-CXCR4 as sender cells. Similarly as described above, after separately transfecting the two H1299 populations, they were then co-cultured at three different ratios: 1:1, 5:10, and 10:1 sender:receiver cells. Coelenterazine was then added to the mix of cells, and the reporter (GFP) expression was analysed 48h after. To quantitate the number of activated receiver cells, we analysed the cells using flow cytometry. A relative quantification was performed by quantifying the number of the cells expressing GFP. We observed a higher amount of cells expressing GFP in wells with 5:1 cells ratio and the highest level of expression with cells in a 10:1 sender:receiver cells ratio when using Gluc-CXCR4 as sender cells (**Fig.6 E-F**). Images of two populations of H1299 cells expressing TL-tTA and pTRE-GFP reporter (receiver cells), and cells expressing Gluc-CB1 or GLuc-CXCR4 (sender cells). When co-cultured, the activated cells showed expression of GFP after the addition of coelenterazine, whereas the negative control cells (without coelenterazine addition) showed no reporter gene expression of GFP.

These results provide proof-of-concept demonstration that TL can be activated by Gluc presented by another cell to enable bioluminescence-mediated transcriptional activation in a cell-cell contact/proximity manner.

## DISCUSSION

Cellular signalling is not just a simple on/off switch. It is a fine-tuned, tightly regulated process involving various biochemical reactions and regulatory mechanisms, that exhibit distinctive kinetics, dynamics and oscillations^60^. In order to understand and control cell signalling, we must mimic cell signalling fluctuations in the context of a complex and dynamic cellular environment. In addition, the control of transcription modulators is critical to understand and regulate the mechanisms that affect cellular signalling pathways, and downstream effects such as cell phenotype, metabolism or fate. Thus, by managing these regulatory molecules, we can effectively control the state of the cell.

In this study, we engineered the synthetic system TANGO-Light (TL) to control the activity of specific transcriptional modulators by optogenetic means. By doing so, we show that we can influence specific downstream gene activation, as well as phenotypical manifestations such as proliferation, and EMT/MET progression. The TL system shows high malleability, as serval signalling pathways could be mimicked (Notch, Wnt) in a dose-dependent manner. We found that continuous signalling does not necessarily result in higher downstream responses e.g. □-Cat when cells were exposed to different lengths of blue light pulses, the reporter’s response increased with longer exposure times, but only up to 6 hours. After that point, the response remained constant, a phenomenon that we have previously reported on Notch^43^. These findings highlight the significance of pattern regulation, shedding light on how cells determine multiple types of responses. This observation is in line with other recent discoveries, for example, the ERK and AKT have opposite effects on pluripotency vs differentiation, and to dissect these, optogenetic systems were generated to find that ERK activity, no matter whether it is short, constant or oscillating is not the key to exit pluripotency, but rather the cumulative amount of activated ERK over time strides phenotypical change^61^. Recovery from ERK-induced pluripotency is counteracted by FGF/AKT receptor dynamics.

Such kinetics of signalling players are becoming better defined, for example FRAP (fluorescence recovery after photobleaching) studies indicate that TFs rapidly recovery after photobleaching, suggesting that a significant number of TF molecules are in fast diffusion states^62^. Studies focused on imaging the dynamics of Sox2 and Oct4 in living embryonic stem (ES) cells have shown that Sox2 and Oct4 molecules use the 3D diffusion-dominant trial-and-error target-seeking mechanism^63^.

We also show that TL system can be activated upon cell-cell contact or proximity, which might open doors or allow cell-cell activation, similar to cell-cell contact such as Notch, which holds the potential for multiple applications. One potential use is in inducing differentiation. Specific signalling cascades can be activated by triggering the TL system upon cell-cell contact to guide stem cells towards desired cell lineages. The TL system could also play a role in tissue development and organization. Activation through cell-cell contact may help in the precise arrangement of cells during embryogenesis or tissue regeneration. Modulating immune responses is another example. By utilizing the TL system, immune cells could be selectively activated upon contact with target cells, allowing for precise control over immune reactions. Summarizing, from a synthetic biology point of view, triggering the TL system through cell-cell contact provides opportunities for engineering artificial cellular networks and designing complex cellular circuits that respond to specific cell-cell interactions.

Optogenetics is likely the most precise system to control the impact of cell signalling on cells and determines their fate due to its rapid response, tight control non-invasive nature. While biological processes have been traditionally regulated using biochemical methods, the use of optogenetic has become more and more commonplace, and the reason behind this is due to the specificity and high spatio-temporal resolution of optogenetic tools.

Transcription factor-based genetic tools are widely used in synthetic/cellular biology applications, such as gene regulation and signal transduction. So far, these biosensors can be used to engineer cells to respond to specific stimulations and produce desired outputs, such as therapeutic proteins. TANGO-Light, together with other synthetic light-induced systems such as iTango and SPARK2^23^ as genetic actuators, have broad applications in signal transduction and cell differentiation processes. Moreover, they can be easily translated into *in vivo* contexts.

What distinguishes the system we have engineered is that, compared to other systems, it does not require the presence of a ligand. Only blue light is required for activation, and as a result, we get a quick and specific responses. In additional, we show the flexibility of TL, where the transcriptional modulators can be easily exchanged like LEGO.

## EXPERIMENTAL SECTION

### Materials and reagents

KOD-Xtreme hot-start DNA polymerase (Merck Millipore), DreamTaq™ Green PCR Master Mix (ThermoFisher Scientific), *DpnI* restriction enzyme (NEB), Gibson Assembly® Master Mix (NEB), ampicillin (BRAND), DNA Clean & Concentrator and Zyppy Plasmid Kits (Zymoresearch), Turbofect™ Transfection Reagent (ThermoFisher Scientific), Lipofectamine 3000™ (ThermoFisher Scientific),Vybrant™ DiD Cell-Labeling Solution (ThermoFisher Scientific), Bright-Glo Luciferase Assay System (Promega), Dulbecco’s Modified Eagle Medium (DMEM), fetal bovine serum (PromoCell), plasmids pCDNA3.1(+)-CMV-βarrestin2-TEV (Addgene plasmid #107245; http://n2t.net/addgene:107245; RRID:Addgene_107245) and Flag Snai1 6SA (Addgene plasmid #16221; http://n2t.net/addgene:16221; RRID:Addgene_16221), plasmid pDONR223_NOTCH1_ICN (Addgene plasmid #82087; http://n2t.net/addgene:82087; RRID:Addgene_82087), plasmid GPCR-TANGO and HTLA cells were a gift from the lab of Richard Axel, Howard Hughes Medical Institute, Department of Biochemistry and Cellular Biophysics, Center for Neurobiology and Behavior, Columbia University, New York, plasmid PA-CXCR4 was a gift from the lab of Minsoo Kim, Ph.D, University of Rochester Medical Center School of Medicine and Dentistry. Primary anti-FLAG mouse antibody (Cell Signaling Technology, Cat. Nr: #8146), and secondary anti-mouse antibody conjugated with Alexa Fluor 555 dye (Invitrogen Cat. Nr: A-31570). Primary anti-NOTCH1 (D1E11) rabbit monoclonal antibody (Cell Signaling Technology, Cat. Nr: #3608), secondary anti-rabbit antibody conjugated with AlexaFluor 532 (Invitrogen Cat. Nr: #A-11009). Secondary antibody: Peroxidase F(ab’)2 Fragment Donkey Anti-Rabbit IgG (H□+□L) (711-036-152, Jackson ImmunoResearch). Immunocytochemistry reagents: Alexa Fluor 488 Tyramide Reagent (B40953, LifeTechnologies), Hoechst 33342 (Cayman), ProLong Gold mounting medium (LifeTechnologies). All PCR primers were bought from Genomed (Warsaw, Poland).

### Molecular cloning

A plasmid containing fusion of PA-CXCR4 with C-terminus of the vasopressin receptor (V2 tail), TEV cleavage site and tTA (PA-CXCR4-TANGO) and inducible expression plasmid of PA-CXCR4-SNAI1 and PA-CXCR4-N1ICD were generated by Gibson Assembly^64^ by combining products from two PCR reactions previously carried out with the addition of homology arms. The same was done for PA-CXCR-Elf3 and PA-CXCR4-SOX 2 constructs. PCR products was digested with *DpnI* to remove methylated DNA and purified using DNA purification kits. The Gibson products were transformed into previously prepared electrocompetent *E.coli* bacteria^65^. The resulting bacterial colonies, selection with ampicillin 100 mg/mL, were subjected to colony PCR and sequence-verified.

### Cell culture and transfection

HTLA cells^28^, human embryonic kidney 293 (HEK293) cell line stably expressing a tTA-dependent luciferase reporter and a β-arrestin2-TEV fusion gene, lung cancer cell line H1299, breast cancer cell line MDA-MB-468 were grown in Dulbecco’s Modified Eagle Medium (DMEM), MDCK-2 an epithelial-like Madin-Darby canine kidney cell line were grown in Eagle’s Minimum Essential Medium (EMEM). All media were supplemented with 10% FBS, penicillin (100 units/mL), and streptomycin (100 μg/mL). Cells were grown at 37°C in an atmosphere of 5% CO_2_.

One day before transfection HTLA cells were seeded 5 x 10^4^ cells/well in a 24-well plate and transfected with the PA-CXCR4-TANGO using Turbofect™ Transfection Reagent following the manufacturer’s protocol.

Lung cancer cells H1299 24h before transfection were seeded 5 x 10^4^ cells/well in a 24-well plate and -transfected with the β-arrestin2-TEV and co-transfected with PA-CXCR4-TANGO or PA-CXCR4-TANGO-SNAI1 using Turbofect™ Transfection Reagent following the manufacturer’s protocol.

Similarly, 24h before transfection lung cancer cells H1299 and breast cancer cells MDA-MB-460 were seeded 5 x 10^4^ cells/well in a 24-well plate and -transfected with the β-arrestin2-TEV and co-transfected with PA-CXCR4-TANGO or PA-CXCR4-TANGO-N1ICD using Turbofect™ Transfection Reagent following the manufacturer’s protocol.

Highly epithelial MDCK-2 cells, were seeded 5 x 10^4^ cells/well in a 24-well plate and - transfected with the β-arrestin2-TEV and co-transfected with PA-CXCR4-TANGO or PA-CXCR4-TANGO-SNAI1 using Lipofectamine3000™ Transfection Reagent following the manufacturer’s protocol. Transfection was done in triplicate. After light activation cells were stained with Vybrant™ DiD Cell-Labeling Solution for better visualization under the confocal microscope.

### Immunocytochemistry and confocal microscopy

TANGO-Light, TL-SNAI1 and TL-N1ICD H1299 cells were seeded on glass bottom Labtec 8-chamber slides (Nunc) at a density of 4□×□10^5^ cells/mL. The cells on one slide were photo-activated by 5 ms blue light pulses, while those on the second slide were kept in the dark. Cells were then washed with PBS and fixed 1:1 Acetone: Methanol for 30 min at −□20 °C. Next, cells were washed with PBS and incubated in Blocking Buffer (BB) for 1 h at room temperature, followed by overnight (4°C) incubation with primary rabbit antibodies against N1ICD diluted 1:500 in BB primary anti-FLAG mouse antibody diluted 1:1000 in BB. After triple washes in PBS, cells were incubated for 1 h (room temperature) with Secondary Peroxidase F(ab’)2 Fragment Donkey Anti-Rabbit IgG (H□+□L) diluted 1:1000 in BB or secondary anti-mouse antibody conjugated with Alexa Fluor 555 dye diluted in 1:1000 in BB. After three PBS washes, cells were stained with Alexa Fluor 488 Tyramide Reagent following to manufacturer’s protocol. Reaction was stopped through incubation in 3% hydrogen peroxide. The nuclei of cells before imaging have been Hoechst 33342 stained (concentration 10 µg/mL). Next, cells were mounted with ProLong Gold mounting medium and visualized under a Nikon ECLIPSE Ti confocal microscope.

### Photoactivation

Transfected HTLA cells, lung cancer cell lines H1299, H2170 and breast cancer MDA-MB-468 cells were photo-stimulated using the MAGI-01 Opto-stimulation system (Radiometech) (**Suppl. Fig. 1)** (previously reported here: optoNotch^43^) with blue LEDs (456□±□2 nm), at an intensity of 3.2 W/m2, in pulses of 0.05 s luminous and 5 s breaks for 1, 3, or 12 h, while unstimulated cells were kept in the dark. Light-stimulation was performed inside the cell culture incubator to avoid changes in environmental conditions for the photo-stimulated cells.

### Luciferase Reporter Assay

At 48 hours after transfection and blue light activation, the cells were resuspended in 150 µl od medium. The equal volume of lysates and luciferase reagent was transferred to a black microplate well and measured for luciferase activity by triplicate using a Bright-Glo Luciferase Assay System with a microplate luminometer (Tecan Infinite 2000 PRO). The results were then statistically analyzed.

### Cell-cell contact/proximity

For this purpose we used an engineered Gluc-Cannabinoid (Gluc-CB1) receptor fusion that is expressed only on the plasma membrane of transfected cells. We used the Gluc-CB1^T210A^ mutation, which results in increased localization in the cell membrane relative to the wild-type receptor^66, 67^. In addition, we used another engineered receptor Gluc fusion, in this case the CXCR4 receptor (Gluc-CXCR4), in which the *Gaussia* Luciferase (Gluc) was fused to the *N*-terminal of CXCR4 receptor.

We transfected two cell populations of H1299 cell line, HEK293T and H1299 separately with the TL-N1CD and the 12xCSL-FFluc reporter (Receiver cells), and the second population separately was transfected with a membrane receptor (CB1 or CXCR4) fused Gluc (Sender cells). After transfection we co-cultured cells together in the same well at three ratios: 1:1, 5:1, and 10:1 sender to receiver cells. To activate the Gluc-CXCR4 or Gluc-CB1 receptors in the TL system, we exposed the co-cultured cells to coelenterazine. Then, 48h after activation, we measure the response in receiver cells.

Analogically to the same concept, we prepared H1299 cells transfected with the same scheme of Sender (Gluc-CXCR4 or Gluc-CB1) and Receiver cells (TL-tTA/βarrestin-TEV and pTRE-GFP reporter). Similarly as described above, after separately transfecting these two H1299 populations, they were then co-cultured at three different ratios: 1:1, 5:10, and 10:1 sender:receiver cells. Coelenterazine was then added to the mix of cells and the reporter was visualised 48h after. To quantitate the activated receiver cells, we analysed the cells using flow cytometry. A relative quantification was performed by quantifying the number of the cells expressing GFP.

### Measuring Gaussia Luciferase activity

TET-ON HeLa cells were seed at density of 10,000 cells per well in 96-well plate and incubate at 37oC, 5% CO2. In the next day, cells were transfected with Gluc-CXCR4 plasmid using Lipofectamine™ 3000 Transfection Reagent (Thermo Fisher Scientific, USA). 100 ng/mL Doxycycline (Dox) was added to cell culture media after 24h of transfection and Gluc activity measurement was performed by the following day using Luminometer Synergy H1 (Biotek Instruments, Winooski, VT, USA), which is set to inject 50 µL of 20 mM coelenterazine and to immediately record photon counts for 1 seconds for each well. The data were collected for both culture media and Gluc-CXCR4 expressing cells in triplicates.

### 3D sandwich assay

The Matrigel at a concentration of 4 mg/ml in medium (lower gel) was poured onto the bottom of a 96-well plate, and incubated at 37 °C for 30–60 min. Then, single cells suspension of either, H1299, H2170 and MDA-MB-468, WT or TL-tTA, TL-SNAI1 and TL-N1ICD, were admixed with Matrigel (upper gel, final concentration of 2 mg/ml of Matrigel) and seeded at a density of 4□×□10^4^ cells/mL. In one of the variant into the medium with cells, and on the top of the gel was added medium with TGFβ. The plates were centrifuged for 5 min, and incubated at 37°C overnight. The cells in one plate were light activated by pulses of 0.05 s every 5 s for 3 h every day during the experiment, while an identical second plate remained in the same conditions but kept in the dark. The organoids were observed by real-time visualization in IncuCyte live-cell imager (Essen Bioscience).

### RNA extraction and RT-qPCR

Trypsin was used to detach the cells after the experiment. Cell were then collected by centrifugation (400*×g*). After removing the supernatant, the cells pellets were lysed according to the ExtractMe Total RNA kit (Blirt) manufacturer’s protocol. cDNAs were synthesized using the High-Capacity cDNA Reverse Transcription Kit with the addition of a RNase Inhibitor (Applied Biosystems). All primers used for qPCR were tested for specificity and sensitivity. Glyceraldehyde 3-phosphate dehydrogenase (GAPDH) was used as a housekeeping gene. PCR reactions were performed with PowerUp SYBR Green Master Mix (Applied Biosystems) through the LightCycler® 480 II instrument (Roche) in triplicates on 96-well plates. The number of cycles needed to reach a specific threshold of detection (CT) was used to calculate relative quantification (RQ). Relative mRNA expression was calculated using the delta CT subtraction and normalized to the expression of *GAPDH*.

### Statistical Analyses

Statistical analyses were performed using GraphPad Prism 5.0 (GraphPad Software Inc., California, U.S.A). Significance among luminescence readings was assessed by one-way ANOVA followed by Tukey’s *post-hoc* test (*, p < 0.05;**, p < 0.01; ***, p < 0.001).

All experiments were performed in at least triplicate, and independently repeated minimum 3 times.

## AUTHOR CONTRIBUTIONS

AP-P and JK performed the experiments and analyses. JC assisted in molecular cloning. THN cloned and characterised the Gluc-CXCR4 donor receptor. AC assisted in β-catenin experiment, and flow cytometer analyses. AP-P, JK and AR-M designed the experimental section of this research work and co-wrote the manuscript draft. AR-M supervised the overall study. All authors read and approved the final manuscript.

## ACKNOWLEDGEMENTS

The authors would like to thank Matthias Nees, Mervi Toriseva and colleagues from FICAN West Cancer Centre Laboratory at the University of Turku in Finland for the warm welcome, Sima – Finnish Spring Mead, and the pleasant working environment in the laboratory. We would like to thank Minsoo Kim and his colleagues who kindly provided the PA-CXCR4 plasmid. The authors would like to also thank Addgene, and the colleagues depositing their plasmids there, for the support to the greater scientific community.

## DECLARATIONS

### Ethical approval

Not needed

