## Supplementary material for "TANGO-Light - optogenetic control of transcriptional modulators": Suppl. Material

**Statements:**

All the raw data supporting the findings in this work can be obtained on request from the

corresponding author.

Plasmids used in this work will be available via Addgene and on request from the corresponding author.

**Plasmids**

For our experiments we use CB1^T210A^ mutation, which results in unaffected expression.

**Nucleic acid Sequence of the construct GLuc- FLAG-CB1^T210A^**

gagggcctatttcccatgattccttcatatttgcatatacgatacaaggctgttagagagataattggaattaatttgactgtaaacacaaagatattagtacaaaatacgtgacgtagaaagtaataatttcttgggtagtttgcagttttaaaattatgttttaaaatggactatcatatgcttaccgtaacttgaaagtatttcgatttcttggctttatatatcttGTGGAAAGGACGAAACACCggGTCTTCgaGAAGACctGTTTTAGAGCTAGAAatagcaagttaaaataaggctagtccgttatcaacttgaaaaagtggcaccgagtcggtgctttttTgttttagagctagaaatagcaagttaaaataaggctagtccgtTTTTagcgcgtgcgccaattctgcagacaaatggctctagaggtacccgttacataacttacggtaaatggcccgcctggctgaccgcccaacgacccccgcccattgacgtcaatagtaacgccaatagggactttccattgacgtcaatgggtggagtatttacggtaaactgcccacttggcagtacatcaagtgtatcatatgccaagtacgccccctattgacgtcaatgacggtaaatggcccgcctggcattGtgcccagtacatgaccttatgggactttcctacttggcagtacatctacgtattagtcatcgctattaccatggtcgaggtgagccccacgttctgcttcactctccccatctcccccccctccccacccccaattttgtatttatttattttttaattattttgtgcagcgatgggggcggggggggggggggggcgcgcgccaggcggggcggggcggggcgaggggcggggcggggcgaggcggagaggtgcggcggcagccaatcagagcggcgcgctccgaaagtttccttttatggcgaggcggcggcggcggcggccctataaaaagcgaagcgcgcggcgggcgggagtcgctgcgacgctgccttcgccccgtgccccgctccgccgccgcctcgcgccgcccgccccggctctgactgaccgcgttactcccacaggtgagcgggcgggacggcccttctcctccgggctgtaattagctgagcaagaggtaagggtttaagggatggttggttggtggggtattaatgtttaattacctggagcacctgcctgaaatcactttttttcaggttGGaccggtgccaccATGGGAGTCAAAGTTCTGTTTGCCCTGATCTGCATCGCTGTGGCCGAGGCCAAGCCCACCGAGAACAACGAAGACTTCAACATCGTGGCCGTGGCCAGCAACTTCGCGACCACGGATCTCGATGCTGACCGCGGGAAGTTGCCCGGCAAGAAGCTGCCGCTGGAGGTGCTCAAAGAGATGGAAGCCAATGCCCGGAAAGCTGGCTGCACCAGGGGCTGTCTGATCTGCCTGTCCCACATCAAGTGCACGCCCAAGATGAAGAAGTTCATCCCAGGACGCTGCCACACCTACGAAGGCGACAAAGAGTCCGCACAGGGCGGCATAGGCGAGGCGATCGTCGACATTCCTGAGATTCCTGGGTTCAAGGACTTGGAGCCCATGGAGCAGTTCATCGCACAGGTCGATCTGTGTGTGGACTGCACAACTGGCTGCCTCAAAGGGCTTGCCAACGTGCAGTGTTCTGACCTGCTCAAGAAGTGGCTGCCGCAACGCTGTGCGACCTTTGCCAGCAAGATCCAGGGCCAGGTGGACAAGATCAAGGGGGCCGGTGGTGACggaCTCGAGGATTACAAGGATGACGACGATAAGAAGTCGATCCTAGATGGCCTTGCAGATACCACCTTCCGCACCATCACCACTGACCTCCTGTACGTGGGCTCAAATGACATTCAGTACGAAGACATCAAAGGTGACATGGCATCCAAATTAGGGTACTTCCCACAGAAATTCCCTTTAACTTCCTTTAGGGGAAGTCCCTTCCAAGAGAAGATGACTGCGGGAGACAACCCCCAGCTAGTCCCAGCAGACCAGGTGAACATTACAGAATTTTACAACAAGTCTCTCTCGTCCTTCAAGGAGAATGAGGAGAACATCCAGTGTGGGGAGAACTTCATGGACATAGAGTGTTTCATGGTCCTGAACCCCAGCCAGCAGCTGGCCATTGCAGTCCTGTCCCTCACGCTGGGCACCTTCACGGTCCTGGAGAACCTCCTGGTGCTGTGCGTCATCCTCCACTCCCGCAGCCTCCGCTGCAGGCCTTCCTACCACTTCATCGGCAGCCTGGCGGTGGCAGACCTCCTGGGGAGTGTCATTTTTGTCTACAGCTTCATTGACTTCCACGTGTTCCACCGCAAAGATAGCCGCAACGTGTTTCTGTTCAAACTGGGTGGGGTCACGGCCTCCTTCACTGCCTCCGTGGGCAGCCTGTTCCTCGCAGCCATCGACAGGTACATATCCATTCACAGGCCCCTGGCCTATAAGAGGATTGTCACCAGGCCCAAGGCCGTGGTGGCGTTTTGCCTGATGTGGACCATAGCCATTGTGATCGCCGTGCTGCCTCTCCTGGGCTGGAACTGCGAGAAACTGCAATCTGTTTGCTCAGACATTTTCCCACACATTGATGAAACCTACCTGATGTTCTGGATCGGGGTCACCAGCGTACTGCTTCTGTTCATCGTGTATGCGTACATGTATATTCTCTGGAAGGCTCACAGCCACGCCGTCCGCATGATTCAGCGTGGCACCCAGAAGAGCATCATCATCCACACGTCTGAGGATGGGAAGGTACAGGTGACCCGGCCAGACCAAGCCCGCATGGACATTAGGTTAGCCAAGACCCTGGTCCTGATCCTGGTGGTGTTGATCATCTGCTGGGGCCCTCTGCTTGCAATCATGGTGTATGATGTCTTTGGGAAGATGAACAAGCTCATTAAGACGGTGTTTGCATTCTGCAGTATGCTCTGCCTGCTGAACTCCACCGTGAACCCCATCATCTATGCTCTGAGGAGTAAGGACCTGCGACACGCTTTCCGGAGCATGTTTCCCTCTTGTGAAGGCACTGCGCAGCCTCTGGATAACAGCATGGGGGACTCGGACTGCCTGCACAAACACGCAAACAATGCAGCCAGTGTTCACAGGGCCGCAGAAAGCTGCATCAAGAGCACGGTCAAGATTGCCAAGGTAACCATGTCTGTGTCCACAGACACGTCTGCCGAGGCTCTGTGAGCCTGATGCCTCCCTGGCAGCACAGGAAAAGAATTTTTTTTTTTAAGCTCAAAATCTAGAAGAGTCTATTGTCTCCTTGGTTATATTTTTTTAACTTTACCATGCTCAATGAAAAGGTGATTGTCACCATGATCACTTATCAGTTTGCTAATGTTTCCATAGTTTAGGTACTCAAACTCCATTCTCCAGGGGTTTACAGTGAAGAAAGCCTGTTGTTTAAGTGACTGAACGATCCTTCAAAGTCTCAATGAAATAGGAGGGAAACCTTTGGCTACACAATTGGAAGTCTAAGAACCCATGGAAAAATGCCATCAAATGAATAATGCCTTTGTAACCACAACTTTCACTATAATGTGAAATGTAACTGTCCGTAGTATCAGAGATGTCCATTTTTACAAGTTATAGTACTAGAGATATTTTGTAAAATGTATTATGTCCTGTGAGATGTGTATCAGTGTTTATGTGCTATTAATATTTGTTTAGTTCAGCAAAACTGAAAGGTAGACTTTTATGAGAACAATGGACAAGCAGTGGATACGTGTCAATGTGTGCACTTTTTTTCTATATTATTGCCCATGATATAACTTTAGAAATAAACCTTAATATTTCTTCAAATATCTCTATTTAATTTTGACACTGAAATAACCGTAAAGGTTTATTTTTCTGTTACCTCAACAAGAAGAATTTGAAGACTTCAAAATATTGAGCAGAATTCATTCATACTTAAAAATTTATTAGCCCTGCATTTTCATAGGAAGACACATTATCTTCTGGACTATAGCTGTTCTAATGGATTATAATCAGAATGGAAGAGAGAAAGCATATTGACTTTTTTTGAGCGACATCTCTGACTTTCTTTAGTCTTTAGCTATTACTGGATCTCTTAAGACAGCATGTGTTAATCTTAATGTATATCGTTATCACTGTGCAGTTGCTGTTTACTTGAATAGTATTGTGTTCCTATATTCCAGGTTTAAGTAGATTTCATGCCTGGGTGGCCAAACAACAGTCTTCATTTTTTTTAATTGAAAAGAAGTAGTGTCTGGATCAGTAAAATTATACTGTGTGTGAGTGTGAATATAAATGTGTGTATGTGTGTTTCTGTCCTGTAACTGTTACAGTAATGTCATAAAGTGAGAAAACTGTGACCAAGTATAAACTTTTACCACTTGCTGCACTCTTGCACATGGATTCAGTTTCTAAAATTGAGTTCTTCCTGTAATCTTGTTGATAAAAATACTGACTCCAACCATTCAAAAATTTCACCCCATCCCTCCTTAAGAGATTGGATCAAGTATTACTAAATTGACCTTTAGGTATTACACAAGACCAGTGCTTAGCAAAAAATAATGACAGGCATCCAAGGAAGGGATGTATTTGTAGTGTTATTGCCAGGAAAGGAGAGTACTTTGGTTTCTGAGCACCGAATATTGAGCAATATGTCAGTCACTAAAAGGAAGACAGTTCTACAGAAAAACAATGGTAACATTTTTCAATAGCGTGTGTAGATAGTATGCACTATATACATCACGTTAAAGTAGGACTATCACACCCAGCCCATGTGGCTAAAAAAGCTGAATCAGACAGTGGATGAGACACACAACGGCAGTGAAGAACCGATACACTTGGCATTGACGTCTAGCTATGCTGTATCTGTGCTTTGCCCACATGCCCTTGGTGACAGCTGAGCACCCAGCTCTGTCTTGGTAGGTTTGGGCTAAGGAACAAATCTCTCCTTTGCTCGTGGTTAGCAAGATACACTCAAGCATGAAGATAAACACAGCTGCTTTCTTCTTACACCCCGGTCTCATGCTCCTTAATGGCGCCATGGGTGCTTGTTGGGCCTTTTTCCAGTAAGGAATGATATTGCTGAAGAATCTACTTAACCCTGACAAATTTTAATTATAATCTCTTCTTATACAGATAAAACATGACTCCTACAAGGCCCCAAGGTTTACATAGTCTGAAGTGAAGTACAGAGCTGGCATCTATCTGGTGATTTCTAGCTCTCGAGATACCCAAGCAGCCTGATGGGGCAGTTCCCCTTCTTACGGTTCACGCTCTAAGGCAGGATGTGGCTTATGAGATACTTTGCATTGTCTGTCTGCACACCTTGAATCTGCCTGCTGGCTCCCTTACTTTACCTCTCTGTCATGTGCAGATGAAGGCTCAGGGTGCTAGAGGATTAGTAAGATCTCTTTCTAAAGACAGGAGAGATTATTTACAAGAAGAACTCACCAGGGTTTAGTTTGCATTTAAGAATTGCCAGTCTTTTGTCCTGCATCATCTTGAACATTAATCCACATGTTTCAGAGCTCACCAGGCAGTACCAATGCTCTTTTCACAGCTATGAAGAGCTAGAGAAATTCTTGTTATGGTAGAAAAATTTCACGATTCATTTTTGAAACTGCATTTGTGCGTATGCAGTGTAGATTTTATAGTGTGTTGTGCTTTCAAGATCTAAATCATATATAATAAATTAAGGGACAATGGGGCTGACAGCACTAAACTTGGTGCTTATTGATATTCTAAGAAATATCTGTGAAATATCATCACGTATGTTATACAACCTTCATTTAAAAAGGTTTAAAACTAGTTAGATTCACTTTGACACTTTTCATATCATTTCTTAACCCAAGTGACGAAAACATTGTCCCCAATGAATATACTCATTAGAATTACCATTTGTTAATATCACTCATTAATTAACCCCATAATTAGATCCATTAATTTAAATGATTTAAATTTAAGTAAGTTTTATAAGGTCTGACATCAGAGGTATCTTACTTTCCTCTGAGGATGATGTACTTGCCCTGACCATGCATTTTACCATCACACATGTTCAGAAAGGGCCAAATTCCCAACCTGCTCATTTTTTTTTTATCAGAGTCATGATGAATCAGTCCTAGAATGTTTCATTTGCACAAGTAGGGCTGCCTCCAAGAGGAACCTCTGATTTATTTTGTATGAAATATATGTGAAAGGATATGAATCTGAGAGATGCTGTAGACATCTGTCCTACACTTGAGATGATTTCCAAGCCTCTCTGGCACTTTGAGTTAAGTCTATCTGGTATTAAATGCCAAGGACCTTTTGCTGCCTAAATCCACTCTGCAGGAAATAGGCCCAACCACCAGATGAGAATTAGGCCCTGGATGAGTAGCGCTATAGTTACTGTCCTGTTGATTAATTTCTGCCATTTCATGTCCATAAAAGAGACCACCCATATCATGCACACAATTAGATTTCTCACACTCTAACTGTATATTTGTATGATATTTTAAAATCTCCTAAATGCTGGGCAATGGCTATTAACAATTAATTGTCTTGCACTGGCCTTCTGATGAAATGTTAACAATGCCTATTGTAATATAGAAAAAAACATTCTATCTACTGATTTGGGCTGAATGTATGTAAATAGGTTTCTAAAAAGTCAGATGTTTGAGCAGTGGCCTACAAATCAGTAATTTTCGGATGGGAGAGTTTCTTTACATTGCCGTGGCATCTTAAAAGCTATCTTCATGTAAATTGACTGTACTAGGCCTACTGGGGATCAGAGTTCCCAAGAAAGGAAACCTTTTCTTGTATCTGGATTCAAATTTATTTCCAATGTTTCAAGCGGGAAACATGACTCTTTATTGTCTGTAAATCTAACATTATTACTTTTCCTCTTAGAAGAATATTGTATTGTTAGATGTTTGTTGAGCTGGTAACATCGTTGCAACCACTGCAATATCTTCGTTAGTAATCTGTATAATACTTTGTATACAAGTACTGGTAAGATTGTTATTAAATGTAGCTTCAGTCATTAAATTACTATAGCAAAGTAGTACTTCTTCTGTAATATTTACAATGTATTAAGCCCACAGTATATTTTATTTCAATGTAATTAAACTGTTAACTTATTCAAAGAGAAAACATCTCATCATGTCTATTGTCCAAAGTTACCTGGAATCAAATAAAAATTCTAGATTACCATGAAGAACATAAAATGCCTTTGAACTCTGCCTTATTTCACAGTCTGATGGCAAAAAATGAGGAAATTGCATCGCATTGTCTGAGTAGGTGTCATTCTATTCTGGGGGGTGGGGTGGGGCAGGACAGCAAGGGGGAGGATTGGGAAGAgAATAGCAGGCATGCTGGGGAgcggccgcaggaacccctagtgatggagttggccactccctctctgcgcgctcgctcgctcactgaggccgggcgaccaaaggtcgcccgacgcccgggctttgcccgggcggcctcagtgagcgagcgagcgcgcagctgcctgcaggggcgcctgatgcggtattttctccttacgcatctgtgcggtatttcacaccgcatacgtcaaagcaaccatagtacgcgccctgtagcggcgcattaagcgcggcgggtgtggtggttacgcgcagcgtgaccgctacacttgccagcgccctagcgcccgctcctttcgctttcttcccttcctttctcgccacgttcgccggctttccccgtcaagctctaaatcgggggctccctttagggttccgatttagtgctttacggcacctcgaccccaaaaaacttgatttgggtgatggttcacgtagtgggccatcgccctgatagacggtttttcgccctttgacgttggagtccacgttctttaatagtggactcttgttccaaactggaacaacactcaaccctatctcgggctattcttttgatttataagggattttgccgatttcggcctattggttaaaaaatgagctgatttaacaaaaatttaacgcgaattttaacaaaatattaacgtttacaattttatggtgcactctcagtacaatctgctctgatgccgcatagttaagccagccccgacacccgccaacacccgctgacgcgccctgacgggcttgtctgctcccggcatccgcttacagacaagctgtgaccgtctccgggagctgcatgtgtcagaggttttcaccgtcatcaccgaaacgcgcgagacgaaagggcctcgtgatacgcctatttttataggttaatgtcatgataataatggtttcttagacgtcaggtggcacttttcggggaaatgtgcgcggaacccctatttgtttatttttctaaatacattcaaatatgtatccgctcatgagacaataaccctgataaatgcttcaataatattgaaaaaggaagagtatgagtattcaacatttccgtgtcgcccttattcccttttttgcggcattttgccttcctgtttttgctcacccagaaacgctggtgaaagtaaaagatgctgaagatcagttgggtgcacgagtgggttacatcgaactggatctcaacagcggtaagatccttgagagttttcgccccgaagaacgttttccaatgatgagcacttttaaagttctgctatgtggcgcggtattatcccgtattgacgccgggcaagagcaactcggtcgccgcatacactattctcagaatgacttggttgagtactcaccagtcacagaaaagcatcttacggatggcatgacagtaagagaattatgcagtgctgccataaccatgagtgataacactgcggccaacttacttctgacaacgatcggaggaccgaaggagctaaccgcttttttgcacaacatgggggatcatgtaactcgccttgatcgttgggaaccggagctgaatgaagccataccaaacgacgagcgtgacaccacgatgcctgtagcaatggcaacaacgttgcgcaaactattaactggcgaactacttactctagcttcccggcaacaattaatagactggatggaggcggataaagttgcaggaccacttctgcgctcggcccttccggctggctggtttattgctgataaatctggagccggtgagcgtggaagccgcggtatcattgcagcactggggccagatggtaagccctcccgtatcgtagttatctacacgacggggagtcaggcaactatggatgaacgaaatagacagatcgctgagataggtgcctcactgattaagcattggtaactgtcagaccaagtttactcatatatactttagattgatttaaaacttcatttttaatttaaaaggatctaggtgaagatcctttttgataatctcatgaccaaaatcccttaacgtgagttttcgttccactgagcgtcagaccccgtagaaaagatcaaaggatcttcttgagatcctttttttctgcgcgtaatctgctgcttgcaaacaaaaaaaccaccgctaccagcggtggtttgtttgccggatcaagagctaccaactctttttccgaaggtaactggcttcagcagagcgcagataccaaatactgtccttctagtgtagccgtagttaggccaccacttcaagaactctgtagcaccgcctacatacctcgctctgctaatcctgttaccagtggctgctgccagtggcgataagtcgtgtcttaccgggttggactcaagacgatagttaccggataaggcgcagcggtcgggctgaacggggggttcgtgcacacagcccagcttggagcgaacgacctacaccgaactgagatacctacagcgtgagctatgagaaagcgccacgcttcccgaagggagaaaggcggacaggtatccggtaagcggcagggtcggaacaggagagcgcacgagggagcttccagggggaaacgcctggtatctttatagtcctgtcgggtttcgccacctctgacttgagcgtcgatttttgtgatgctcgtcaggggggcggagcctatggaaaaacgccagcaacgcggcctttttacggttcctggccttttgctggccttttgctcacatgt

**Nucleic acid Sequence of the construct pTRE-SP-HA-GLuc-CXCR4**

AGAATTCCTCGAGTTTACTCCCTATCAGTGATAGAGAACGTATGAAGAGTTTACTCCCTATCAGTGATAGAGAACGTATGCAGACTTTACTCCCTATCAGTGATAGAGAACGTATAAGGAGTTTACTCCCTATCAGTGATAGAGAACGTATGACCAGTTTACTCCCTATCAGTGATAGAGAACGTATCTACAGTTTACTCCCTATCAGTGATAGAGAACGTATATCCAGTTTACTCCCTATCAGTGATAGAGAACGTATAAGCTTTAGGCGTGTACGGTGGGCGCCTATAAAAGCAGAGCTCGTTTAGTGAACCGTCAGATCGCCTGGAGCAATTCCACAACACTTTTGTCTTATACCAACTTTCCGTACCACTTCCTACCCTCGTAAAGTCGACATGAAGCAGCGGTTCTCGGCGCTGCAGCTGCTGAAGCTGCTGCTGCTGCTGCAGCCGCCGCTGCCACGAGCGTACCCATACGATGTTCCAGATTACGCTCTGCGCGCCAAGCCCACCGAGAACAACGAAGACTTCAACATCGTGGCCGTGGCCAGCAACTTCGCGACCACGGATCTCGATGCTGACCGCGGGAAGTTGCCCGGCAAGAAGCTGCCGCTGGAGGTGCTCAAAGAGATGGAAGCCAATGCCCGGAAAGCTGGCTGCACCAGGGGCTGTCTGATCTGCCTGTCCCACATCAAGTGCACGCCCAAGATGAAGAAGTTCATCCCAGGACGCTGCCACACCTACGAAGGCGACAAAGAGTCCGCACAGGGCGGCATAGGCGAGGCGATCGTCGACATTCCTGAGATTCCTGGGTTCAAGGACTTGGAGCCCATGGAGCAGTTCATCGCACAGGTCGATCTGTGTGTGGACTGCACAACTGGCTGCCTCAAAGGGCTTGCCAACGTGCAGTGTTCTGACCTGCTCAAGAAGTGGCTGCCGCAACGCTGTGCGACCTTTGCCAGCAAGATCCAGGGCCAGGTGGACAAGATCAAGGGGGCCGGTGGTGACGGCGGCGGTGGTAGCgaggggatatcaatctacacatcagataattacacggaagagatgggttccggcgattacgactctatgaaagagccgtgttttagagaggaaaacgcgaactttaacaagatctttctccccaccatctacagcatcatctttctcacaggcatcgtagggaacggcctggtcatcctggtgatgggataccaaaagaagctgaggtcaatgaccgacaagtataggctccatctgtccgtggccgacctcctgtttgtgattaccctgcctttttgggcagttgacgctgtcgctaattggtacttcggcaacttcctctgtaaggcagtgcacgttatctacactgtgaatctttatagttccgtcttgatcttggcctttatcagtctcgacaggtatttggcgattgtgcacgctaccaactcacaacgacctagaaaactcctggctgagaaggtggtttacgttggtgtgtggattccagctctcctgttgacaataccagacttcattttcgctaatgtgagcgaggccgatgacagatacatttgtgaccgattttacccaaacgatctgtgggtagtggtatttcagttccaacacattatggtcgggctgatcttgcccggcattgtcatactgtcttgctactgcatcattatttctaagctgtcacactcaaaaggccaccaaaagaggaaggctctgaaaacaacggtgatcctgatactggccttcttcgcatgttggctgccctactatatcggcatcagcattgactcatttatactcctggaaattatcaagcagggctgcgagttcgagaacaccgttcataagtggatttctataaccgaggccctcgccttctttcactgttgtttgaatccgattctctacgcgtttcttggcgccaaatttaaaacaagcgcccaacatgcactgacatcagtgtctagggggagctctctgaaaatcctctccaagggaaaacgaggcggacatagcagtgtcagcactgagtccgaatccagctcatttcatagctctatcgataccggtggacgcaccccacccagcctgggtccccaagatgagtcctgcaccaccgccagctcctccctggccaaggacacttcatcgaccggtgagaacctgtacttccagctaagattagataaaagtaaagtgattaacagcgcattagagctgcttaatgaggtcggaatcgaaggtttaacaacccgtaaactcgcccagaagctaggtgtagagcagcctacattgtattggcatgtaaaaaataagcgggctttgctcgacgccttagccattgagatgttagataggcaccatactcacttttgccctttagaaggggaaagctggcaagattttttacgtaataacgctaaaagttttagatgtgctttactaagtcatcgcgatggagcaaaagtacatttaggtacacggcctacagaaaaacagtatgaaactctcgaaaatcaattagcctttttatgccaacaaggtttttcactagagaatgcattatatgcactcagcgctgtggggcattttactttaggttgcgtattggaagatcaagagcatcaagtcgctaaagaagaaagggaaacacctactactgatagtatgccgccattattacgacaagctatcgaattatttgatcaccaaggtgcagagccagccttcttattcggccttgaattgatcatatgcggattagaaaaacaacttaaatgtgaaagtgggtccgcgtacagccgcgcgcgtacgaaaaacaattacgggtctaccatcgagggcctgctcgatctcccggacgacgacgcccccgaagaggcggggctggcggctccgcgcctgtcctttctccccgcgggacacacgcgcagactgtcgacggcccccccgaccgatgtcagcctgggggacgagctccacttagacggcgaggacgtggcgatggcgcatgccgacgcgctagacgatttcgatctggacatgttgggggacggggattccccgggtccgggatttaccccccacgactccgccccctacggcgctctggatatggccgacttcgagtttgagcagatgtttaccgatgcccttggaattgacgagtacggtgggtagACGCGTCATATGGCTAGCCTGCAGGGATCCAATGTAACTGTATTCAGCGATGACGAAATTCTTAGCTATTGTAATACTCTAGAGGATCTTTGTGAAGGAACCTTACTTCTGTGGTGTGACATAATTGGACAAACTACCTACAGAGATTTAAAGCTCTAAGGTAAATATAAAATTTTTAAGTGTATAATGTGTTAAACTACTGATTCTAATTGTTTGTGTATTTTAGATTCCAACCTATGGAACTGATGAATGGGAGCAGTGGTGGAATGCCTTTAATGAGGAAAACCTGTTTTGCTCAGAAGAAATGCCATCTAGTGATGATGAGGCTACTGCTGACTCTCAACATTCTACTCCTCCAAAAAAGAAGAGAAAGGTAGAAGACCCCAAGGACTTTCCTTCAGAATTGCTAAGTTTTTTGAGTCATGCTGTGTTTAGTAATAGAACTCTTGCTTGCTTTGCTATTTACACCACAAAGGAAAAAGCTGCACTGCTATACAAGAAAATTATGGAAAAATATTCTGTAACCTTTATAAGTAGGCATAACAGTTATAATCATAACATACTGTTTTTTCTTACTCCACACAGGCATAGAGTGTCTGCTATTAATAACTATGCTCAAAAATTGTGTACCTTTAGCTTTTTAATTTGTAAAGGGGTTAATAAGGAATATTTGATGTATAGTGCCTTGACTAGAGATCATAATCAGCCATACCACATTTGTAGAGGTTTTACTTGCTTTAAAAAACCTCCCACACCTCCCCCTGAACCTGAAACATAAAATGAATGCAATTGTTGTTGTTAACTTGTTTATTGCAGCTTATAATGGTTACAAATAAAGCAATAGCATCACAAATTTCACAAATAAAGCATTTTTTTCACTGCATTCTAGTTGTGGTTTGTCCAAACTCATCAATGTATCTTATCATGTCTGCGGCTCTAGAGCTGCATTAATGAATCGGCCAACGCGCGGGGAGAGGCGGTTTGCGTATTGGGCGCTCTTCCGCTTCCTCGCTCACTGACTCGCTGCGCTCGGTCGTTCGGCTGCGGCGAGCGGTATCAGCTCACTCAAAGGCGGTAATACGGTTATCCACAGAATCAGGGGATAACGCAGGAAAGAACATGTGAGCAAAAGGCCAGCAAAAGGCCAGGAACCGTAAAAAGGCCGCGTTGCTGGCGTTTTTCCATAGGCTCCGCCCCCCTGACGAGCATCACAAAAATCGACGCTCAAGTCAGAGGTGGCGAAACCCGACAGGACTATAAAGATACCAGGCGTTTCCCCCTGGAAGCTCCCTCGTGCGCTCTCCTGTTCCGACCCTGCCGCTTACCGGATACCTGTCCGCCTTTCTCCCTTCGGGAAGCGTGGCGCTTTCTCATAGCTCACGCTGTAGGTATCTCAGTTCGGTGTAGGTCGTTCGCTCCAAGCTGGGCTGTGTGCACGAACCCCCCGTTCAGCCCGACCGCTGCGCCTTATCCGGTAACTATCGTCTTGAGTCCAACCCGGTAAGACACGACTTATCGCCACTGGCAGCAGCCACTGGTAACAGGATTAGCAGAGCGAGGTATGTAGGCGGTGCTACAGAGTTCTTGAAGTGGTGGCCTAACTACGGCTACACTAGAAGAACAGTATTTGGTATCTGCGCTCTGCTGAAGCCAGTTACCTTCGGAAAAAGAGTTGGTAGCTCTTGATCCGGCAAACAAACCACCGCTGGTAGCGGTGGTTTTTTTGTTTGCAAGCAGCAGATTACGCGCAGAAAAAAAGGATCTCAAGAAGATCCTTTGATCTTTTCTACGGGGTCTGACGCTCAGTGGAACGAAAACTCACGTTAAGGGATTTTGGTCATGAGATTATCAAAAAGGATCTTCACCTAGATCCTTTTAAATTAAAAATGAAGTTTTAAATCAATCTAAAGTATATATGAGTAACCTGAGGCTATGGCAGGGCCTGCCGCCCCGACGTTGGCTGCGAGCCCTGGGCCTTCACCCGAACTTGGGGGGTGGGGTGGGGAAAAGGAAGAAACGCGGGCGTATTGGCCCCAATGGGGTCTCGGTGGGGTATCGACAGAGTGCCAGCCCTGGGACCGAACCCCGCGTTTATGAACAAACGACCCAACACCGTGCGTTTTATTCTGTCTTTTTATTGCCGTCATAGCGCGGGTTCCTTCCGGTATTGTCTCCTTCCGTGTTTCAGTTAGCCTCCCCCTAGGGTGGGCGAAGAACTCCAGCATGAGATCCCCGCGCTGGAGGATCATCCAGCCGGCGTCCCGGAAAACGATTCCGAAGCCCAACCTTTCATAGAAGGCGGCGGTGGAATCGAAATCTCGTGATGGCAGGTTGGGCGTCGCTTGGTCGGTCATTTCGAACCCCAGAGTCCCGCTCAGAAGAACTCGTCAAGAAGGCGATAGAAGGCGATGCGCTGCGAATCGGGAGCGGCGATACCGTAAAGCACGAGGAAGCGGTCAGCCCATTCGCCGCCAAGCTCTTCAGCAATATCACGGGTAGCCAACGCTATGTCCTGATAGCGGTCCGCCACACCCAGCCGGCCACAGTCGATGAATCCAGAAAAGCGGCCATTTTCCACCATGATATTCGGCAAGCAGGCATCGCCATGGGTCACGACGAGATCCTCGCCGTCGGGCATGCTCGCCTTGAGCCTGGCGAACAGTTCGGCTGGCGCGAGCCCCTGATGCTCTTCGTCCAGATCATCCTGATCGACAAGACCGGCTTCCATCCGAGTACGTGCTCGCTCGATGCGATGTTTCGCTTGGTGGTCGAATGGGCAGGTAGCCGGATCAAGCGTATGCAGCCGCCGCATTGCATCAGCCATGATGGATACTTTCTCGGCAGGAGCAAGGTGAGATGACAGGAGATCCTGCCCCGGCACTTCGCCCAATAGCAGCCAGTCCCTTCCCGCTTCAGTGACAACGTCGAGCACAGCTGCGCAAGGAACGCCCGTCGTGGCCAGCCACGATAGCCGCGCTGCCTCGTCTTGCAGTTCATTCAGGGCACCGGACAGGTCGGTCTTGACAAAAAGAACCGGGCGCCCCTGCGCTGACAGCCGGAACACGGCGGCATCAGAGCAGCCGATTGTCTGTTGTGCCCAGTCATAGCCGAATAGCCTCTCCACCCAAGCGGCCGGAGAACCTGCGTGCAATCCATCTTGTTCAATCATGCGAAACGATCCTCATCCTGTCTCTTGATCGATCTTTGCAAAAGCCTAGGCCTCCAAAAAAGCCTCCTCACTACTTCTGGAATAGCTCAGAGGCCGAGGCGGCCTCGGCCTCTGCATAAATAAAAAAAATTAGTCAGCCATGGGGCGGAGAATGGGCGGAACTGGGCGGAGTTAGGGGCGGGATGGGCGGAGTTAGGGGCGGGACTATGGTTGCTGACTAATTGAGATGCATGCTTTGCATACTTCTGCCTGCTGGGGAGCCTGGGGACTTTCCACACCTGGTTGCTGACTAATTGAGATGCATGCTTTGCATACTTCTGCCTGCTGGGGAGCCTGGGGACTTTCCACACCCTAACTGACACACATTCCACAGCTGGTTCTTTCCGCCTCAGGACTCTTCCTTTTTCAATATTATTGAAGCATTTATCAGGGTTATTGTCTCATGAGCGGATACATATTTGAATGTATTTAGAAAAATAAACAAATAGGGGTTCCGCGCACATTTCCCCGAAAAGTGCCACCTGACGTCTAAGAAACCATTATTATCATGACATTAACCTATAAAAATAGGCGTATCACGAGGCCCTTTCGTCTTCA

**Nucleic acid Sequence of the construct PA-CXCR4-TEVcs-tTA**

GACGGATCGGGAGATCTCCCGATCCCCTATGGTGCACTCTCAGTACAATCTGCTCTGATGCCGCATAGTTAAGCCAGTATCTGCTCCCTGCTTGTGTGTTGGAGGTCGCTGAGTAGTGCGCGAGCAAAATTTAAGCTACAACAAGGCAAGGCTTGACCGACAATTGCATGAAGAATCTGCTTAGGGTTAGGCGTTTTGCGCTGCTTCGCGATGTACGGGCCAGATATACGCGTTGACATTGATTATTGACTAGTTATTAATAGTAATCAATTACGGGGTCATTAGTTCATAGCCCATATATGGAGTTCCGCGTTACATAACTTACGGTAAATGGCCCGCCTGGCTGACCGCCCAACGACCCCCGCCCATTGACGTCAATAATGACGTATGTTCCCATAGTAACGCCAATAGGGACTTTCCATTGACGTCAATGGGTGGAGTATTTACGGTAAACTGCCCACTTGGCAGTACATCAAGTGTATCATATGCCAAGTACGCCCCCTATTGACGTCAATGACGGTAAATGGCCCGCCTGGCATTATGCCCAGTACATGACCTTATGGGACTTTCCTACTTGGCAGTACATCTACGTATTAGTCATCGCTATTACCATGGTGATGCGGTTTTGGCAGTACATCAATGGGCGTGGATAGCGGTTTGACTCACGGGGATTTCCAAGTCTCCACCCCATTGACGTCAATGGGAGTTTGTTTTGGCACCAAAATCAACGGGACTTTCCAAAATGTCGTAACAACTCCGCCCCATTGACGCAAATGGGCGGTAGGCGTGTACGGTGGGAGGTCTATATAAGCAGAGCTCTCTGGCTAACTAGAGAACCCACTGCTTACTGGCTTATCGAAATTAATACGACTCACTATAGGGAGACCCAAGCTGGCTAGCGTTTAAACTTAAGCTTGGTACCGAGCTCGGATCCACTAGTCCAGTGTGGTGGAATTCTGCAGATATCCAGCACAGTGGCGGCCGCGCCACCATGAACGGTACCGAAGGCCCAAACTTCTACGTTCCTTTCTCCAACAAGACGGGCGTGGTGCGCAGCCCGTTCGAGGCTCCGCAGTACTACCTGGCGGAGCCCTGGCAGTTCTCCATGCTGGCCGCCTACATGTTCCTGCTGATCATGCTTGGCTTCCCGATCAACTTCCTCACGCTGTACGTCATGGGTTACCAGAAGAAACTGAGAAGCATGCTCAACTACATCCTGCTCAACCTGGCCGTGGCAGATCTCTTCATGGTCTTCGGTGGCTTCACCACCACCCTCTACACCTCTCTCCATGGGTACTTCGTCTTTGGGCCGACGGGCTGCAACCTCGAGGGCTTCTTTGCCACCCTGGGCGGTGAAATTGCACTGTGGTCTCTGGTAGTACTGGCGATCGAGCGGTACGTGGTGGTGGTCCACGCCACCAACAGTCAGAGGCCAAGGAAGCTGTTGGCCATCATGGGCGTCGCCTTCACCTGGGTCATGGCTCTGGCCTGTGCGGCCCCGCCGCTCGTCGGCTGGTCTAGATACATCCCGGAGGGCATGCAGTGCTCGTGCGGGATCGATTACTACACGCCGCACGAGGAGACCAACAATGAGTCGTTCGTCATCTACATGTTCGTGGTCCACTTCATCATCCCGCTGATTGTCATCTTCTTCTGCTATGGCATTATCATCTCCAAGCTGTCACACTCCAAGGGCCACCAGAAGCGCAAGGCCCTCCGTATGGTTATCATCATGGTCATCGCTTTCCTAATCTGCTGGCTGCCATATGCTGGTGTGGCGTTCTACATCTTCACCCATCAGGGCTCTGACTTTGGGCCCATCTTCATGACCATCCCGGCTTTCTTTGCCAAGACGTCTGCCGTCTACAACCCGGTCATCTACATCATGATGAACAAGCAGTTCCGGACCTCTGCCCAGCACGCACTCACCTCTGTGAGCAGAGGGTCCAGCCTCAAGATCCTCTCCAAAGGAAAGCGAGGTGGACATTCATCTGTTTCCACTGAGTCTGAGTCTTCAAGTTTTCACTCCAGCATCGATACCGGTGGACGCACCCCACCCAGCCTGGGTCCCCAAGATGAGTCCTGCACCACCGCCAGCTCCTCCCTGGCCAAGGACACTTCATCGACCGGTGAGAACCTGTACTTCCAGCTAAGATTAGATAAAAGTAAAGTGATTAACAGCGCATTAGAGCTGCTTAATGAGGTCGGAATCGAAGGTTTAACAACCCGTAAACTCGCCCAGAAGCTAGGTGTAGAGCAGCCTACATTGTATTGGCATGTAAAAAATAAGCGGGCTTTGCTCGACGCCTTAGCCATTGAGATGTTAGATAGGCACCATACTCACTTTTGCCCTTTAGAAGGGGAAAGCTGGCAAGATTTTTTACGTAATAACGCTAAAAGTTTTAGATGTGCTTTACTAAGTCATCGCGATGGAGCAAAAGTACATTTAGGTACACGGCCTACAGAAAAACAGTATGAAACTCTCGAAAATCAATTAGCCTTTTTATGCCAACAAGGTTTTTCACTAGAGAATGCATTATATGCACTCAGCGCTGTGGGGCATTTTACTTTAGGTTGCGTATTGGAAGATCAAGAGCATCAAGTCGCTAAAGAAGAAAGGGAAACACCTACTACTGATAGTATGCCGCCATTATTACGACAAGCTATCGAATTATTTGATCACCAAGGTGCAGAGCCAGCCTTCTTATTCGGCCTTGAATTGATCATATGCGGATTAGAAAAACAACTTAAATGTGAAAGTGGGTCCGCGTACAGCCGCGCGCGTACGAAAAACAATTACGGGTCTACCATCGAGGGCCTGCTCGATCTCCCGGACGACGACGCCCCCGAAGAGGCGGGGCTGGCGGCTCCGCGCCTGTCCTTTCTCCCCGCGGGACACACGCGCAGACTGTCGACGGCCCCCCCGACCGATGTCAGCCTGGGGGACGAGCTCCACTTAGACGGCGAGGACGTGGCGATGGCGCATGCCGACGCGCTAGACGATTTCGATCTGGACATGTTGGGGGACGGGGATTCCCCGGGTCCGGGATTTACCCCCCACGACTCCGCCCCCTACGGCGCTCTGGATATGGCCGACTTCGAGTTTGAGCAGATGTTTACCGATGCCCTTGGAATTGACGAGTACGGTGGG TCTAGAGGGCCCGTTTAAACCCGCTGATCAGCCTCGACTGTGCCTTCTAGTTGCCAGCCATCTGTTGTTTGCCCCTCCCCCGTGCCTTCCTTGACCCTGGAAGGTGCCACTCCCACTGTCCTTTCCTAATAAAATGAGGAAATTGCATCGCATTGTCTGAGTAGGTGTCATTCTATTCTGGGGGGTGGGGTGGGGCAGGACAGCAAGGGGGAGGATTGGGAAGACAATAGCAGGCATGCTGGGGATGCGGTGGGCTCTATGGCTTCTGAGGCGGAAAGAACCAGCTGGGGCTCTAGGGGGTATCCCCACGCGCCCTGTAGCGGCGCATTAAGCGCGGCGGGTGTGGTGGTTACGCGCAGCGTGACCGCTACACTTGCCAGCGCCCTAGCGCCCGCTCCTTTCGCTTTCTTCCCTTCCTTTCTCGCCACGTTCGCCGGCTTTCCCCGTCAAGCTCTAAATCGGGGGCTCCCTTTAGGGTTCCGATTTAGTGCTTTACGGCACCTCGACCCCAAAAAACTTGATTAGGGTGATGGTTCACGTAGTGGGCCATCGCCCTGATAGACGGTTTTTCGCCCTTTGACGTTGGAGTCCACGTTCTTTAATAGTGGACTCTTGTTCCAAACTGGAACAACACTCAACCCTATCTCGGTCTATTCTTTTGATTTATAAGGGATTTTGCCGATTTCGGCCTATTGGTTAAAAAATGAGCTGATTTAACAAAAATTTAACGCGAATTAATTCTGTGGAATGTGTGTCAGTTAGGGTGTGGAAAGTCCCCAGGCTCCCCAGCAGGCAGAAGTATGCAAAGCATGCATCTCAATTAGTCAGCAACCAGGTGTGGAAAGTCCCCAGGCTCCCCAGCAGGCAGAAGTATGCAAAGCATGCATCTCAATTAGTCAGCAACCATAGTCCCGCCCCTAACTCCGCCCATCCCGCCCCTAACTCCGCCCAGTTCCGCCCATTCTCCGCCCCATGGCTGACTAATTTTTTTTATTTATGCAGAGGCCGAGGCCGCCTCTGCCTCTGAGCTATTCCAGAAGTAGTGAGGAGGCTTTTTTGGAGGCCTAGGCTTTTGCAAAAAGCTCCCGGGAGCTTGTATATCCATTTTCGGATCTGATCAAGAGACAGGATGAGGATCGTTTCGCATGATTGAACAAGATGGATTGCACGCAGGTTCTCCGGCCGCTTGGGTGGAGAGGCTATTCGGCTATGACTGGGCACAACAGACAATCGGCTGCTCTGATGCCGCCGTGTTCCGGCTGTCAGCGCAGGGGCGCCCGGTTCTTTTTGTCAAGACCGACCTGTCCGGTGCCCTGAATGAACTGCAGGACGAGGCAGCGCGGCTATCGTGGCTGGCCACGACGGGCGTTCCTTGCGCAGCTGTGCTCGACGTTGTCACTGAAGCGGGAAGGGACTGGCTGCTATTGGGCGAAGTGCCGGGGCAGGATCTCCTGTCATCTCACCTTGCTCCTGCCGAGAAAGTATCCATCATGGCTGATGCAATGCGGCGGCTGCATACGCTTGATCCGGCTACCTGCCCATTCGACCACCAAGCGAAACATCGCATCGAGCGAGCACGTACTCGGATGGAAGCCGGTCTTGTCGATCAGGATGATCTGGACGAAGAGCATCAGGGGCTCGCGCCAGCCGAACTGTTCGCCAGGCTCAAGGCGCGCATGCCCGACGGCGAGGATCTCGTCGTGACCCATGGCGATGCCTGCTTGCCGAATATCATGGTGGAAAATGGCCGCTTTTCTGGATTCATCGACTGTGGCCGGCTGGGTGTGGCGGACCGCTATCAGGACATAGCGTTGGCTACCCGTGATATTGCTGAAGAGCTTGGCGGCGAATGGGCTGACCGCTTCCTCGTGCTTTACGGTATCGCCGCTCCCGATTCGCAGCGCATCGCCTTCTATCGCCTTCTTGACGAGTTCTTCTGAGCGGGACTCTGGGGTTCGAAATGACCGACCAAGCGACGCCCAACCTGCCATCACGAGATTTCGATTCCACCGCCGCCTTCTATGAAAGGTTGGGCTTCGGAATCGTTTTCCGGGACGCCGGCTGGATGATCCTCCAGCGCGGGGATCTCATGCTGGAGTTCTTCGCCCACCCCAACTTGTTTATTGCAGCTTATAATGGTTACAAATAAAGCAATAGCATCACAAATTTCACAAATAAAGCATTTTTTTCACTGCATTCTAGTTGTGGTTTGTCCAAACTCATCAATGTATCTTATCATGTCTGTATACCGTCGACCTCTAGCTAGAGCTTGGCGTAATCATGGTCATAGCTGTTTCCTGTGTGAAATTGTTATCCGCTCACAATTCCACACAACATACGAGCCGGAAGCATAAAGTGTAAAGCCTGGGGTGCCTAATGAGTGAGCTAACTCACATTAATTGCGTTGCGCTCACTGCCCGCTTTCCAGTCGGGAAACCTGTCGTGCCAGCTGCATTAATGAATCGGCCAACGCGCGGGGAGAGGCGGTTTGCGTATTGGGCGCTCTTCCGCTTCCTCGCTCACTGACTCGCTGCGCTCGGTCGTTCGGCTGCGGCGAGCGGTATCAGCTCACTCAAAGGCGGTAATACGGTTATCCACAGAATCAGGGGATAACGCAGGAAAGAACATGTGAGCAAAAGGCCAGCAAAAGGCCAGGAACCGTAAAAAGGCCGCGTTGCTGGCGTTTTTCCATAGGCTCCGCCCCCCTGACGAGCATCACAAAAATCGACGCTCAAGTCAGAGGTGGCGAAACCCGACAGGACTATAAAGATACCAGGCGTTTCCCCCTGGAAGCTCCCTCGTGCGCTCTCCTGTTCCGACCCTGCCGCTTACCGGATACCTGTCCGCCTTTCTCCCTTCGGGAAGCGTGGCGCTTTCTCATAGCTCACGCTGTAGGTATCTCAGTTCGGTGTAGGTCGTTCGCTCCAAGCTGGGCTGTGTGCACGAACCCCCCGTTCAGCCCGACCGCTGCGCCTTATCCGGTAACTATCGTCTTGAGTCCAACCCGGTAAGACACGACTTATCGCCACTGGCAGCAGCCACTGGTAACAGGATTAGCAGAGCGAGGTATGTAGGCGGTGCTACAGAGTTCTTGAAGTGGTGGCCTAACTACGGCTACACTAGAAGAACAGTATTTGGTATCTGCGCTCTGCTGAAGCCAGTTACCTTCGGAAAAAGAGTTGGTAGCTCTTGATCCGGCAAACAAACCACCGCTGGTAGCGGTGGTTTTTTTGTTTGCAAGCAGCAGATTACGCGCAGAAAAAAAGGATCTCAAGAAGATCCTTTGATCTTTTCTACGGGGTCTGACGCTCAGTGGAACGAAAACTCACGTTAAGGGATTTTGGTCATGAGATTATCAAAAAGGATCTTCACCTAGATCCTTTTAAATTAAAAATGAAGTTTTAAATCAATCTAAAGTATATATGAGTAAACTTGGTCTGACAGTTACCAATGCTTAATCAGTGAGGCACCTATCTCAGCGATCTGTCTATTTCGTTCATCCATAGTTGCCTGACTCCCCGTCGTGTAGATAACTACGATACGGGAGGGCTTACCATCTGGCCCCAGTGCTGCAATGATACCGCGAGACCCACGCTCACCGGCTCCAGATTTATCAGCAATAAACCAGCCAGCCGGAAGGGCCGAGCGCAGAAGTGGTCCTGCAACTTTATCCGCCTCCATCCAGTCTATTAATTGTTGCCGGGAAGCTAGAGTAAGTAGTTCGCCAGTTAATAGTTTGCGCAACGTTGTTGCCATTGCTACAGGCATCGTGGTGTCACGCTCGTCGTTTGGTATGGCTTCATTCAGCTCCGGTTCCCAACGATCAAGGCGAGTTACATGATCCCCCATGTTGTGCAAAAAAGCGGTTAGCTCCTTCGGTCCTCCGATCGTTGTCAGAAGTAAGTTGGCCGCAGTGTTATCACTCATGGTTATGGCAGCACTGCATAATTCTCTTACTGTCATGCCATCCGTAAGATGCTTTTCTGTGACTGGTGAGTACTCAACCAAGTCATTCTGAGAATAGTGTATGCGGCGACCGAGTTGCTCTTGCCCGGCGTCAATACGGGATAATACCGCGCCACATAGCAGAACTTTAAAAGTGCTCATCATTGGAAAACGTTCTTCGGGGCGAAAACTCTCAAGGATCTTACCGCTGTTGAGATCCAGTTCGATGTAACCCACTCGTGCACCCAACTGATCTTCAGCATCTTTTACTTTCACCAGCGTTTCTGGGTGAGCAAAAACAGGAAGGCAAAATGCCGCAAAAAAGGGAATAAGGGCGACACGGAAATGTTGAATACTCATACTCTTCCTTTTTCAATATTATTGAAGCATTTATCAGGGTTATTGTCTCATGAGCGGATACATATTTGAATGTATTTAGAAAAATAAACAAATAGGGGTTCCGCGCACATTTCCCCGAAAAGTGCCACCTGACGTC

**Nucleic acid Sequence of the construct PA-CXCR4-TEVcs-FLAG- SNAIL-6SA**

GACGGATCGGGAGATCTCCCGATCCCCTATGGTGCACTCTCAGTACAATCTGCTCTGATGCCGCATAGTTAAGCCAGTATCTGCTCCCTGCTTGTGTGTTGGAGGTCGCTGAGTAGTGCGCGAGCAAAATTTAAGCTACAACAAGGCAAGGCTTGACCGACAATTGCATGAAGAATCTGCTTAGGGTTAGGCGTTTTGCGCTGCTTCGCGATGTACGGGCCAGATATACGCGTTGACATTGATTATTGACTAGTTATTAATAGTAATCAATTACGGGGTCATTAGTTCATAGCCCATATATGGAGTTCCGCGTTACATAACTTACGGTAAATGGCCCGCCTGGCTGACCGCCCAACGACCCCCGCCCATTGACGTCAATAATGACGTATGTTCCCATAGTAACGCCAATAGGGACTTTCCATTGACGTCAATGGGTGGAGTATTTACGGTAAACTGCCCACTTGGCAGTACATCAAGTGTATCATATGCCAAGTACGCCCCCTATTGACGTCAATGACGGTAAATGGCCCGCCTGGCATTATGCCCAGTACATGACCTTATGGGACTTTCCTACTTGGCAGTACATCTACGTATTAGTCATCGCTATTACCATGGTGATGCGGTTTTGGCAGTACATCAATGGGCGTGGATAGCGGTTTGACTCACGGGGATTTCCAAGTCTCCACCCCATTGACGTCAATGGGAGTTTGTTTTGGCACCAAAATCAACGGGACTTTCCAAAATGTCGTAACAACTCCGCCCCATTGACGCAAATGGGCGGTAGGCGTGTACGGTGGGAGGTCTATATAAGCAGAGCTCTCTGGCTAACTAGAGAACCCACTGCTTACTGGCTTATCGAAATTAATACGACTCACTATAGGGAGACCCAAGCTGGCTAGCGTTTAAACTTAAGCTTGGTACCGAGCTCGGATCCACTAGTCCAGTGTGGTGGAATTCTGCAGATATCCAGCACAGTGGCGGCCGCGCCACCCGCTTTCCTAATCTGCTGGCTGCCATATGCTGGTGTGGCGTTCTACATCTTCACCCATCAGGGCTCTGACTTTGGGCCCATCTTCATGACCATCCCGGCTTTCTTTGCCAAGACGTCTGCCGTCTACAACCCGGTCATCTACATCATGATGAACAAGCAGTTCCGGACCTCTGCCCAGCACGCACTCACCTCTGTGAGCAGAGGGTCCAGCCTCAAGATCCTCTCCAAAGGAAAGCGAGGTGGACATTCATCTGTTTCCACTGAGTCTGAGTCTTCAAGTTTTCACTCCAGCATCGATACCGGTGGACGCACCCCACCCAGCCTGGGTCCCCAAGATGAGTCCTGCACCACCGCCAGCTCCTCCCTGGCCAAGGACACTTCATCGACCGGTGAGAACCTGTACTTCCAGCTAatggattacaaggatgacgacgataagagcccgggcggatccccgcgctctttcctcgtcaggaagccctccgaccccaatcggaagcctaactacagcgagctgcaggactctaatccagagtttaccttccagcagccctacgaccaggcccacctgctggcagccatcccacctccggagatcctcaaccccaccgcctcgctgccaatgctcatctgggactctgtcctggcgccccaagcccagccaattgcctgggcctcccttcggctccaggagagtcccagggtggcagagctgacctccctgtcagatgaggacgctgggaaaggcgcccagccccccagcccacccgcaccggctcctgcgtccttctccgcgacgtcagtcgcttccttggaggccgaggcctatgctgccttcccaggcttgggccaagtgcccaagcagctggcccagctctctgaggccaaggatctccaggctcgaaaggccttcaactgcaaatactgcaacaaggaatacctcagcctgggtgccctcaagatgcacatccgaagccacacgctgccctgcgtctgcggaacctgcgggaaggccttctctaggccctggctgctacaaggccatgtccggacccacactggcgagaagcccttctcctgtccccactgcagccgtgccttcgctgaccgctccaacctgcgggcccacctccagacccactcagatgtcaagaagtaccagtgccaggcgtgtgctcggaccttctcccgaatgtccctgctccacaagcaccaagagtccggctgctcaggatgtccccgctgaTCTAGAGGGCCCGTTTAAACCCGCTGATCAGCCTCGACTGTGCCTTCTAGTTGCCAGCCATCTGTTGTTTGCCCCTCCCCCGTGCCTTCCTTGACCCTGGAAGGTGCCACTCCCACTGTCCTTTCCTAATAAAATGAGGAAATTGCATCGCATTGTCTGAGTAGGTGTCATTCTATTCTGGGGGGTGGGGTGGGGCAGGACAGCAAGGGGGAGGATTGGGAAGACAATAGCAGGCATGCTGGGGATGCGGTGGGCTCTATGGCTTCTGAGGCGGAAAGAACCAGCTGGGGCTCTAGGGGGTATCCCCACGCGCCCTGTAGCGGCGCATTAAGCGCGGCGGGTGTGGTGGTTACGCGCAGCGTGACCGCTACACTTGCCAGCGCCCTAGCGCCCGCTCCTTTCGCTTTCTTCCCTTCCTTTCTCGCCACGTTCGCCGGCTTTCCCCGTCAAGCTCTAAATCGGGGGCTCCCTTTAGGGTTCCGATTTAGTGCTTTACGGCACCTCGACCCCAAAAAACTTGATTAGGGTGATGGTTCACGTAGTGGGCCATCGCCCTGATAGACGGTTTTTCGCCCTTTGACGTTGGAGTCCACGTTCTTTAATAGTGGACTCTTGTTCCAAACTGGAACAACACTCAACCCTATCTCGGTCTATTCTTTTGATTTATAAGGGATTTTGCCGATTTCGGCCTATTGGTTAAAAAATGAGCTGATTTAACAAAAATTTAACGCGAATTAATTCTGTGGAATGTGTGTCAGTTAGGGTGTGGAAAGTCCCCAGGCTCCCCAGCAGGCAGAAGTATGCAAAGCATGCATCTCAATTAGTCAGCAACCAGGTGTGGAAAGTCCCCAGGCTCCCCAGCAGGCAGAAGTATGCAAAGCATGCATCTCAATTAGTCAGCAACCATAGTCCCGCCCCTAACTCCGCCCATCCCGCCCCTAACTCCGCCCAGTTCCGCCCATTCTCCGCCCCATGGCTGACTAATTTTTTTTATTTATGCAGAGGCCGAGGCCGCCTCTGCCTCTGAGCTATTCCAGAAGTAGTGAGGAGGCTTTTTTGGAGGCCTAGGCTTTTGCAAAAAGCTCCCGGGAGCTTGTATATCCATTTTCGGATCTGATCAAGAGACAGGATGAGGATCGTTTCGCATGATTGAACAAGATGGATTGCACGCAGGTTCTCCGGCCGCTTGGGTGGAGAGGCTATTCGGCTATGACTGGGCACAACAGACAATCGGCTGCTCTGATGCCGCCGTGTTCCGGCTGTCAGCGCAGGGGCGCCCGGTTCTTTTTGTCAAGACCGACCTGTCCGGTGCCCTGAATGAACTGCAGGACGAGGCAGCGCGGCTATCGTGGCTGGCCACGACGGGCGTTCCTTGCGCAGCTGTGCTCGACGTTGTCACTGAAGCGGGAAGGGACTGGCTGCTATTGGGCGAAGTGCCGGGGCAGGATCTCCTGTCATCTCACCTTGCTCCTGCCGAGAAAGTATCCATCATGGCTGATGCAATGCGGCGGCTGCATACGCTTGATCCGGCTACCTGCCCATTCGACCACCAAGCGAAACATCGCATCGAGCGAGCACGTACTCGGATGGAAGCCGGTCTTGTCGATCAGGATGATCTGGACGAAGAGCATCAGGGGCTCGCGCCAGCCGAACTGTTCGCCAGGCTCAAGGCGCGCATGCCCGACGGCGAGGATCTCGTCGTGACCCATGGCGATGCCTGCTTGCCGAATATCATGGTGGAAAATGGCCGCTTTTCTGGATTCATCGACTGTGGCCGGCTGGGTGTGGCGGACCGCTATCAGGACATAGCGTTGGCTACCCGTGATATTGCTGAAGAGCTTGGCGGCGAATGGGCTGACCGCTTCCTCGTGCTTTACGGTATCGCCGCTCCCGATTCGCAGCGCATCGCCTTCTATCGCCTTCTTGACGAGTTCTTCTGAGCGGGACTCTGGGGTTCGAAATGACCGACCAAGCGACGCCCAACCTGCCATCACGAGATTTCGATTCCACCGCCGCCTTCTATGAAAGGTTGGGCTTCGGAATCGTTTTCCGGGACGCCGGCTGGATGATCCTCCAGCGCGGGGATCTCATGCTGGAGTTCTTCGCCCACCCCAACTTGTTTATTGCAGCTTATAATGGTTACAAATAAAGCAATAGCATCACAAATTTCACAAATAAAGCATTTTTTTCACTGCATTCTAGTTGTGGTTTGTCCAAACTCATCAATGTATCTTATCATGTCTGTATACCGTCGACCTCTAGCTAGAGCTTGGCGTAATCATGGTCATAGCTGTTTCCTGTGTGAAATTGTTATCCGCTCACAATTCCACACAACATACGAGCCGGAAGCATAAAGTGTAAAGCCTGGGGTGCCTAATGAGTGAGCTAACTCACATTAATTGCGTTGCGCTCACTGCCCGCTTTCCAGTCGGGAAACCTGTCGTGCCAGCTGCATTAATGAATCGGCCAACGCGCGGGGAGAGGCGGTTTGCGTATTGGGCGCTCTTCCGCTTCCTCGCTCACTGACTCGCTGCGCTCGGTCGTTCGGCTGCGGCGAGCGGTATCAGCTCACTCAAAGGCGGTAATACGGTTATCCACAGAATCAGGGGATAACGCAGGAAAGAACATGTGAGCAAAAGGCCAGCAAAAGGCCAGGAACCGTAAAAAGGCCGCGTTGCTGGCGTTTTTCCATAGGCTCCGCCCCCCTGACGAGCATCACAAAAATCGACGCTCAAGTCAGAGGTGGCGAAACCCGACAGGACTATAAAGATACCAGGCGTTTCCCCCTGGAAGCTCCCTCGTGCGCTCTCCTGTTCCGACCCTGCCGCTTACCGGATACCTGTCCGCCTTTCTCCCTTCGGGAAGCGTGGCGCTTTCTCATAGCTCACGCTGTAGGTATCTCAGTTCGGTGTAGGTCGTTCGCTCCAAGCTGGGCTGTGTGCACGAACCCCCCGTTCAGCCCGACCGCTGCGCCTTATCCGGTAACTATCGTCTTGAGTCCAACCCGGTAAGACACGACTTATCGCCACTGGCAGCAGCCACTGGTAACAGGATTAGCAGAGCGAGGTATGTAGGCGGTGCTACAGAGTTCTTGAAGTGGTGGCCTAACTACGGCTACACTAGAAGAACAGTATTTGGTATCTGCGCTCTGCTGAAGCCAGTTACCTTCGGAAAAAGAGTTGGTAGCTCTTGATCCGGCAAACAAACCACCGCTGGTAGCGGTGGTTTTTTTGTTTGCAAGCAGCAGATTACGCGCAGAAAAAAAGGATCTCAAGAAGATCCTTTGATCTTTTCTACGGGGTCTGACGCTCAGTGGAACGAAAACTCACGTTAAGGGATTTTGGTCATGAGATTATCAAAAAGGATCTTCACCTAGATCCTTTTAAATTAAAAATGAAGTTTTAAATCAATCTAAAGTATATATGAGTAAACTTGGTCTGACAGTTACCAATGCTTAATCAGTGAGGCACCTATCTCAGCGATCTGTCTATTTCGTTCATCCATAGTTGCCTGACTCCCCGTCGTGTAGATAACTACGATACGGGAGGGCTTACCATCTGGCCCCAGTGCTGCAATGATACCGCGAGACCCACGCTCACCGGCTCCAGATTTATCAGCAATAAACCAGCCAGCCGGAAGGGCCGAGCGCAGAAGTGGTCCTGCAACTTTATCCGCCTCCATCCAGTCTATTAATTGTTGCCGGGAAGCTAGAGTAAGTAGTTCGCCAGTTAATAGTTTGCGCAACGTTGTTGCCATTGCTACAGGCATCGTGGTGTCACGCTCGTCGTTTGGTATGGCTTCATTCAGCTCCGGTTCCCAACGATCAAGGCGAGTTACATGATCCCCCATGTTGTGCAAAAAAGCGGTTAGCTCCTTCGGTCCTCCGATCGTTGTCAGAAGTAAGTTGGCCGCAGTGTTATCACTCATGGTTATGGCAGCACTGCATAATTCTCTTACTGTCATGCCATCCGTAAGATGCTTTTCTGTGACTGGTGAGTACTCAACCAAGTCATTCTGAGAATAGTGTATGCGGCGACCGAGTTGCTCTTGCCCGGCGTCAATACGGGATAATACCGCGCCACATAGCAGAACTTTAAAAGTGCTCATCATTGGAAAACGTTCTTCGGGGCGAAAACTCTCAAGGATCTTACCGCTGTTGAGATCCAGTTCGATGTAACCCACTCGTGCACCCAACTGATCTTCAGCATCTTTTACTTTCACCAGCGTTTCTGGGTGAGCAAAAACAGGAAGGCAAAATGCCGCAAAAAAGGGAATAAGGGCGACACGGAAATGTTGAATACTCATACTCTTCCTTTTTCAATATTATTGAAGCATTTATCAGGGTTATTGTCTCATGAGCGGATACATATTTGAATGTATTTAGAAAAATAAACAAATAGGGGTTCCGCGCACATTTCCCCGAAAAGTGCCACCTGACGTC

**Nucleic acid Sequence of the construct PA-CXCR4-TEVcs-3xFLAG-NICD**

GACGGATCGGGAGATCTCCCGATCCCCTATGGTGCACTCTCAGTACAATCTGCTCTGATGCCGCATAGTTAAGCCAGTATCTGCTCCCTGCTTGTGTGTTGGAGGTCGCTGAGTAGTGCGCGAGCAAAATTTAAGCTACAACAAGGCAAGGCTTGACCGACAATTGCATGAAGAATCTGCTTAGGGTTAGGCGTTTTGCGCTGCTTCGCGATGTACGGGCCAGATATACGCGTTGACATTGATTATTGACTAGTTATTAATAGTAATCAATTACGGGGTCATTAGTTCATAGCCCATATATGGAGTTCCGCGTTACATAACTTACGGTAAATGGCCCGCCTGGCTGACCGCCCAACGACCCCCGCCCATTGACGTCAATAATGACGTATGTTCCCATAGTAACGCCAATAGGGACTTTCCATTGACGTCAATGGGTGGAGTATTTACGGTAAACTGCCCACTTGGCAGTACATCAAGTGTATCATATGCCAAGTACGCCCCCTATTGACGTCAATGACGGTAAATGGCCCGCCTGGCATTATGCCCAGTACATGACCTTATGGGACTTTCCTACTTGGCAGTACATCTACGTATTAGTCATCGCTATTACCATGGTGATGCGGTTTTGGCAGTACATCAATGGGCGTGGATAGCGGTTTGACTCACGGGGATTTCCAAGTCTCCACCCCATTGACGTCAATGGGAGTTTGTTTTGGCACCAAAATCAACGGGACTTTCCAAAATGTCGTAACAACTCCGCCCCATTGACGCAAATGGGCGGTAGGCGTGTACGGTGGGAGGTCTATATAAGCAGAGCTCTCTGGCTAACTAGAGAACCCACTGCTTACTGGCTTATCGAAATTAATACGACTCACTATAGGGAGACCCAAGCTGGCTAGCGTTTAAACTTAAGCTTGGTACCGAGCTCGGATCCACTAGTCCAGTGTGGTGGAATTCTGCAGATATCCAGCACAGTGGCGGCCGCGCCACC ATGAACGGTACCGAAGGCCCAAACTTCTACGTTCCTTTCTCCAACAAGACGGGCGTGGTGCGCAGCCCGTTCGAGGCTCCGCAGTACTACCTGGCGGAGCCCTGGCAGTTCTCCATGCTGGCCGCCTACATGTTCCTGCTGATCATGCTTGGCTTCCCGATCAACTTCCTCACGCTGTACGTCATGGGTTACCAGAAGAAACTGAGAAGCATGCTCAACTACATCCTGCTCAACCTGGCCGTGGCAGATCTCTTCATGGTCTTCGGTGGCTTCACCACCACCCTCTACACCTCTCTCCATGGGTACTTCGTCTTTGGGCCGACGGGCTGCAACCTCGAGGGCTTCTTTGCCACCCTGGGCGGTGAAATTGCACTGTGGTCTCTGGTAGTACTGGCGATCGAGCGGTACGTGGTGGTGGTCCACGCCACCAACAGTCAGAGGCCAAGGAAGCTGTTGGCCATCATGGGCGTCGCCTTCACCTGGGTCATGGCTCTGGCCTGTGCGGCCCCGCCGCTCGTCGGCTGGTCTAGATACATCCCGGAGGGCATGCAGTGCTCGTGCGGGATCGATTACTACACGCCGCACGAGGAGACCAACAATGAGTCGTTCGTCATCTACATGTTCGTGGTCCACTTCATCATCCCGCTGATTGTCATCTTCTTCTGCTATGGCATTATCATCTCCAAGCTGTCACACTCCAAGGGCCACCAGAAGCGCAAGGCCCTCCGTATGGTTATCATCATGGTCATCGCTTTCCTAATCTGCTGGCTGCCATATGCTGGTGTGGCGTTCTACATCTTCACCCATCAGGGCTCTGACTTTGGGCCCATCTTCATGACCATCCCGGCTTTCTTTGCCAAGACGTCTGCCGTCTACAACCCGGTCATCTACATCATGATGAACAAGCAGTTCCGGACCTCTGCCCAGCACGCACTCACCTCTGTGAGCAGAGGGTCCAGCCTCAAGATCCTCTCCAAAGGAAAGCGAGGTGGACATTCATCTGTTTCCACTGAGTCTGAGTCTTCAAGTTTTCACTCCAGCTTCCACTGAGTCTGAGTCTTCAAGTTTTCACTCCAGCATCGATACCGGTGGACGCACCCCACCCAGCCTGGGTCCCCAAGATGAGTCCTGCACCACCGCCAGCTCCTCCCTGGCCAAGGACACTTCATCGACCGGTGAGAACCTGTACTTCCAGCTAGACTATAAGGACCACGACGGAGACTACAAGGATCATGATATTGATTACAAAGACGATGACGATAAGATGGCCgtgctgctgtcccgcaagcgccggcggcagcatggccagctctggttccctgagggcttcaaagtgtctgaggccagcaagaagaagcggcgggagcccctcggcgaggactccgtgggcctcaagcccctgaagaacgcttcagacggtgccctcatggacgacaaccagaatgagtggggggacgaggacctggagaccaagaagttccggttcgaggagcccgtggttctgcctgacctggacgaccagacagaccaccggcagtggactcagcagcacctggatgccgctgacctgcgcatgtctgccatggcccccacaccgccccagggtgaggttgacgccgactgcatggacgtcaatgtccgcgggcctgatggcttcaccccgctcatgatcgcctcctgcagcgggggcggcctggagacgggcaacagcgaggaagaggaggacgcgccggccgtcatctccgacttcatctaccagggcgccagcctgcacaaccagacagaccgcacgggcgagaccgccttgcacctggccgcccgctactcacgctctgatgccgccaagcgcctgctggaggccagcgcagatgccaacatccaggacaacatgggccgcaccccgctgcatgcggctgtgtctgccgacgcacaaggtgtcttccagatcctgatccggaaccgagccacagacctggatgcccgcatgcatgatggcacgacgccactgatcctggctgcccgcctggccgtggagggcatgctggaggacctcatcaactcacacgccgacgtcaacgccgtagatgacctgggcaagtccgccctgcactgggccgccgccgtgaacaatgtggatgccgcagttgtgctcctgaagaacggggctaacaaagatatgcagaacaacagggaggagacacccctgtttctggccgcccgggagggcagctacgagaccgccaaggtgctgctggaccactttgccaaccgggacatcacggatcatatggaccgcctgccgcgcgacatcgcacaggagcgcatgcatcacgacatcgtgaggctgctggacgagtacaacctggtgcgcagcccgcagctgcacggagccccgctggggggcacgcccaccctgtcgcccccgctctgctcgcccaacggctacctgggcagcctcaagcccggcgtgcagggcaagaaggtccgcaagcccagcagcaaaggcctggcctgtggaagcaaggaggccaaggacctcaaggcacggaggaagaagtcccaggacggcaagggctgcctgctggacagctccggcatgctctcgcccgtggactccctggagtcaccccatggctacctgtcagacgtggcctcgccgccactgctgccctccccgttccagcagtctccgtccgtgcccctcaaccacctgcctgggatgcccgacacccacctgggcatcgggcacctgaacgtggcggccaagcccgagatggcggcgctgggtgggggcggccggctggcctttgagactggcccacctcgtctctcccacctgcctgtggcctctggcaccagcaccgtcctgggctccagcagcggaggggccctgaatttcactgtgggcgggtccaccagtttgaatggtcaatgcgagtggctgtcccggctgcagagcggcatggtgccgaaccaatacaaccctctgcgggggagtgtggcaccaggccccctgagcacacaggccccctccctgcagcatggcatggtaggcccgctgcacagtagccttgctgccagcgccctgtcccagatgatgagctaccagggcctgcccagcacccggctggccacccagcctcacctggtgcagacccagcaggtgcagccacaaaacttacagatgcagcagcagaacctgcagccagcaaacatccagcagcagcaaagcctgcagccgccaccaccaccaccacagccgcaccttggcgtgagctcagcagccagcggccacctgggccggagcttcctgagtggagagccgagccaggcagacgtgcagccactgggccccagcagcctggcggtgcacactattctgccccaggagagccccgccctgcccacgtcgctgccatcctcgctggtcccacccgtgaccgcagcccagttcctgacgcccccctcgcagcacagctactcctcgcctgtggacaacacccccagccaccagctacaggtgcctgagcaccccttcctcaccccgtcccctgagtcccctgaccagtggtccagctcgtccccgcattccaacgtctccgactggtccgagggcgtctccagccctcccaccagcatgcagtcccagatcgcccgcattccggaggccttcaagTAGTCTAGAGGGCCCGTTTAAACCCGCTGATCAGCCTCGACTGTGCCTTCTAGTTGCCAGCCATCTGTTGTTTGCCCCTCCCCCGTGCCTTCCTTGACCCTGGAAGGTGCCACTCCCACTGTCCTTTCCTAATAAAATGAGGAAATTGCATCGCATTGTCTGAGTAGGTGTCATTCTATTCTGGGGGGTGGGGTGGGGCAGGACAGCAAGGGGGAGGATTGGGAAGACAATAGCAGGCATGCTGGGGATGCGGTGGGCTCTATGGCTTCTGAGGCGGAAAGAACCAGCTGGGGCTCTAGGGGGTATCCCCACGCGCCCTGTAGCGGCGCATTAAGCGCGGCGGGTGTGGTGGTTACGCGCAGCGTGACCGCTACACTTGCCAGCGCCCTAGCGCCCGCTCCTTTCGCTTTCTTCCCTTCCTTTCTCGCCACGTTCGCCGGCTTTCCCCGTCAAGCTCTAAATCGGGGGCTCCCTTTAGGGTTCCGATTTAGTGCTTTACGGCACCTCGACCCCAAAAAACTTGATTAGGGTGATGGTTCACGTAGTGGGCCATCGCCCTGATAGACGGTTTTTCGCCCTTTGACGTTGGAGTCCACGTTCTTTAATAGTGGACTCTTGTTCCAAACTGGAACAACACTCAACCCTATCTCGGTCTATTCTTTTGATTTATAAGGGATTTTGCCGATTTCGGCCTATTGGTTAAAAAATGAGCTGATTTAACAAAAATTTAACGCGAATTAATTCTGTGGAATGTGTGTCAGTTAGGGTGTGGAAAGTCCCCAGGCTCCCCAGCAGGCAGAAGTATGCAAAGCATGCATCTCAATTAGTCAGCAACCAGGTGTGGAAAGTCCCCAGGCTCCCCAGCAGGCAGAAGTATGCAAAGCATGCATCTCAATTAGTCAGCAACCATAGTCCCGCCCCTAACTCCGCCCATCCCGCCCCTAACTCCGCCCAGTTCCGCCCATTCTCCGCCCCATGGCTGACTAATTTTTTTTATTTATGCAGAGGCCGAGGCCGCCTCTGCCTCTGAGCTATTCCAGAAGTAGTGAGGAGGCTTTTTTGGAGGCCTAGGCTTTTGCAAAAAGCTCCCGGGAGCTTGTATATCCATTTTCGGATCTGATCAAGAGACAGGATGAGGATCGTTTCGCATGATTGAACAAGATGGATTGCACGCAGGTTCTCCGGCCGCTTGGGTGGAGAGGCTATTCGGCTATGACTGGGCACAACAGACAATCGGCTGCTCTGATGCCGCCGTGTTCCGGCTGTCAGCGCAGGGGCGCCCGGTTCTTTTTGTCAAGACCGACCTGTCCGGTGCCCTGAATGAACTGCAGGACGAGGCAGCGCGGCTATCGTGGCTGGCCACGACGGGCGTTCCTTGCGCAGCTGTGCTCGACGTTGTCACTGAAGCGGGAAGGGACTGGCTGCTATTGGGCGAAGTGCCGGGGCAGGATCTCCTGTCATCTCACCTTGCTCCTGCCGAGAAAGTATCCATCATGGCTGATGCAATGCGGCGGCTGCATACGCTTGATCCGGCTACCTGCCCATTCGACCACCAAGCGAAACATCGCATCGAGCGAGCACGTACTCGGATGGAAGCCGGTCTTGTCGATCAGGATGATCTGGACGAAGAGCATCAGGGGCTCGCGCCAGCCGAACTGTTCGCCAGGCTCAAGGCGCGCATGCCCGACGGCGAGGATCTCGTCGTGACCCATGGCGATGCCTGCTTGCCGAATATCATGGTGGAAAATGGCCGCTTTTCTGGATTCATCGACTGTGGCCGGCTGGGTGTGGCGGACCGCTATCAGGACATAGCGTTGGCTACCCGTGATATTGCTGAAGAGCTTGGCGGCGAATGGGCTGACCGCTTCCTCGTGCTTTACGGTATCGCCGCTCCCGATTCGCAGCGCATCGCCTTCTATCGCCTTCTTGACGAGTTCTTCTGAGCGGGACTCTGGGGTTCGAAATGACCGACCAAGCGACGCCCAACCTGCCATCACGAGATTTCGATTCCACCGCCGCCTTCTATGAAAGGTTGGGCTTCGGAATCGTTTTCCGGGACGCCGGCTGGATGATCCTCCAGCGCGGGGATCTCATGCTGGAGTTCTTCGCCCACCCCAACTTGTTTATTGCAGCTTATAATGGTTACAAATAAAGCAATAGCATCACAAATTTCACAAATAAAGCATTTTTTTCACTGCATTCTAGTTGTGGTTTGTCCAAACTCATCAATGTATCTTATCATGTCTGTATACCGTCGACCTCTAGCTAGAGCTTGGCGTAATCATGGTCATAGCTGTTTCCTGTGTGAAATTGTTATCCGCTCACAATTCCACACAACATACGAGCCGGAAGCATAAAGTGTAAAGCCTGGGGTGCCTAATGAGTGAGCTAACTCACATTAATTGCGTTGCGCTCACTGCCCGCTTTCCAGTCGGGAAACCTGTCGTGCCAGCTGCATTAATGAATCGGCCAACGCGCGGGGAGAGGCGGTTTGCGTATTGGGCGCTCTTCCGCTTCCTCGCTCACTGACTCGCTGCGCTCGGTCGTTCGGCTGCGGCGAGCGGTATCAGCTCACTCAAAGGCGGTAATACGGTTATCCACAGAATCAGGGGATAACGCAGGAAAGAACATGTGAGCAAAAGGCCAGCAAAAGGCCAGGAACCGTAAAAAGGCCGCGTTGCTGGCGTTTTTCCATAGGCTCCGCCCCCCTGACGAGCATCACAAAAATCGACGCTCAAGTCAGAGGTGGCGAAACCCGACAGGACTATAAAGATACCAGGCGTTTCCCCCTGGAAGCTCCCTCGTGCGCTCTCCTGTTCCGACCCTGCCGCTTACCGGATACCTGTCCGCCTTTCTCCCTTCGGGAAGCGTGGCGCTTTCTCATAGCTCACGCTGTAGGTATCTCAGTTCGGTGTAGGTCGTTCGCTCCAAGCTGGGCTGTGTGCACGAACCCCCCGTTCAGCCCGACCGCTGCGCCTTATCCGGTAACTATCGTCTTGAGTCCAACCCGGTAAGACACGACTTATCGCCACTGGCAGCAGCCACTGGTAACAGGATTAGCAGAGCGAGGTATGTAGGCGGTGCTACAGAGTTCTTGAAGTGGTGGCCTAACTACGGCTACACTAGAAGAACAGTATTTGGTATCTGCGCTCTGCTGAAGCCAGTTACCTTCGGAAAAAGAGTTGGTAGCTCTTGATCCGGCAAACAAACCACCGCTGGTAGCGGTGGTTTTTTTGTTTGCAAGCAGCAGATTACGCGCAGAAAAAAAGGATCTCAAGAAGATCCTTTGATCTTTTCTACGGGGTCTGACGCTCAGTGGAACGAAAACTCACGTTAAGGGATTTTGGTCATGAGATTATCAAAAAGGATCTTCACCTAGATCCTTTTAAATTAAAAATGAAGTTTTAAATCAATCTAAAGTATATATGAGTAAACTTGGTCTGACAGTTACCAATGCTTAATCAGTGAGGCACCTATCTCAGCGATCTGTCTATTTCGTTCATCCATAGTTGCCTGACTCCCCGTCGTGTAGATAACTACGATACGGGAGGGCTTACCATCTGGCCCCAGTGCTGCAATGATACCGCGAGACCCACGCTCACCGGCTCCAGATTTATCAGCAATAAACCAGCCAGCCGGAAGGGCCGAGCGCAGAAGTGGTCCTGCAACTTTATCCGCCTCCATCCAGTCTATTAATTGTTGCCGGGAAGCTAGAGTAAGTAGTTCGCCAGTTAATAGTTTGCGCAACGTTGTTGCCATTGCTACAGGCATCGTGGTGTCACGCTCGTCGTTTGGTATGGCTTCATTCAGCTCCGGTTCCCAACGATCAAGGCGAGTTACATGATCCCCCATGTTGTGCAAAAAAGCGGTTAGCTCCTTCGGTCCTCCGATCGTTGTCAGAAGTAAGTTGGCCGCAGTGTTATCACTCATGGTTATGGCAGCACTGCATAATTCTCTTACTGTCATGCCATCCGTAAGATGCTTTTCTGTGACTGGTGAGTACTCAACCAAGTCATTCTGAGAATAGTGTATGCGGCGACCGAGTTGCTCTTGCCCGGCGTCAATACGGGATAATACCGCGCCACATAGCAGAACTTTAAAAGTGCTCATCATTGGAAAACGTTCTTCGGGGCGAAAACTCTCAAGGATCTTACCGCTGTTGAGATCCAGTTCGATGTAACCCACTCGTGCACCCAACTGATCTTCAGCATCTTTTACTTTCACCAGCGTTTCTGGGTGAGCAAAAACAGGAAGGCAAAATGCCGCAAAAAAGGGAATAAGGGCGACACGGAAATGTTGAATACTCATACTCTTCCTTTTTCAATATTATTGAAGCATTTATCAGGGTTATTGTCTCATGAGCGGATACATATTTGAATGTATTTAGAAAAATAAACAAATAGGGGTTCCGCGCACATTTCCCCGAAAAGTGCCACCTGACGTC

**Nucleic acid Sequence of the construct PA-CXCR4-TEVcs-N90β-catenin**

ATGAACGGTACCGAAGGCCCAAACTTCTACGTTCCTTTCTCCAACAAGACGGGCGTGGTGCGCAGCCCGTTCGAGGCTCCGCAGTACTACCTGGCGGAGCCCTGGCAGTTCTCCATGCTGGCCGCCTACATGTTCCTGCTGATCATGCTTGGCTTCCCGATCAACTTCCTCACGCTGTACGTCATGGGTTACCAGAAGAAACTGAGAAGCATGCTCAACTACATCCTGCTCAACCTGGCCGTGGCAGATCTCTTCATGGTCTTCGGTGGCTTCACCACCACCCTCTACACCTCTCTCCATGGGTACTTCGTCTTTGGGCCGACGGGCTGCAACCTCGAGGGCTTCTTTGCCACCCTGGGCGGTGAAATTGCACTGTGGTCTCTGGTAGTACTGGCGATCGAGCGGTACGTGGTGGTGGTCCACGCCACCAACAGTCAGAGGCCAAGGAAGCTGTTGGCCATCATGGGCGTCGCCTTCACCTGGGTCATGGCTCTGGCCTGTGCGGCCCCGCCGCTCGTCGGCTGGTCTAGATACATCCCGGAGGGCATGCAGTGCTCGTGCGGGATCGATTACTACACGCCGCACGAGGAGACCAACAATGAGTCGTTCGTCATCTACATGTTCGTGGTCCACTTCATCATCCCGCTGATTGTCATCTTCTTCTGCTATGGCATTATCATCTCCAAGCTGTCACACTCCAAGGGCCACCAGAAGCGCAAGGCCCTCCGTATGGTTATCATCATGGTCATCGCTTTCCTAATCTGCTGGCTGCCATATGCTGGTGTGGCGTTCTACATCTTCACCCATCAGGGCTCTGACTTTGGGCCCATCTTCATGACCATCCCGGCTTTCTTTGCCAAGACGTCTGCCGTCTACAACCCGGTCATCTACATCATGATGAACAAGCAGTTCCGGACCTCTGCCCAGCACGCACTCACCTCTGTGAGCAGAGGGTCCAGCCTCAAGATCCTCTCCAAAGGAAAGCGAGGTGGACATTCATCTGTTTCCACTGAGTCTGAGTCTTCAAGTTTTCACTCCAGCTTATTAATAGTAATCAATTACGGGGTCATTAGTTCATAGCCCATATATGGAGTTCCGCGTTACATAACTTACGGTAAATGGCCCGCCTGGCTGACCGCCCAACGACCCCCGCCCATTGACGTCAATAATGACGTATGTTCCCATAGTAACGCCAATAGGGACTTTCCATTGACGTCAATGGGTGGAGTATTTACGGTAAACTGCCCACTTGGCAGTACATCAAGTGTATCATATGCCAAGTACGCCCCCTATTGACGTCAATGACGGTAAATGGCCCGCCTGGCATTATGCCCAGTACATGACCTTATGGGACTTTCCTACTTGGCAGTACATCTACGTATTAGTCATCGCTATTACCATGGTGATGCGGTTTTGGCAGTACATCAATGGGCGTGGATAGCGGTTTGACTCACGGGGATTTCCAAGTCTCCACCCCATTGACGTCAATGGGAGTTTGTTTTGGCACCAAAATCAACGGGACTTTCCAAAATGTCGTAACAACTCCGCCCCATTGACGCAAATGGGCGGTAGGCGTGTACGGTGGGAGGTCTATATAAGCAGAGCTCTCTGGCTAACTAGAGAACCCACTGCTTACTGGCTTATCGAAATTAATACGACTCACTATAGGGAGACCCAAGCTGGCTAGCGTTTAAACTTAAGCTTGGTACCGAGCTCGGATCCACCATGGGGGAGAAACCCGGGACCAGGGTCTTCAAGAAGTCGAGCCCTAACTGCAAGCTCACCGTGTACTTGGGCAAGCGGGACTTCGTAGATCACCTGGACAAAGTGGACCCTGTAGATGGCGTGGTGCTTGTGGACCCTGACTACCTGAAGGACCGCAAAGTGTTTGTGACCCTCACCTGCGCCTTCCGCTATGGCCGTGAAGACCTGGATGTGCTGGGCTTGTCCTTCCGCAAAGACCTGTTCATCGCCACCTACCAGGCCTTCCCCCCGGTGCCCAACCCMCCCCGGCCCCCCACCCGCCTGCAGGACCGGCTGCTGAGGAAGCTGGGCCAGCATGCCCACCCCTTCTTCTTCACCATACCCCAGAATCTTCCATGCTCCGTCACACTGCAGCCAGGCCCAGAGGATACAGGAAAGGCCTGCGGCGTAGACTTTGAGATTCGAGCCTTCTGTGCTAAATCACTAGAAGAGAAAAGCCACAAAAGGAACTCTGTGCGGCTGGTGATCCGAAAGGTGCAGTTCGCCCCGGAGAAACCCGGCCCCCAGCCTTCAGCCGAAACCACACGCCACTTCCTCATGTCTGACCGGTCCCTGCACCTCGAGGCTTCCCTGGACAAGGAGCTGTACTACCATGGGGAGCCCCTCAATGTAAATGTCCACGTCACCAACAACTCCACCAAGACCGTCAAGAAGATCAAAGTCTCTGTGAGACAGTACGCCGACATCTGCCTCTTCAGCACCGCCCAGTACAAGTGTCCTGTGGCTCAACTCGAACAAGATGACCAGGTATCTCCCAGCTCCACATTCTGTAAGGTGTACACCATAACCCCACTGCTCAGCGACAACCGGGAGAAGCGGGGTCTCGCCCTGGATGGGAAACTCAAGCACGAGGACACCAACCTGGCTTCCAGCACCATCGTGAAGGAGGGTGCCAACAAGGAGGTGCTGGGAATCCTGGTGTCCTACAGGGTCAAGGTGAAGCTGGTGGTGTCTCGAGGCGGGGATGTCTCTGTGGAGCTGCCTTTTGTTCTTATGCACCCCAAGCCCCACGACCACATCCCCCTCCCCAGACCCCAGTCAGCCGCTCCGGAGACAGATGTCCCTGTGGACACCAACCTCATTGAATTTGATACCAACTATGCCACAGATGATGACATTGTGTTTGAGGACTTTGCCCGGCTTCGGCTGAAGGGGATGAAGGATGACGACTATGATGATCAACTCTGCGGATCCGGTGAGAGTCTCTTCAAAGGGCCGCGCGATTATAACCCCATCTCATCCACAATTTGTCACCTCACAAATGAGTCCGATGGACATACAACAAGCCTTTACGGCATTGGCTTCGGTCCATTCATCATCACCAACAAGCATCTTTTCAGACGCAATAACGGCACCTTGCTGGTGCAGTCCCTCCACGGCGTCTTTAAAGTCAAGAACACCACGACCCTTCAGCAACACCTGATCGACGGAAGGGATATGATTATCATTAGAATGCCCAAAGATTTCCCGCCATTTCCTCAGAAACTCAAGTTTAGAGAGCCACAGAGGGAAGAAAGGATCTGCTTGGTGACAACAAATTTTCAAACTAAGTCTATGTCATCCATGGTATCAGATACCAGCTGTACGTTTCCTAGTTCAGACGGAATATTCTGGAAGCATTGGATACAGACAAAGGACGGCCAATGCGGTAGTCCACTGGTGTCTACTCGGGATGGTTTCATCGTCGGCATTCATAGCGCCAGCAACTTCACCAATACTAATAACTACTTCACCAGCGTGCCCAAGAATTTCATGGAGCTTCTGACAAACCAGGAAGCCCAGCAGTGGGTTTCTGGATGGCGCTTGAACGCTGACTCCGTTCTGTGGGGAGGCCATAAAGTGTTTATGAGTAAACCTGAAGAACCCTTTCAGCCGGTGAAAGAGGCCACTCAGCTTATGAATGAGCTGGTTTATAGCCAGTAGGAATTCTGCAGATATCCAGCACAGTGGCGGCCGCTCGAGTCTAGAGGGCCCGTTTAAACCCGCTGATCAGCCTCGACTGTGCCTTCTAGTTGCCAGCCATCTGTTGTTTGCCCCTCCCCCGTGCCTTCCTTGACCCTGGAAGGTGCCACTCCCACTGTCCTTTCCTAATAAAATGAGGAAATTGCATCGCATTGTCTGAGTAGGTGTCATTCTATTCTGGGGGGTGGGGTGGGGCAGGACAGCAAGGGGGAGGATTGGGAAGACAATAGCAGGCATGCTGGGGATGCGGTGGGCTCTATGGCTTCTGAGGCGGAAAGAACCAGCTGGGGCTCTAGGGGGTATCCCCACGCGCCCTGTAGCGGCGCATTAAGCGCGGCGGGTGTGGTGGTTACGCGCAGCGTGACCGCTACACTTGCCAGCGCCCTAGCGCCCGCTCCTTTCGCTTTCTTCCCTTCCTTTCTCGCCACGTTCGCCGGCTTTCCCCGTCAAGCTCTAAATCGGGGGCTCCCTTTAGGGTTCCGATTTAGTGCTTTACGGCACCTCGACCCCAAAAAACTTGATTAGGGTGATGGTTCACGTAGTGGGCCATCGCCCTGATAGACGGTTTTTCGCCCTTTGACGTTGGAGTCCACGTTCTTTAATAGTGGACTCTTGTTCCAAACTGGAACAACACTCAACCCTATCTCGGTCTATTCTTTTGATTTATAAGGGATTTTGCCGATTTCGGCCTATTGGTTAAAAAATGAGCTGATTTAACAAAAATTTAACGCGAATTAATTCTGTGGAATGTGTGTCAGTTAGGGTGTGGAAAGTCCCCAGGCTCCCCAGCAGGCAGAAGTATGCAAAGCATGCATCTCAATTAGTCAGCAACCAGGTGTGGAAAGTCCCCAGGCTCCCCAGCAGGCAGAAGTATGCAAAGCATGCATCTCAATTAGTCAGCAACCATAGTCCCGCCCCTAACTCCGCCCATCCCGCCCCTAACTCCGCCCAGTTCCGCCCATTCTCCGCCCCATGGCTGACTAATTTTTTTTATTTATGCAGAGGCCGAGGCCGCCTCTGCCTCTGAGCTATTCCAGAAGTAGTGAGGAGGCTTTTTTGGAGGCCTAGGCTTTTGCAAAAAGCTCCCGGGAGCTTGTATATCCATTTTCGGATCTGATCAAGAGACAGGATGAGGATCGTTTCGCATGATTGAACAAGATGGATTGCACGCAGGTTCTCCGGCCGCTTGGGTGGAGAGGCTATTCGGCTATGACTGGGCACAACAGACAATCGGCTGCTCTGATGCCGCCGTGTTCCGGCTGTCAGCGCAGGGGCGCCCGGTTCTTTTTGTCAAGACCGACCTGTCCGGTGCCCTGAATGAACTGCAGGACGAGGCAGCGCGGCTATCGTGGCTGGCCACGACGGGCGTTCCTTGCGCAGCTGTGCTCGACGTTGTCACTGAAGCGGGAAGGGACTGGCTGCTATTGGGCGAAGTGCCGGGGCAGGATCTCCTGTCATCTCACCTTGCTCCTGCCGAGAAAGTATCCATCATGGCTGATGCAATGCGGCGGCTGCATACGCTTGATCCGGCTACCTGCCCATTCGACCACCAAGCGAAACATCGCATCGAGCGAGCACGTACTCGGATGGAAGCCGGTCTTGTCGATCAGGATGATCTGGACGAAGAGCATCAGGGGCTCGCGCCAGCCGAACTGTTCGCCAGGCTCAAGGCGCGCATGCCCGACGGCGAGGATCTCGTCGTGACCCATGGCGATGCCTGCTTGCCGAATATCATGGTGGAAAATGGCCGCTTTTCTGGATTCATCGACTGTGGCCGGCTGGGTGTGGCGGACCGCTATCAGGACATAGCGTTGGCTACCCGTGATATTGCTGAAGAGCTTGGCGGCGAATGGGCTGACCGCTTCCTCGTGCTTTACGGTATCGCCGCTCCCGATTCGCAGCGCATCGCCTTCTATCGCCTTCTTGACGAGTTCTTCTGAGCGGGACTCTGGGGTTCGAAATGACCGACCAAGCGACGCCCAACCTGCCATCACGAGATTTCGATTCCACCGCCGCCTTCTATGAAAGGTTGGGCTTCGGAATCGTTTTCCGGGACGCCGGCTGGATGATCCTCCAGCGCGGGGATCTCATGCTGGAGTTCTTCGCCCACCCCAACTTGTTTATTGCAGCTTATAATGGTTACAAATAAAGCAATAGCATCACAAATTTCACAAATAAAGCATTTTTTTCACTGCATTCTAGTTGTGGTTTGTCCAAACTCATCAATGTATCTTATCATGTCTGTATACCGTCGACCTCTAGCTAGAGCTTGGCGTAATCATGGTCATAGCTGTTTCCTGTGTGAAATTGTTATCCGCTCACAATTCCACACAACATACGAGCCGGAAGCATAAAGTGTAAAGCCTGGGGTGCCTAATGAGTGAGCTAACTCACATTAATTGCGTTGCGCTCACTGCCCGCTTTCCAGTCGGGAAACCTGTCGTGCCAGCTGCATTAATGAATCGGCCAACGCGCGGGGAGAGGCGGTTTGCGTATTGGGCGCTCTTCCGCTTCCTCGCTCACTGACTCGCTGCGCTCGGTCGTTCGGCTGCGGCGAGCGGTATCAGCTCACTCAAAGGCGGTAATACGGTTATCCACAGAATCAGGGGATAACGCAGGAAAGAACATGTGAGCAAAAGGCCAGCAAAAGGCCAGGAACCGTAAAAAGGCCGCGTTGCTGGCGTTTTTCCATAGGCTCCGCCCCCCTGACGAGCATCACAAAAATCGACGCTCAAGTCAGAGGTGGCGAAACCCGACAGGACTATAAAGATACCAGGCGTTTCCCCCTGGAAGCTCCCTCGTGCGCTCTCCTGTTCCGACCCTGCCGCTTACCGGATACCTGTCCGCCTTTCTCCCTTCGGGAAGCGTGGCGCTTTCTCATAGCTCACGCTGTAGGTATCTCAGTTCGGTGTAGGTCGTTCGCTCCAAGCTGGGCTGTGTGCACGAACCCCCCGTTCAGCCCGACCGCTGCGCCTTATCCGGTAACTATCGTCTTGAGTCCAACCCGGTAAGACACGACTTATCGCCACTGGCAGCAGCCACTGGTAACAGGATTAGCAGAGCGAGGTATGTAGGCGGTGCTACAGAGTTCTTGAAGTGGTGGCCTAACTACGGCTACACTAGAAGAACAGTATTTGGTATCTGCGCTCTGCTGAAGCCAGTTACCTTCGGAAAAAGAGTTGGTAGCTCTTGATCCGGCAAACAAACCACCGCTGGTAGCGGTGGTTTTTTTGTTTGCAAGCAGCAGATTACGCGCAGAAAAAAAGGATCTCAAGAAGATCCTTTGATCTTTTCTACGGGGTCTGACGCTCAGTGGAACGAAAACTCACGTTAAGGGATTTTGGTCATGAGATTATCAAAAAGGATCTTCACCTAGATCCTTTTAAATTAAAAATGAAGTTTTAAATCAATCTAAAGTATATATGAGTAAACTTGGTCTGACAGTTACCAATGCTTAATCAGTGAGGCACCTATCTCAGCGATCTGTCTATTTCGTTCATCCATAGTTGCCTGACTCCCCGTCGTGTAGATAACTACGATACGGGAGGGCTTACCATCTGGCCCCAGTGCTGCAATGATACCGCGAGACCCACGCTCACCGGCTCCAGATTTATCAGCAATAAACCAGCCAGCCGGAAGGGCCGAGCGCAGAAGTGGTCCTGCAACTTTATCCGCCTCCATCCAGTCTATTAATTGTTGCCGGGAAGCTAGAGTAAGTAGTTCGCCAGTTAATAGTTTGCGCAACGTTGTTGCCATTGCTACAGGCATCGTGGTGTCACGCTCGTCGTTTGGTATGGCTTCATTCAGCTCCGGTTCCCAACGATCAAGGCGAGTTACATGATCCCCCATGTTGTGCAAAAAAGCGGTTAGCTCCTTCGGTCCTCCGATCGTTGTCAGAAGTAAGTTGGCCGCAGTGTTATCACTCATGGTTATGGCAGCACTGCATAATTCTCTTACTGTCATGCCATCCGTAAGATGCTTTTCTGTGACTGGTGAGTACTCAACCAAGTCATTCTGAGAATAGTGTATGCGGCGACCGAGTTGCTCTTGCCCGGCGTCAATACGGGATAATACCGCGCCACATAGCAGAACTTTAAAAGTGCTCATCATTGGAAAACGTTCTTCGGGGCGAAAACTCTCAAGGATCTTACCGCTGTTGAGATCCAGTTCGATGTAACCCACTCGTGCACCCAACTGATCTTCAGCATCTTTTACTTTCACCAGCGTTTCTGGGTGAGCAAAAACAGGAAGGCAAAATGCCGCAAAAAAGGGAATAAGGGCGACACGGAAATGTTGAATACTCATACTCTTCCTTTTTCAATATTATTGAAGCATTTATCAGGGTTATTGTCTCATGAGCGGATACATATTTGAATGTATTTAGAAAAATAAACAAATAGGGGTTCCGCGCACATTTCCCCGAAAAGTGCCACCTGACG

**Nucleic acid Sequence of the construct PA-CXCR4-TEVcs-SOX2**

GACGGATCGGGAGATCTCCCGATCCCCTATGGTGCACTCTCAGTACAATCTGCTCTGATGCCGCATAGTTAAGCCAGTATCTGCTCCCTGCTTGTGTGTTGGAGGTCGCTGAGTAGTGCGCGAGCAAAATTTAAGCTACAACAAGGCAAGGCTTGACCGACAATTGCATGAAGAATCTGCTTAGGGTTAGGCGTTTTGCGCTGCTTCGCGATGTACGGGCCAGATATACGCGTTGACATTGATTATTGACTAGTTATTAATAGTAATCAATTACGGGGTCATTAGTTCATAGCCCATATATGGAGTTCCGCGTTACATAACTTACGGTAAATGGCCCGCCTGGCTGACCGCCCAACGACCCCCGCCCATTGACGTCAATAATGACGTATGTTCCCATAGTAACGCCAATAGGGACTTTCCATTGACGTCAATGGGTGGAGTATTTACGGTAAACTGCCCACTTGGCAGTACATCAAGTGTATCATATGCCAAGTACGCCCCCTATTGACGTCAATGACGGTAAATGGCCCGCCTGGCATTATGCCCAGTACATGACCTTATGGGACTTTCCTACTTGGCAGTACATCTACGTATTAGTCATCGCTATTACCATGGTGATGCGGTTTTGGCAGTACATCAATGGGCGTGGATAGCGGTTTGACTCACGGGGATTTCCAAGTCTCCACCCCATTGACGTCAATGGGAGTTTGTTTTGGCACCAAAATCAACGGGACTTTCCAAAATGTCGTAACAACTCCGCCCCATTGACGCAAATGGGCGGTAGGCGTGTACGGTGGGAGGTCTATATAAGCAGAGCTCTCTGGCTAACTAGAGAACCCACTGCTTACTGGCTTATCGAAATTAATACGACTCACTATAGGGAGACCCAAGCTGGCTAGCGTTTAAACTTAAGCTTGGTACCGAGCTCGGATCCACTAGTCCAGTGTGGTGGAATTCTGCAGATATCCAGCACAGTGGCGGCCGCGCCACCCGCTTTCCTAATCTGCTGGCTGCCATATGCTGGTGTGGCGTTCTACATCTTCACCCATCAGGGCTCTGACTTTGGGCCCATCTTCATGACCATCCCGGCTTTCTTTGCCAAGACGTCTGCCGTCTACAACCCGGTCATCTACATCATGATGAACAAGCAGTTCCGGACCTCTGCCCAGCACGCACTCACCTCTGTGAGCAGAGGGTCCAGCCTCAAGATCCTCTCCAAAGGAAAGCGAGGTGGACATTCATCTGTTTCCACTGAGTCTGAGTCTTCAAGTTTTCACTCCAGCATCGATACCGGTGGACGCACCCCACCCAGCCTGGGTCCCCAAGATGAGTCCTGCACCACCGCCAGCTCCTCCCTGGCCAAGGACACTTCATCGACCGGTGAGAACCTGTACTTCCAGCTAATGTACAACATGATGGAGACGGAGCTGAAGCCGCCGGGCCCGCAGCAAACTTCGGGGGGCGGCGGCGGCAACTCCACCGCGGCGGCGGCCGGCGGCAACCAGAAAAACAGCCCGGACCGCGTCAAGCGGCCCATGAATGCCTTCATGGTGTGGTCCCGCGGGCAGCGGCGCAAGATGGCCCAGGAGAACCCCAAGATGCACAACTCGGAGATCAGCAAGCGCCTGGGCGCCGAGTGGAAACTTTTGTCGGAGACGGAGAAGCGGCCGTTCATCGACGAGGCTAAGCGGCTGCGAGCGCTGCACATGAAGGAGCACCCGGATTATAAATACCGGCCCCGGCGGAAAACCAAGACGCTCATGAAGAAGGATAAGTACACGCTGCCCGGCGGGCTGCTGGCCCCCGGCGGCAATAGCATGGCGAGCGGGGTCGGGGTGGGCGCCGGCCTGGGCGCGGGCGTGAACCAGCGCATGGACAGTTACGCGCACATGAACGGCTGGAGCAACGGCAGCTACAGCATGATGCAGGACCAGCTGGGCTACCCGCAGCACCCGGGCCTCAATGCGCACGGCGCAGCGCAGATGCAGCCCATGCACCGCTACGACGTGAGCGCCCTGCAGTACAACTCCATGACCAGCTCGCAGACCTACATGAACGGCTCGCCCACCTACAGCATGTCCTACTCGCAGCAGGGCACCCCTGGCATGGCTCTTGGCTCCATGGGTTCGGTGGTCAAGTCCGAGGCCAGCTCCAGCCCCCCTGTGGTTACCTCTTCCTCCCACTCCAGGGCGCCCTGCCAGGCCGGGGACCTCCGGGACATGATCAGCATGTATCTCCCCGGCGCCGAGGTGCCGGAACCCGCCGCCCCCAGCAGACTTCACATGTCCCAGCACTACCAGAGCGGCCCGGTGCCCGGCACGGCCATTAACGGCACACTGCCCCTCTCACACATGTAGTCTAGAGGGCCCGTTTAAACCCGCTGATCAGCCTCGACTGTGCCTTCTAGTTGCCAGCCATCTGTTGTTTGCCCCTCCCCCGTGCCTTCCTTGACCCTGGAAGGTGCCACTCCCACTGTCCTTTCCTAATAAAATGAGGAAATTGCATCGCATTGTCTGAGTAGGTGTCATTCTATTCTGGGGGGTGGGGTGGGGCAGGACAGCAAGGGGGAGGATTGGGAAGACAATAGCAGGCATGCTGGGGATGCGGTGGGCTCTATGGCTTCTGAGGCGGAAAGAACCAGCTGGGGCTCTAGGGGGTATCCCCACGCGCCCTGTAGCGGCGCATTAAGCGCGGCGGGTGTGGTGGTTACGCGCAGCGTGACCGCTACACTTGCCAGCGCCCTAGCGCCCGCTCCTTTCGCTTTCTTCCCTTCCTTTCTCGCCACGTTCGCCGGCTTTCCCCGTCAAGCTCTAAATCGGGGGCTCCCTTTAGGGTTCCGATTTAGTGCTTTACGGCACCTCGACCCCAAAAAACTTGATTAGGGTGATGGTTCACGTAGTGGGCCATCGCCCTGATAGACGGTTTTTCGCCCTTTGACGTTGGAGTCCACGTTCTTTAATAGTGGACTCTTGTTCCAAACTGGAACAACACTCAACCCTATCTCGGTCTATTCTTTTGATTTATAAGGGATTTTGCCGATTTCGGCCTATTGGTTAAAAAATGAGCTGATTTAACAAAAATTTAACGCGAATTAATTCTGTGGAATGTGTGTCAGTTAGGGTGTGGAAAGTCCCCAGGCTCCCCAGCAGGCAGAAGTATGCAAAGCATGCATCTCAATTAGTCAGCAACCAGGTGTGGAAAGTCCCCAGGCTCCCCAGCAGGCAGAAGTATGCAAAGCATGCATCTCAATTAGTCAGCAACCATAGTCCCGCCCCTAACTCCGCCCATCCCGCCCCTAACTCCGCCCAGTTCCGCCCATTCTCCGCCCCATGGCTGACTAATTTTTTTTATTTATGCAGAGGCCGAGGCCGCCTCTGCCTCTGAGCTATTCCAGAAGTAGTGAGGAGGCTTTTTTGGAGGCCTAGGCTTTTGCAAAAAGCTCCCGGGAGCTTGTATATCCATTTTCGGATCTGATCAAGAGACAGGATGAGGATCGTTTCGCATGATTGAACAAGATGGATTGCACGCAGGTTCTCCGGCCGCTTGGGTGGAGAGGCTATTCGGCTATGACTGGGCACAACAGACAATCGGCTGCTCTGATGCCGCCGTGTTCCGGCTGTCAGCGCAGGGGCGCCCGGTTCTTTTTGTCAAGACCGACCTGTCCGGTGCCCTGAATGAACTGCAGGACGAGGCAGCGCGGCTATCGTGGCTGGCCACGACGGGCGTTCCTTGCGCAGCTGTGCTCGACGTTGTCACTGAAGCGGGAAGGGACTGGCTGCTATTGGGCGAAGTGCCGGGGCAGGATCTCCTGTCATCTCACCTTGCTCCTGCCGAGAAAGTATCCATCATGGCTGATGCAATGCGGCGGCTGCATACGCTTGATCCGGCTACCTGCCCATTCGACCACCAAGCGAAACATCGCATCGAGCGAGCACGTACTCGGATGGAAGCCGGTCTTGTCGATCAGGATGATCTGGACGAAGAGCATCAGGGGCTCGCGCCAGCCGAACTGTTCGCCAGGCTCAAGGCGCGCATGCCCGACGGCGAGGATCTCGTCGTGACCCATGGCGATGCCTGCTTGCCGAATATCATGGTGGAAAATGGCCGCTTTTCTGGATTCATCGACTGTGGCCGGCTGGGTGTGGCGGACCGCTATCAGGACATAGCGTTGGCTACCCGTGATATTGCTGAAGAGCTTGGCGGCGAATGGGCTGACCGCTTCCTCGTGCTTTACGGTATCGCCGCTCCCGATTCGCAGCGCATCGCCTTCTATCGCCTTCTTGACGAGTTCTTCTGAGCGGGACTCTGGGGTTCGAAATGACCGACCAAGCGACGCCCAACCTGCCATCACGAGATTTCGATTCCACCGCCGCCTTCTATGAAAGGTTGGGCTTCGGAATCGTTTTCCGGGACGCCGGCTGGATGATCCTCCAGCGCGGGGATCTCATGCTGGAGTTCTTCGCCCACCCCAACTTGTTTATTGCAGCTTATAATGGTTACAAATAAAGCAATAGCATCACAAATTTCACAAATAAAGCATTTTTTTCACTGCATTCTAGTTGTGGTTTGTCCAAACTCATCAATGTATCTTATCATGTCTGTATACCGTCGACCTCTAGCTAGAGCTTGGCGTAATCATGGTCATAGCTGTTTCCTGTGTGAAATTGTTATCCGCTCACAATTCCACACAACATACGAGCCGGAAGCATAAAGTGTAAAGCCTGGGGTGCCTAATGAGTGAGCTAACTCACATTAATTGCGTTGCGCTCACTGCCCGCTTTCCAGTCGGGAAACCTGTCGTGCCAGCTGCATTAATGAATCGGCCAACGCGCGGGGAGAGGCGGTTTGCGTATTGGGCGCTCTTCCGCTTCCTCGCTCACTGACTCGCTGCGCTCGGTCGTTCGGCTGCGGCGAGCGGTATCAGCTCACTCAAAGGCGGTAATACGGTTATCCACAGAATCAGGGGATAACGCAGGAAAGAACATGTGAGCAAAAGGCCAGCAAAAGGCCAGGAACCGTAAAAAGGCCGCGTTGCTGGCGTTTTTCCATAGGCTCCGCCCCCCTGACGAGCATCACAAAAATCGACGCTCAAGTCAGAGGTGGCGAAACCCGACAGGACTATAAAGATACCAGGCGTTTCCCCCTGGAAGCTCCCTCGTGCGCTCTCCTGTTCCGACCCTGCCGCTTACCGGATACCTGTCCGCCTTTCTCCCTTCGGGAAGCGTGGCGCTTTCTCATAGCTCACGCTGTAGGTATCTCAGTTCGGTGTAGGTCGTTCGCTCCAAGCTGGGCTGTGTGCACGAACCCCCCGTTCAGCCCGACCGCTGCGCCTTATCCGGTAACTATCGTCTTGAGTCCAACCCGGTAAGACACGACTTATCGCCACTGGCAGCAGCCACTGGTAACAGGATTAGCAGAGCGAGGTATGTAGGCGGTGCTACAGAGTTCTTGAAGTGGTGGCCTAACTACGGCTACACTAGAAGAACAGTATTTGGTATCTGCGCTCTGCTGAAGCCAGTTACCTTCGGAAAAAGAGTTGGTAGCTCTTGATCCGGCAAACAAACCACCGCTGGTAGCGGTGGTTTTTTTGTTTGCAAGCAGCAGATTACGCGCAGAAAAAAAGGATCTCAAGAAGATCCTTTGATCTTTTCTACGGGGTCTGACGCTCAGTGGAACGAAAACTCACGTTAAGGGATTTTGGTCATGAGATTATCAAAAAGGATCTTCACCTAGATCCTTTTAAATTAAAAATGAAGTTTTAAATCAATCTAAAGTATATATGAGTAAACTTGGTCTGACAGTTACCAATGCTTAATCAGTGAGGCACCTATCTCAGCGATCTGTCTATTTCGTTCATCCATAGTTGCCTGACTCCCCGTCGTGTAGATAACTACGATACGGGAGGGCTTACCATCTGGCCCCAGTGCTGCAATGATACCGCGAGACCCACGCTCACCGGCTCCAGATTTATCAGCAATAAACCAGCCAGCCGGAAGGGCCGAGCGCAGAAGTGGTCCTGCAACTTTATCCGCCTCCATCCAGTCTATTAATTGTTGCCGGGAAGCTAGAGTAAGTAGTTCGCCAGTTAATAGTTTGCGCAACGTTGTTGCCATTGCTACAGGCATCGTGGTGTCACGCTCGTCGTTTGGTATGGCTTCATTCAGCTCCGGTTCCCAACGATCAAGGCGAGTTACATGATCCCCCATGTTGTGCAAAAAAGCGGTTAGCTCCTTCGGTCCTCCGATCGTTGTCAGAAGTAAGTTGGCCGCAGTGTTATCACTCATGGTTATGGCAGCACTGCATAATTCTCTTACTGTCATGCCATCCGTAAGATGCTTTTCTGTGACTGGTGAGTACTCAACCAAGTCATTCTGAGAATAGTGTATGCGGCGACCGAGTTGCTCTTGCCCGGCGTCAATACGGGATAATACCGCGCCACATAGCAGAACTTTAAAAGTGCTCATCATTGGAAAACGTTCTTCGGGGCGAAAACTCTCAAGGATCTTACCGCTGTTGAGATCCAGTTCGATGTAACCCACTCGTGCACCCAACTGATCTTCAGCATCTTTTACTTTCACCAGCGTTTCTGGGTGAGCAAAAACAGGAAGGCAAAATGCCGCAAAAAAGGGAATAAGGGCGACACGGAAATGTTGAATACTCATACTCTTCCTTTTTCAATATTATTGAAGCATTTATCAGGGTTATTGTCTCATGAGCGGATACATATTTGAATGTATTTAGAAAAATAAACAAATAGGGGTTCCGCGCACATTTCCCCGAAAAGTGCCACCTGACGTC

**Nucleic acid Sequence of the construct PA-CXCR4-TEVcs-Elf3**

GACGGATCGGGAGATCTCCCGATCCCCTATGGTGCACTCTCAGTACAATCTGCTCTGATGCCGCATAGTTAAGCCAGTATCTGCTCCCTGCTTGTGTGTTGGAGGTCGCTGAGTAGTGCGCGAGCAAAATTTAAGCTACAACAAGGCAAGGCTTGACCGACAATTGCATGAAGAATCTGCTTAGGGTTAGGCGTTTTGCGCTGCTTCGCGATGTACGGGCCAGATATACGCGTTGACATTGATTATTGACTAGTTATTAATAGTAATCAATTACGGGGTCATTAGTTCATAGCCCATATATGGAGTTCCGCGTTACATAACTTACGGTAAATGGCCCGCCTGGCTGACCGCCCAACGACCCCCGCCCATTGACGTCAATAATGACGTATGTTCCCATAGTAACGCCAATAGGGACTTTCCATTGACGTCAATGGGTGGAGTATTTACGGTAAACTGCCCACTTGGCAGTACATCAAGTGTATCATATGCCAAGTACGCCCCCTATTGACGTCAATGACGGTAAATGGCCCGCCTGGCATTATGCCCAGTACATGACCTTATGGGACTTTCCTACTTGGCAGTACATCTACGTATTAGTCATCGCTATTACCATGGTGATGCGGTTTTGGCAGTACATCAATGGGCGTGGATAGCGGTTTGACTCACGGGGATTTCCAAGTCTCCACCCCATTGACGTCAATGGGAGTTTGTTTTGGCACCAAAATCAACGGGACTTTCCAAAATGTCGTAACAACTCCGCCCCATTGACGCAAATGGGCGGTAGGCGTGTACGGTGGGAGGTCTATATAAGCAGAGCTCTCTGGCTAACTAGAGAACCCACTGCTTACTGGCTTATCGAAATTAATACGACTCACTATAGGGAGACCCAAGCTGGCTAGCGTTTAAACTTAAGCTTGGTACCGAGCTCGGATCCACTAGTCCAGTGTGGTGGAATTCTGCAGATATCCAGCACAGTGGCGGCCGCGCCACCCGCTTTCCTAATCTGCTGGCTGCCATATGCTGGTGTGGCGTTCTACATCTTCACCCATCAGGGCTCTGACTTTGGGCCCATCTTCATGACCATCCCGGCTTTCTTTGCCAAGACGTCTGCCGTCTACAACCCGGTCATCTACATCATGATGAACAAGCAGTTCCGGACCTCTGCCCAGCACGCACTCACCTCTGTGAGCAGAGGGTCCAGCCTCAAGATCCTCTCCAAAGGAAAGCGAGGTGGACATTCATCTGTTTCCACTGAGTCTGAGTCTTCAAGTTTTCACTCCAGCATCGATACCGGTGGACGCACCCCACCCAGCCTGGGTCCCCAAGATGAGTCCTGCACCACCGCCAGCTCCTCCCTGGCCAAGGACACTTCATCGACCGGTGAGAACCTGTACTTCCAGCTAATGGCTGCAACCTGTGAGATTAGCAACATTTTTAGCAACTACTTCAGTGCGATGTACAGCTCGGAGGACTCCACCCTGGCCTCTGTTCCCCCTGCTGCCACCTTTGGGGCCGATGACTTGGTACTGACCCTGAGCAACCCCCAGATGTCATTGGAGGGTACAGAGAAGGCCAGCTGGTTGGGGGAACAGCCCCAGTTCTGGTCGAAGACGCAGGTTCTGGACTGGATCAGCTACCAAGTGGAGAAGAACAAGTACGACGCAAGCGCCATTGACTTCTCACGATGTGACATGGATGGCGCCACCCTCTGCAATTGTGCCCTTGAGGAGCTGCGTCTGGTCTTTGGGCCTCTGGGGGACCAACTCCATGCCCAGCTGCGAGACCTCACTTCCAGCTCTTCTGATGAGCTCAGTTGGATCATTGAGCTGCTGGAGAAGGATGGCATGGCCTTCCAGGAGGCCCTAGACCCAGGGCCCTTTGACCAGGGCAGCCCCTTTGCCCAGGAGCTGCTGGACGACGGTCAGCAAGCCAGCCCCTACCACCCCGGCAGCTGTGGCGCAGGAGCCCCCTCCCCTGGCAGCTCTGACGTCTCCACCGCAGGGACTGGTGCTTCTCGGAGCTCCCACTCCTCAGACTCCGGTGGAAGTGACGTGGACCTGGATCCCACTGATGGCAAGCTCTTCCCCAGCGATGGTTTTCGTGACTGCAAGAAGGGGGATCCCAAGCACGGGAAGCGGAAACGAGGCCGGCCCCGAAAGCTGAGCAAAGAGTACTGGGACTGTCTCGAGGGCAAGAAGAGCAAGCACGCGCCCAGAGGCACCCACCTGTGGGAGTTCATCCGGGACATCCTCATCCACCCGGAGCTCAACGAGGGCCTCATGAAGTGGGAGAATCGGCATGAAGGCGTCTTCAAGTTCCTGCGCTCCGAGGCTGTGGCCCAACTATGGGGCCAAAAGAAAAAGAACAGCAACATGACCTACGAGAAGCTGAGCCGGGCCATGAGGTACTACTACAAACGGGAGATCCTGGAACGGGTGGATGGCCGGCGACTCGTCTACAAGTTTGGCAAAAACTCAAGCGGCTGGAAGGAGGAAGAGGTTCTCCAGAGTCGGAACTAGTCTAGAGGGCCCGTTTAAACCCGCTGATCAGCCTCGACTGTGCCTTCTAGTTGCCAGCCATCTGTTGTTTGCCCCTCCCCCGTGCCTTCCTTGACCCTGGAAGGTGCCACTCCCACTGTCCTTTCCTAATAAAATGAGGAAATTGCATCGCATTGTCTGAGTAGGTGTCATTCTATTCTGGGGGGTGGGGTGGGGCAGGACAGCAAGGGGGAGGATTGGGAAGACAATAGCAGGCATGCTGGGGATGCGGTGGGCTCTATGGCTTCTGAGGCGGAAAGAACCAGCTGGGGCTCTAGGGGGTATCCCCACGCGCCCTGTAGCGGCGCATTAAGCGCGGCGGGTGTGGTGGTTACGCGCAGCGTGACCGCTACACTTGCCAGCGCCCTAGCGCCCGCTCCTTTCGCTTTCTTCCCTTCCTTTCTCGCCACGTTCGCCGGCTTTCCCCGTCAAGCTCTAAATCGGGGGCTCCCTTTAGGGTTCCGATTTAGTGCTTTACGGCACCTCGACCCCAAAAAACTTGATTAGGGTGATGGTTCACGTAGTGGGCCATCGCCCTGATAGACGGTTTTTCGCCCTTTGACGTTGGAGTCCACGTTCTTTAATAGTGGACTCTTGTTCCAAACTGGAACAACACTCAACCCTATCTCGGTCTATTCTTTTGATTTATAAGGGATTTTGCCGATTTCGGCCTATTGGTTAAAAAATGAGCTGATTTAACAAAAATTTAACGCGAATTAATTCTGTGGAATGTGTGTCAGTTAGGGTGTGGAAAGTCCCCAGGCTCCCCAGCAGGCAGAAGTATGCAAAGCATGCATCTCAATTAGTCAGCAACCAGGTGTGGAAAGTCCCCAGGCTCCCCAGCAGGCAGAAGTATGCAAAGCATGCATCTCAATTAGTCAGCAACCATAGTCCCGCCCCTAACTCCGCCCATCCCGCCCCTAACTCCGCCCAGTTCCGCCCATTCTCCGCCCCATGGCTGACTAATTTTTTTTATTTATGCAGAGGCCGAGGCCGCCTCTGCCTCTGAGCTATTCCAGAAGTAGTGAGGAGGCTTTTTTGGAGGCCTAGGCTTTTGCAAAAAGCTCCCGGGAGCTTGTATATCCATTTTCGGATCTGATCAAGAGACAGGATGAGGATCGTTTCGCATGGGGGAGAAACCCGGGACCAGGGTCTTCAAGAAGTCGAGCCCTAACTGCAAGCTCACCGTGTACTTGGGCAAGCGGGACTTCGTAGATCACCTGGACAAAGTGGACCCTGTAGATGGCGTGGTGCTTGTGGACCCTGACTACCTGAAGGACCGCAAAGTGTTTGTGACCCTCACCTGCGCCTTCCGCTATGGCCGTGAAGACCTGGATGTGCTGGGCTTGTCCTTCCGCAAAGACCTGTTCATCGCCACCTACCAGGCCTTCCCCCCGGTGCCCAACCCMCCCCGGCCCCCCACCCGCCTGCAGGACCGGCTGCTGAGGAAGCTGGGCCAGCATGCCCACCCCTTCTTCTTCACCATACCCCAGAATCTTCCATGCTCCGTCACACTGCAGCCAGGCCCAGAGGATACAGGAAAGGCCTGCGGCGTAGACTTTGAGATTCGAGCCTTCTGTGCTAAATCACTAGAAGAGAAAAGCCACAAAAGGAACTCTGTGCGGCTGGTGATCCGAAAGGTGCAGTTCGCCCCGGAGAAACCCGGCCCCCAGCCTTCAGCCGAAACCACACGCCACTTCCTCATGTCTGACCGGTCCCTGCACCTCGAGGCTTCCCTGGACAAGGAGCTGTACTACCATGGGGAGCCCCTCAATGTAAATGTCCACGTCACCAACAACTCCACCAAGACCGTCAAGAAGATCAAAGTCTCTGTGAGACAGTACGCCGACATCTGCCTCTTCAGCACCGCCCAGTACAAGTGTCCTGTGGCTCAACTCGAACAAGATGACCAGGTATCTCCCAGCTCCACATTCTGTAAGGTGTACACCATAACCCCACTGCTCAGCGACAACCGGGAGAAGCGGGGTCTCGCCCTGGATGGGAAACTCAAGCACGAGGACACCAACCTGGCTTCCAGCACCATCGTGAAGGAGGGTGCCAACAAGGAGGTGCTGGGAATCCTGGTGTCCTACAGGGTCAAGGTGAAGCTGGTGGTGTCTCGAGGCGGGGATGTCTCTGTGGAGCTGCCTTTTGTTCTTATGCACCCCAAGCCCCACGACCACATCCCCCTCCCCAGACCCCAGTCAGCCGCTCCGGAGACAGATGTCCCTGTGGACACCAACCTCATTGAATTTGATACCAACTATGCCACAGATGATGACATTGTGTTTGAGGACTTTGCCCGGCTTCGGCTGAAGGGGATGAAGGATGACGACTATGATGATCAACTCTGCGGATCCGGTGAGAGTCTCTTCAAAGGGCCGCGCGATTATAACCCCATCTCATCCACAATTTGTCACCTCACAAATGAGTCCGATGGACATACAACAAGCCTTTACGGCATTGGCTTCGGTCCATTCATCATCACCAACAAGCATCTTTTCAGACGCAATAACGGCACCTTGCTGGTGCAGTCCCTCCACGGCGTCTTTAAAGTCAAGAACACCACGACCCTTCAGCAACACCTGATCGACGGAAGGGATATGATTATCATTAGAATGCCCAAAGATTTCCCGCCATTTCCTCAGAAACTCAAGTTTAGAGAGCCACAGAGGGAAGAAAGGATCTGCTTGGTGACAACAAATTTTCAAACTAAGTCTATGTCATCCATGGTATCAGATACCAGCTGTACGTTTCCTAGTTCAGACGGAATATTCTGGAAGCATTGGATACAGACAAAGGACGGCCAATGCGGTAGTCCACTGGTGTCTACTCGGGATGGTTTCATCGTCGGCATTCATAGCGCCAGCAACTTCACCAATACTAATAACTACTTCACCAGCGTGCCCAAGAATTTCATGGAGCTTCTGACAAACCAGGAAGCCCAGCAGTGGGTTTCTGGATGGCGCTTGAACGCTGACTCCGTTCTGTGGGGAGGCCATAAAGTGTTTATGAGTAAACCTGAAGAACCCTTTCAGCCGGTGAAAGAGGCCACTCAGCTTATGAATGAGCTGGTTTATAGCCAGGCTACTAACTTCAGCCTGCTGAAGCAGGCTGGCGACGTGGAGGAGAACCCTGGACCTATGATTGAACAAGATGGATTGCACGCAGGTTCTCCGGCCGCTTGGGTGGAGAGGCTATTCGGCTATGACTGGGCACAACAGACAATCGGCTGCTCTGATGCCGCCGTGTTCCGGCTGTCAGCGCAGGGGCGCCCGGTTCTTTTTGTCAAGACCGACCTGTCCGGTGCCCTGAATGAACTGCAGGACGAGGCAGCGCGGCTATCGTGGCTGGCCACGACGGGCGTTCCTTGCGCAGCTGTGCTCGACGTTGTCACTGAAGCGGGAAGGGACTGGCTGCTATTGGGCGAAGTGCCGGGGCAGGATCTCCTGTCATCTCACCTTGCTCCTGCCGAGAAAGTATCCATCATGGCTGATGCAATGCGGCGGCTGCATACGCTTGATCCGGCTACCTGCCCATTCGACCACCAAGCGAAACATCGCATCGAGCGAGCACGTACTCGGATGGAAGCCGGTCTTGTCGATCAGGATGATCTGGACGAAGAGCATCAGGGGCTCGCGCCAGCCGAACTGTTCGCCAGGCTCAAGGCGCGCATGCCCGACGGCGAGGATCTCGTCGTGACCCATGGCGATGCCTGCTTGCCGAATATCATGGTGGAAAATGGCCGCTTTTCTGGATTCATCGACTGTGGCCGGCTGGGTGTGGCGGACCGCTATCAGGACATAGCGTTGGCTACCCGTGATATTGCTGAAGAGCTTGGCGGCGAATGGGCTGACCGCTTCCTCGTGCTTTACGGTATCGCCGCTCCCGATTCGCAGCGCATCGCCTTCTATCGCCTTCTTGACGAGTTCTTCTGAGCGGGACTCTGGGGTTCGAAATGACCGACCAAGCGACGCCCAACCTGCCATCACGAGATTTCGATTCCACCGCCGCCTTCTATGAAAGGTTGGGCTTCGGAATCGTTTTCCGGGACGCCGGCTGGATGATCCTCCAGCGCGGGGATCTCATGCTGGAGTTCTTCGCCCACCCCAACTTGTTTATTGCAGCTTATAATGGTTACAAATAAAGCAATAGCATCACAAATTTCACAAATAAAGCATTTTTTTCACTGCATTCTAGTTGTGGTTTGTCCAAACTCATCAATGTATCTTATCATGTCTGTATACCGTCGACCTCTAGCTAGAGCTTGGCGTAATCATGGTCATAGCTGTTTCCTGTGTGAAATTGTTATCCGCTCACAATTCCACACAACATACGAGCCGGAAGCATAAAGTGTAAAGCCTGGGGTGCCTAATGAGTGAGCTAACTCACATTAATTGCGTTGCGCTCACTGCCCGCTTTCCAGTCGGGAAACCTGTCGTGCCAGCTGCATTAATGAATCGGCCAACGCGCGGGGAGAGGCGGTTTGCGTATTGGGCGCTCTTCCGCTTCCTCGCTCACTGACTCGCTGCGCTCGGTCGTTCGGCTGCGGCGAGCGGTATCAGCTCACTCAAAGGCGGTAATACGGTTATCCACAGAATCAGGGGATAACGCAGGAAAGAACATGTGAGCAAAAGGCCAGCAAAAGGCCAGGAACCGTAAAAAGGCCGCGTTGCTGGCGTTTTTCCATAGGCTCCGCCCCCCTGACGAGCATCACAAAAATCGACGCTCAAGTCAGAGGTGGCGAAACCCGACAGGACTATAAAGATACCAGGCGTTTCCCCCTGGAAGCTCCCTCGTGCGCTCTCCTGTTCCGACCCTGCCGCTTACCGGATACCTGTCCGCCTTTCTCCCTTCGGGAAGCGTGGCGCTTTCTCATAGCTCACGCTGTAGGTATCTCAGTTCGGTGTAGGTCGTTCGCTCCAAGCTGGGCTGTGTGCACGAACCCCCCGTTCAGCCCGACCGCTGCGCCTTATCCGGTAACTATCGTCTTGAGTCCAACCCGGTAAGACACGACTTATCGCCACTGGCAGCAGCCACTGGTAACAGGATTAGCAGAGCGAGGTATGTAGGCGGTGCTACAGAGTTCTTGAAGTGGTGGCCTAACTACGGCTACACTAGAAGAACAGTATTTGGTATCTGCGCTCTGCTGAAGCCAGTTACCTTCGGAAAAAGAGTTGGTAGCTCTTGATCCGGCAAACAAACCACCGCTGGTAGCGGTGGTTTTTTTGTTTGCAAGCAGCAGATTACGCGCAGAAAAAAAGGATCTCAAGAAGATCCTTTGATCTTTTCTACGGGGTCTGACGCTCAGTGGAACGAAAACTCACGTTAAGGGATTTTGGTCATGAGATTATCAAAAAGGATCTTCACCTAGATCCTTTTAAATTAAAAATGAAGTTTTAAATCAATCTAAAGTATATATGAGTAAACTTGGTCTGACAGTTACCAATGCTTAATCAGTGAGGCACCTATCTCAGCGATCTGTCTATTTCGTTCATCCATAGTTGCCTGACTCCCCGTCGTGTAGATAACTACGATACGGGAGGGCTTACCATCTGGCCCCAGTGCTGCAATGATACCGCGAGACCCACGCTCACCGGCTCCAGATTTATCAGCAATAAACCAGCCAGCCGGAAGGGCCGAGCGCAGAAGTGGTCCTGCAACTTTATCCGCCTCCATCCAGTCTATTAATTGTTGCCGGGAAGCTAGAGTAAGTAGTTCGCCAGTTAATAGTTTGCGCAACGTTGTTGCCATTGCTACAGGCATCGTGGTGTCACGCTCGTCGTTTGGTATGGCTTCATTCAGCTCCGGTTCCCAACGATCAAGGCGAGTTACATGATCCCCCATGTTGTGCAAAAAAGCGGTTAGCTCCTTCGGTCCTCCGATCGTTGTCAGAAGTAAGTTGGCCGCAGTGTTATCACTCATGGTTATGGCAGCACTGCATAATTCTCTTACTGTCATGCCATCCGTAAGATGCTTTTCTGTGACTGGTGAGTACTCAACCAAGTCATTCTGAGAATAGTGTATGCGGCGACCGAGTTGCTCTTGCCCGGCGTCAATACGGGATAATACCGCGCCACATAGCAGAACTTTAAAAGTGCTCATCATTGGAAAACGTTCTTCGGGGCGAAAACTCTCAAGGATCTTACCGCTGTTGAGATCCAGTTCGATGTAACCCACTCGTGCACCCAACTGATCTTCAGCATCTTTTACTTTCACCAGCGTTTCTGGGTGAGCAAAAACAGGAAGGCAAAATGCCGCAAAAAAGGGAATAAGGGCGACACGGAAATGTTGAATACTCATACTCTTCCTTTTTCAATATTATTGAAGCATTTATCAGGGTTATTGTCTCATGAGCGGATACATATTTGAATGTATTTAGAAAAATAAACAAATAGGGGTTCCGCGCACATTTCCCCGAAAAGTGCCACCTGACGTC

**Photoactivation**

Photoactivation with blue light at an intensity of 3.2 W/m2.

Through collaborative efforts with Radiometech, we developed the MAGI-01 device along with its LED illumination platform. This was achieved utilizing expertise of one of our team members Joanna Kałafut in the field of biophysics, who played an essential role in the technical design and configuration of parameters enabling precise selection of the time and intensity of irradiation of cells with blue light.

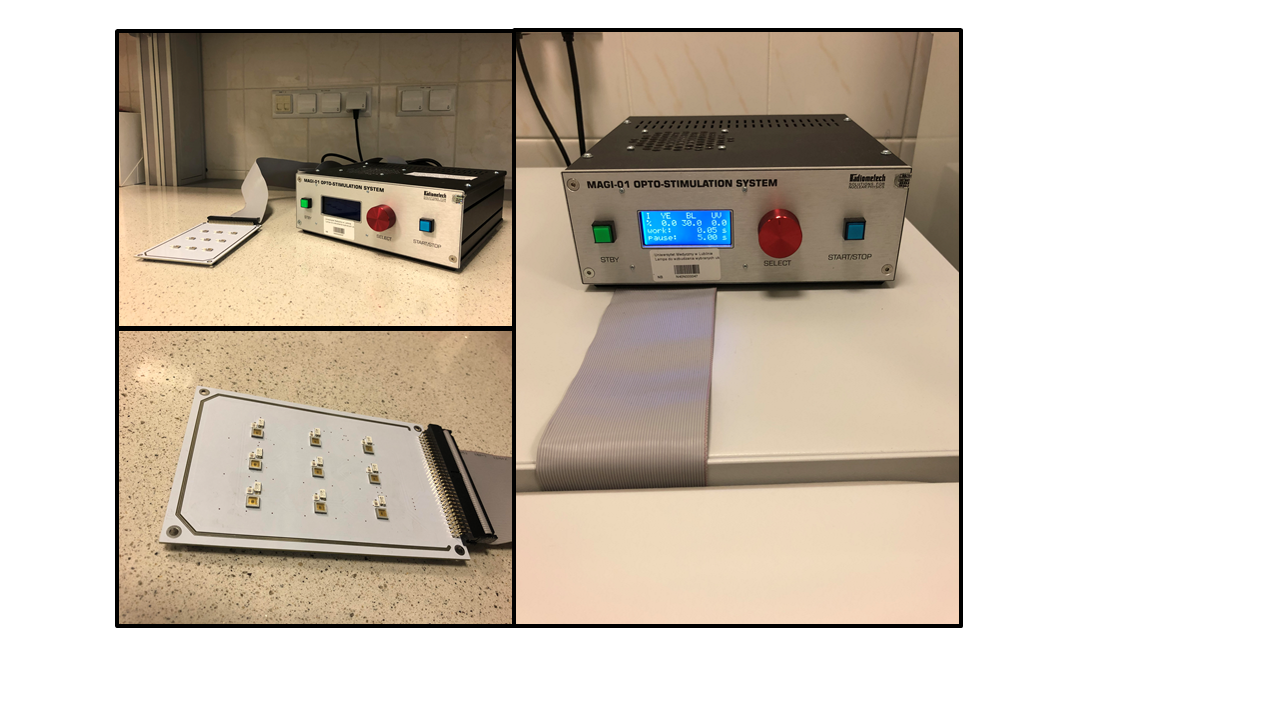

**Figure 1.** The MAGI-01 device and the illumination (LED) platform. The control panel and the diode illumination platform (left), which has been designed to fit in a cell culture incubator (right image). Image reproduced from Kałafut et al.(Kałafut et al., 2022) (CC BY).

**EMT induction**

Cells were culturing in the presence of medium with TGFβ, added 12h after transfection.

**RT-qPCR Primers**

| Gene | Primer sequences | |
| --- | --- | --- |
| *GAPDH* | Forward | 5′- CTCTGCTCCTCCTGTTCGAC -3’ |
|  | Reverse | 5′- GCCCAATACGACCAAATCC -3’ |
| *NOTCH1* | Forward | 5′- GCAGATGGCTCAACGGCACTG-3’ |
|  | Reverse | 5′- GGGGTCTCCTCCTTGCTATCCTG-3’ |
| *JAG1* | Forward | 5′- GCCGAGGTCCTATACGTTGC-3’ |
|  | Reverse | 5′- CCGAGTGAGAAGCCTTTTCAA-3’ |
| *HES1* | Forward | 5′- TCAACACGACACCGGATAAAC -3’ |
|  | Reverse | 5’- GCCGCGAGCTATCTTTCTTCA -3’ |
| *HEY1* | Forward | 5′- CGGCTCTAGGTTCCATGTCC -3’ |
|  | Reverse | 5’- GCTTAGCAGATCCCTGCTTCT -3’ |
| *MMP9* | Forward | 5′- GCTGACTACGATAAGGACGGCA-3’ |
|  | Reverse | 5′- TAGTGGTGCAGGCAGAGTAGGA-3’ |
| *NANOG* | Forward | 5′- TGAGTGTGGATCCAGCTTGT-3’ |
|  | Reverse | 5′- CCATGGAGGAAGGAAGAGGAGAGA-3’ |
| *SLUG* | Forward | 5′- ATCTGCGGCAAGGCGTTTTCCA-3’ |
|  | Reverse | 5′- GAGCCCTCAGATTTGACCTGTC-3’ |
| *SNAIL* | Forward | 5′- TCGGAAGCCTAACTACAGCGA-3’ |
|  | Reverse | 5′- AGATGAGCATTGGCAGCGAG-3’ |
| *STAT3* | Forward | 5′- CTTTGAGACCGAGGTGTATCACC-3’ |
|  | Reverse | 5′- GGTCAGCATGTTGTACCACAGG-3’ |
| *VIM* | Forward | 5′- AGTCCACTGAGTACCGGAGAC-3’ |
|  | Reverse | 5′- CATTTCACGCATCTGGCGTTC-3’ |
| *ZEB1* | Forward | 5′- GGCATACACCTACTCAACTACGG-3’ |
|  | Reverse | 5′- TGGGCGGTGTAGAATCAGAGTC-3’ |
| *CDH1* | Forward | 5′- ATTTTTCCCTCGACACCCGAT-3’ |
|  | Reverse | 5′- TCCCAGGCGTAGACCAAGA-3’ |
| *FIBRONECTIN1* | Forward | 5′- ACAACACCGAGGTGACTGAGAC-3’ |
|  | Reverse | 5′- GGACACAACGATGCTTCCTGAG-3’ |

**Activity of GLuc**

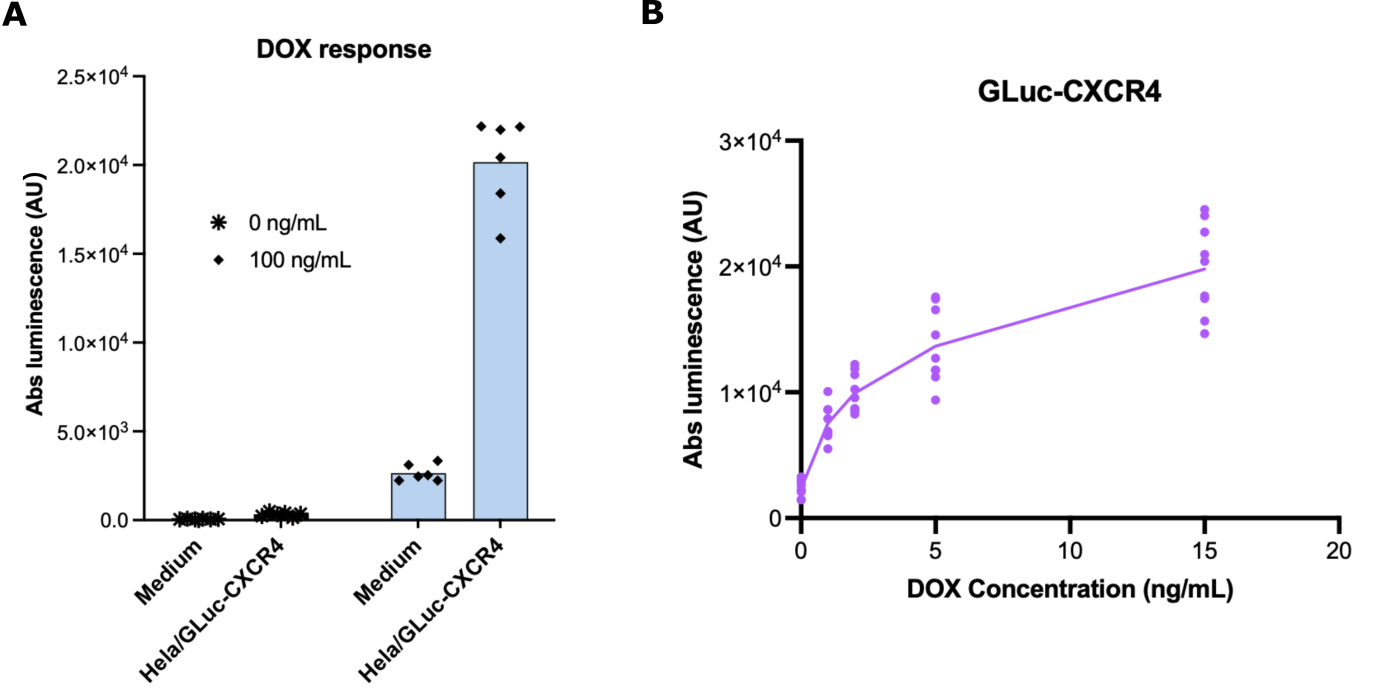

**Figure 2. (A)** To assess if the receptor resides only on the plasma membrane and is not cleaved or secreted into the medium, HeLa TET-ON cells were transfected with the Gluc-CXCR4 plasmid and cultured in the presence of 100 ng/mL doxycycline after 24 hours of transfection or in the absence doxycycline as control. **(B)** Dose depending Gluc-CXCR4 luminescence activity. After 24 hours of transfection of Gluc-CXCR4, Doxycycline was added to culture media at concentration of 0 – 1 – 2 – 5 – 15 ng/mL respectively to G-luciferase activity was measured in the next day using a Synergy H1 Luminometer (Biotek Instruments, Winooski, VT, USA) by injected with 50 μl of 20 mM coelenterazine and the number of photons were recorded for 1 second after injecting. Data were collected for both culture media and cells expressing Gluc-CXCR4 in triplicate (with and without Doxycycline) **(A)** and for cells expressing Gluc-CXCR4 **(B)**.
